# A gene expression atlas of a juvenile nervous system

**DOI:** 10.1101/2025.11.21.689793

**Authors:** Seth R. Taylor, Claire Olson, Lidia Ripoll-Sanchez, Giulio Valperga, Rebecca McWhirter, S. Talmage Barney, Alexander Atkinson, Sidharth Goel, Alexis Weinreb, Andrew Hardin, Alexis Rolfson, Jacob Pattee, G. Robert Aguilar, Daniel M. Merritt, Matthew Eroglu, Maryam Majeed, Ethan Grundvig, Ethan Child, Isabel Beets, Petra E. Vértes, William R. Schafer, Erdem Varol, Marc Hammarlund, Oliver Hobert, David M. Miller

## Abstract

The architecture of metazoan nervous systems is typically established in the embryo, but substantial numbers of neurons are added postnatally as existing neurons refine connectivity and differentiate. To reveal molecular determinants of postembryonic neurogenesis and maturation, we produced gene expression profiles of all neuron types and their progenitors in the first larval stage (L1) of *C. elegans*. We identified thousands of differentially expressed genes across individual neurons between the L1 and earlier embryonic and later L4 larval stage. Key neuropeptide signaling networks are remodeled during larval development. Gene regulatory network analysis revealed potential transcription factors driving temporal changes in gene expression. We utilized connectomic data along with our neuron-specific atlas to identify potential molecular determinants of membrane contact and synaptic connectivity. These expression data are available through a user-friendly interface at CeNGEN.org for independent investigations of the maturation, connectivity and function of a developing nervous system.

## INTRODUCTION

Neural circuits are generated by defined patterns of cell division giving rise to post-mitotic neurons that adopt distinctive morphologies and synaptic connections. In mammals, most neurons are generated during embryonic development^1,2^. Postnatal neurons may also differentiate during development to modify circuit architecture and physiology. Gene expression, regulated within neuron type and across development, orchestrates the overall neurogenic program and thus embodies the genetic blueprint for the brain.

The *C. elegans* nervous system provides an attractive model for studies of neurogenic pathways. Over two-thirds of the neurons (222/302) in the adult hermaphrodite nervous system are generated in the embryo^3,4^. Most additional neurons are added during the first (L1) larval stage^4^ from a defined set of progenitors and undergo post-mitotic differentiation as they are integrated with existing circuits. The differentiated state of embryonically derived neurons is also modified during larval development resulting in cell volume expansion and circuit remodeling^5–11^. These anatomical changes are accompanied by behavioral differences that likely depend on dynamic gene expression during larval maturation^12–15^.

Here we describe single cell RNA-seq (scRNA-seq) profiles of all *C. elegans* neurons in the L1 larval stage to complement our previous late larval atlas and other partial early larval atlases^16–19^. Importantly, these data encompass the entire L1 nervous system. We captured neuronal precursor lineages in the L1 including all progenitors of postembryonic ventral cord motor neurons and analyzed developmentally expressed genes for neurogenesis and differentiation across the entire nervous system. Our approach identified thousands of differentially expressed genes across development and gene regulatory network analysis identified transcription factors (TFs) that may orchestrate these effects. We determined that L1 stage neuropeptide signaling networks are remodeled as larval development progresses and new neuropeptide-dependent behaviors emerge^12,20^. We compared gene expression profiles with connectomic and protein-protein interaction data to identify cell adhesion molecules associated with specific neuronal circuits. Our data provide a resource for investigating the maturation of individual neurons and their progenitors for an entire nervous system.

## RESULTS AND DISCUSSION

### A comprehensive gene expression profile of L1-stage neurons

We used 10X Genomics single-cell RNA sequencing (scRNA-Seq) to profile the mid-late L1 nervous system (Figure 1A-C, Figure S1A-B, Tables S1, S2)^21^. After quality control (see Methods), we generated profiles of 151,415 cells from 16 experiments (Figure S1C, Table S3) with medians of 841 unique molecule identifiers (UMIs) 373 genes per cell. Using previously published datasets as references^16,17,22,23^, we assigned anatomical identities to 93.5% of the cells. In addition to neurons, we annotated cells belonging to postembryonic progenitor lineages and terminally differentiated non-neuronal cell classes (Figure S1C-E, Table S4). Altogether, we identified 89,073 post-mitotic neurons with medians of 701 UMIs and 326 genes per cell (Figure 1B).

**Figure 1.**
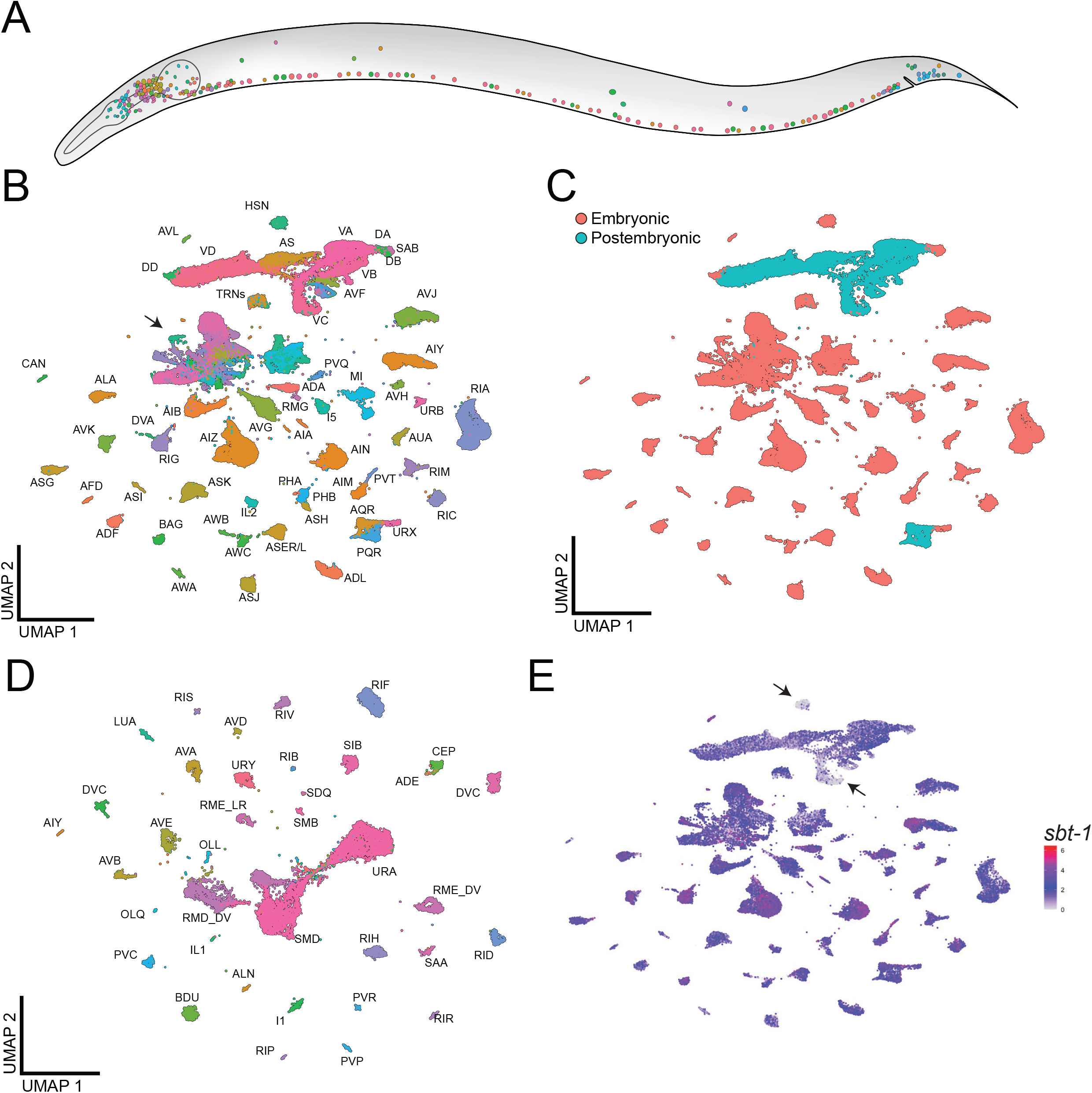
scRNA-Seq profiles of the L1 hermaphrodite nervous system. A) Colors denote anatomically defined neuron classes. B-C) UMAP projections of 89,073 L1 neurons, colored by class (B) or neuron generation time (C). Arrow in B denotes cell groups in sub-UMAP in panel D. D) Sub-UMAP of cluster panel B (arrow). E) UMAP as in (B) colored by expression of the post-mitotic neuron marker *sbt-1*. Arrows: HSN and VC, which show low *sbt-1* expression. See also Figures S1-S3, Tables S1-S4.

The 302 neurons of the *C. elegans* adult hermaphrodite comprise 118 distinct classes based on morphology and connectivity^24,25^. Ninety-six classes are generated embryonically^3^, six classes are generated in mid-L1, 11 classes in late L1, and five classes are generated or differentiate in the L2 or later^4^. Our L1 dataset contains transcriptomes of all hermaphrodite neuron classes at mid L1 (Figure 1B-E), including the immature HSN neurons (see Methods, Figure S2A-D), and most classes born in late L1.

Several anatomically-defined neuron classes have been further separated into molecularly distinct subclasses in the L4 and adult nervous system^16,25,26^. Consistent with previous work in the embryo^22,27,28^, we detected ASE and AWC subclasses in the L1 larva. We found that IL2, RME and RMD subclasses are distinguishable by the L1 stage (Figure S3A)^16,23,26^. We detected two subclasses for the DA and DB motor neurons. The most anterior neurons of each class, DA1 and DB2, adopted distinguishable gene expression profiles (Table S4, Figure S3B-F), consistent with recent scRNA-Seq data from adults^26^. We also detected subclasses for the postembryonic VA, VB, and VD ventral cord motor neuron classes (discussed below). In total, we identified 126 transcriptionally distinct neuron classes.

### Temporally regulated genes distinguish progenitors from post-mitotic neurons

Postembryonically derived neurons generated in the L1 derive from a limited number of progenitor lineages. We captured cell-specific profiles of progenitor cells and post-mitotic neurons from the G1, Q, P and T lineages. The K cell and DVB were not identified (see Methods). Cell populations corresponding to neuroblast lineages were identified by expression of known cell cycle genes, Hox TFs, and additional reference genes from the literature (see Methods, Table S4, Figure S4A-F). In the Q lineage (Figure 2A), we captured post-mitotic AQR and PQR neurons, the neuroblasts QL.pa/QR.pa, and AVM, PVM, and SDQ neurons. UMAP embeddings for the QL.pa/QR.pa lineages match the pattern of known cell divisions. Two branches diverge from the progenitor population, corresponding to SDQL/R and AVM/PVM (Figure 2B, Figure S4A-B). We annotated three progenitor populations and all three postmitotic neuron classes from the T.pp lineage (see Methods, Figure 2C-D, Figure S4C-H). We noted expression of the proneural TF *lin-32* in the T.pp neuroblasts (Figure S4E-F). Previous work demonstrated that *lin-32* regulates neuronal fate in several lineages^29–36^. To identify whether *lin-32* influences neuronal fate in the T-lineage, we determined that the loss of function allele *lin-32(u282)* dramatically reduced the number of VAB-7 expressing neurons in the tail (Figure S4I-J).

**Figure 2.**
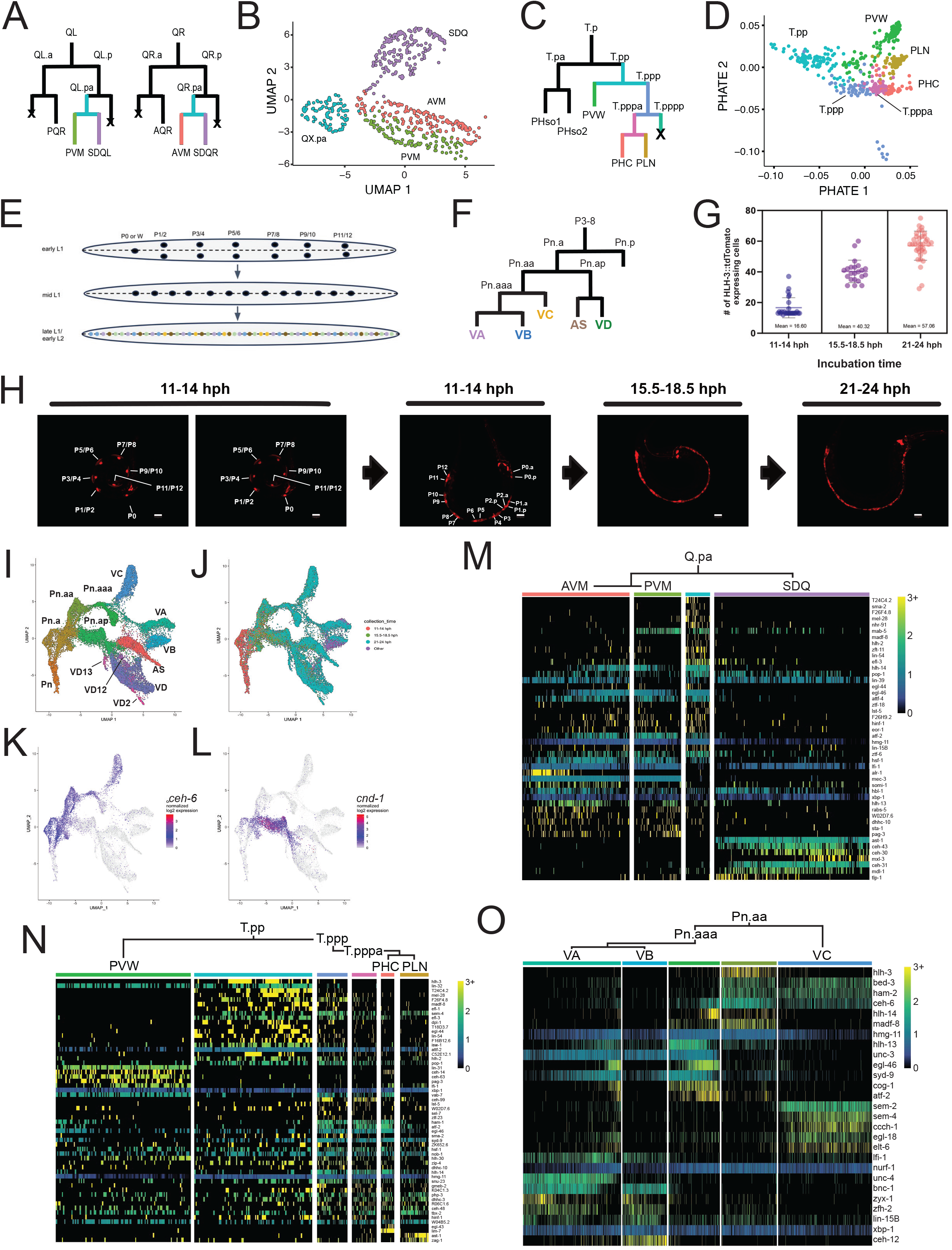
Neuronal progenitors in the Q, T, and P lineages. A) QL and QR lineage diagrams. B) Sub-UMAP of Q.pa lineage colored by neuron class. C) The T.p lineage diagram. D) PHATE plot of T.pp lineage colored by cell class. E) P-cell nuclei migration to the ventral cord and subsequent cell divisions during the L1 stage. F) P3-P8 lineage diagrams. G) *hlh-3*::tdTomato expressing cells counted during each collection period (hph, hours post hatch). H) Confocal micrographs of *hlh-3*::tdTomato expressing cells in the ventral cord for each collection period. Note bilateral array of P-cells before migration to the ventral cord, depicted with two different Z-planes in the same individual. Scale bars = 10 µm. I-L) Sub-UMAPs of P1-P12 lineages colored by cell type (I), collection time (J), expression of the TF *ceh-6* (K), and expression of the TF *cnd-1* (L). M-O) Heatmaps of scaled expression values of TFs differentially expressed between cell types in the Q lineage (M), T lineage (N) and P.aa sub-lineage (O). See also Figures S4, S5 and Tables S5, S6.

The 13 P cell progenitors (P0 – P12) produce the majority of postembryonic neurons. In newly born L1 larva, P1-P12 appear as left-right pairs of cells on either side of the body; P0 (aka “W”) is positioned anteriorly. In mid L1, P1-P12 migrate toward the ventral cord and in late L1 undergo stereotypical cell divisions (Figure 2E). P3-P8 give rise to identical lineages: the anterior daughter, Pn.a, generates VA, VB, VC, AS and VD ventral cord motor neurons, while the posterior daughter, Pn.p, generates an epidermal cell (Figure 2F). Anterior (P0-P2) and posterior (P9-P12) P cells adopt related patterns of division (Figure S5A)^4,37^. We used *hlh-3p::tdTomato* to capture P-lineage progenitors and neurons before, during and after terminal divisions (see Methods, Figure 2G-H). We identified 39,054 cells spanning the P-lineage, including 10 progenitor classes (Pn, Pn.p, Pn.a, Pn.aa, Pn.aaa, P0, P0.a, P0.p, P0.aa/P1.aaa) and 6 post-mitotic neuron classes (Figure 2I-J; Figure S4K-L; Figure S5B) (see Methods).

### P0-P1 lineages are transcriptionally distinct from P2-P12 lineages

P-lineage populations comprised two distinct groups in UMAP projections. One group mirrors known cell divisions of P2-P12 lineages, featuring VC, VA and VB neurons branching from one progenitor cluster (Pn.aaa) and VD and AS neurons branching from another (Pn.ap) (Figure 2I-L; Figure S5B). The second group of P-lineage cells reflects the P0-P1 lineages, including VB and AVF neurons branching from one progenitor cluster and VA and VD neurons branching from another (Figure S5B-C). Hox gene expression patterns confirmed our P0/P1 and P2-12 annotations (see Methods, Figure S5C-J)^26^. Adult ventral cord motor neuron classes can be further categorized into transcriptionally distinct subclasses^16,26^. In the L1 data, we identified VB1 and the VD subclasses VD2, VD12, and VD13 (Figure 2I, Figure S5K-L), but not subclusters of P2-P12 derived VA, VB, VC, or AS motor neurons previously identified (e.g., VA2, VA11, VA12; Figure S5M-N), despite similar sampling and sequencing depth (Table S5, see Methods). This finding suggests that transcriptional differences among some postembryonic motor neuron subclasses become more pronounced throughout larval development.

### Differential expression of TFs in postembryonic neuronal lineages

To identify candidate regulators of postembryonic cell fate determination, we examined differential gene expression within the Q, T and P lineages. We identified differentially expressed genes (DEGs) between every pair of parent-daughter cells and sister cells within each lineage (Table S6). To illustrate our results, we highlight TFs that are >4 fold enriched across the Q (Figure 2M) and T (Figure 2N) lineages, as well as the Pn.aa sub-lineage that gives rise to VA, VB and VC motor neurons (Figure 2O). These genes provide a useful starting point for investigating the gene regulatory mechanisms that drive postembryonic neuron differentiation.

### Determining gene expression in early post-mitotic neuron maturation

To identify gene expression changes in early post-mitotic neuronal maturation, we calculated pseudotime trajectories within the postembryonic Q, T and P progenitor lineages (Figure 3A-D) as well as for 15 embryonic progenitor and neuron classes (see Methods, Figure 3E-F). Embryonic pseudotime estimates strongly correlated with the embryo time estimates based on correlation to whole embryo RNAseq data^22^ (Figure 3G), indicating pseudotime values accurately reflect relative developmental ages. We identified 4,452 genes with significantly altered expression along pseudotime from 16 postembryonic classes and 1,342 genes from 15 embryonic classes, with a significant overlap between stages (see Methods, Figure 3H-N, Table S7). For example, the neuropeptide processing gene *pghm-1* increases with maturation in 20 neurons (9 postembryonic, 11 embryonic, Figure 3O), whereas the H1 linker histone *his-24* decreases during the differentiation of 18 neurons (6 postembryonic, 12 embryonic, Figure 3P). Several genes exhibit mixed patterns across neurons. The *hil-7* gene, which also encodes an H1 linker histone, shows differential expression in 27 neuron classes with a wide range of expression profiles (Figure 3Q).

**Figure 3.**
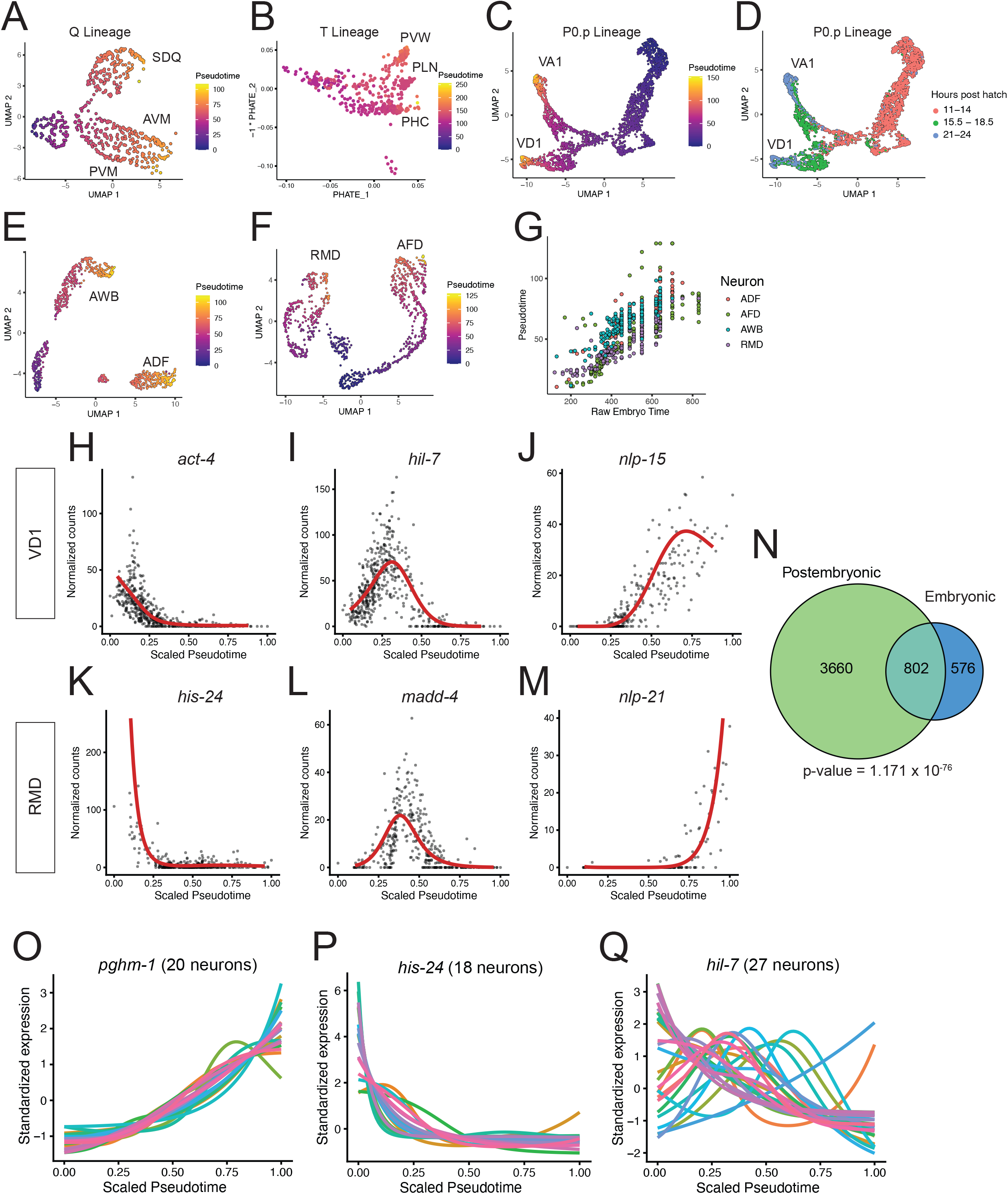
Gene expression changes during early neuronal differentiation. A) UMAP of Q.pa lineage colored by pseudotime. B) PHATE plot of T.pp lineage colored by pseudotime. C-D) UMAPs of P0.p lineage colored by pseudotime (C) and hours post hatch (D). E-F) Sub-UMAPs of embryonic progenitors and neurons colored by pseudotime for AWB/ADF (E) and RMD/AFD (F). G) Scatterplot showing raw embryo time^22^ vs pseudotime for RMD, AFD, AWB and ADF. H-M) Genes with significant expression changes over pseudotime for the postembryonic VD1 neuron (H-J) and the embryonic RMD neurons (K-M). Each dot denotes an individual cell; y-axis is normalized expression (UMI counts / Size Factor). Pseudotime values were scaled from 0-1 for each neuron class. The red line denotes a fit to gene expression. N) Venn diagram of genes exhibiting expression changes along pseudotime in postembryonic neurons (green) and embryonic neurons (blue). O-P) Expression profiles of *pghm-1* (O), *his-24* (P) and *hil-7* (Q) along scaled pseudotime for neurons with significant changes. See also Figure S6 and Table S7.

To further explore similarities between embryonic and larval neuronal progenitors and newborn neurons, we calculated pseudobulk profiles of immediate neuronal progenitors and postmitotic neurons. Pairwise Spearman correlations and hierarchical clustering revealed distinct separations between larval and embryonic populations and between progenitors and postmitotic neurons within each stage (Figure S6A). Network analysis of transcriptional similarities between progenitors and neurons (Figure S6B, see Methods) yielded largely distinct communities of postembryonic neurons, larval progenitors, embryonic progenitors, and embryonic neurons. While larval progenitors showed the highest correlations with other larval progenitors, they also showed similarities with many embryonic progenitors, further supporting shared genetic programs of neuronal differentiation in the embryonic and larval stages.

### Defining gene expression across the L1 nervous system

We performed dynamic thresholding^16,26^ of our L1 data to identify genes confidently expressed in each neuron type (see Methods, L1 CengenApp at cengen.shinyapps.io/L1app). The number of genes detected (threshold 2) per neuron class (median = 5478) was positively correlated with the number of cells captured per neuron class (median 357 cells, Figure S7A, Spearman rank correlation = 0.366; p = 2.477 × 10^−5^). 13 sparsely sampled neuron classes have the fewest detected genes and true positive rates (Figure S7B) and consequently are likely to include higher numbers of false negatives.

We catalogued expression of gene families with key roles in the nervous system, e.g., TFs, cell adhesion molecules, and neuropeptides and their receptors. To illustrate our findings, we note that most homeodomain family TFs are sparsely expressed across neuron types (Figure 4A-B), whereas other TF families (e.g. AT-Hook, bZIP) are more broadly detected (Figure 4B). Similar results were obtained at the L4 stage, indicating that these characteristic patterns of expression for TF classes (i.e., sparse vs broad) are largely established early in neuronal development.

**Figure 4.**
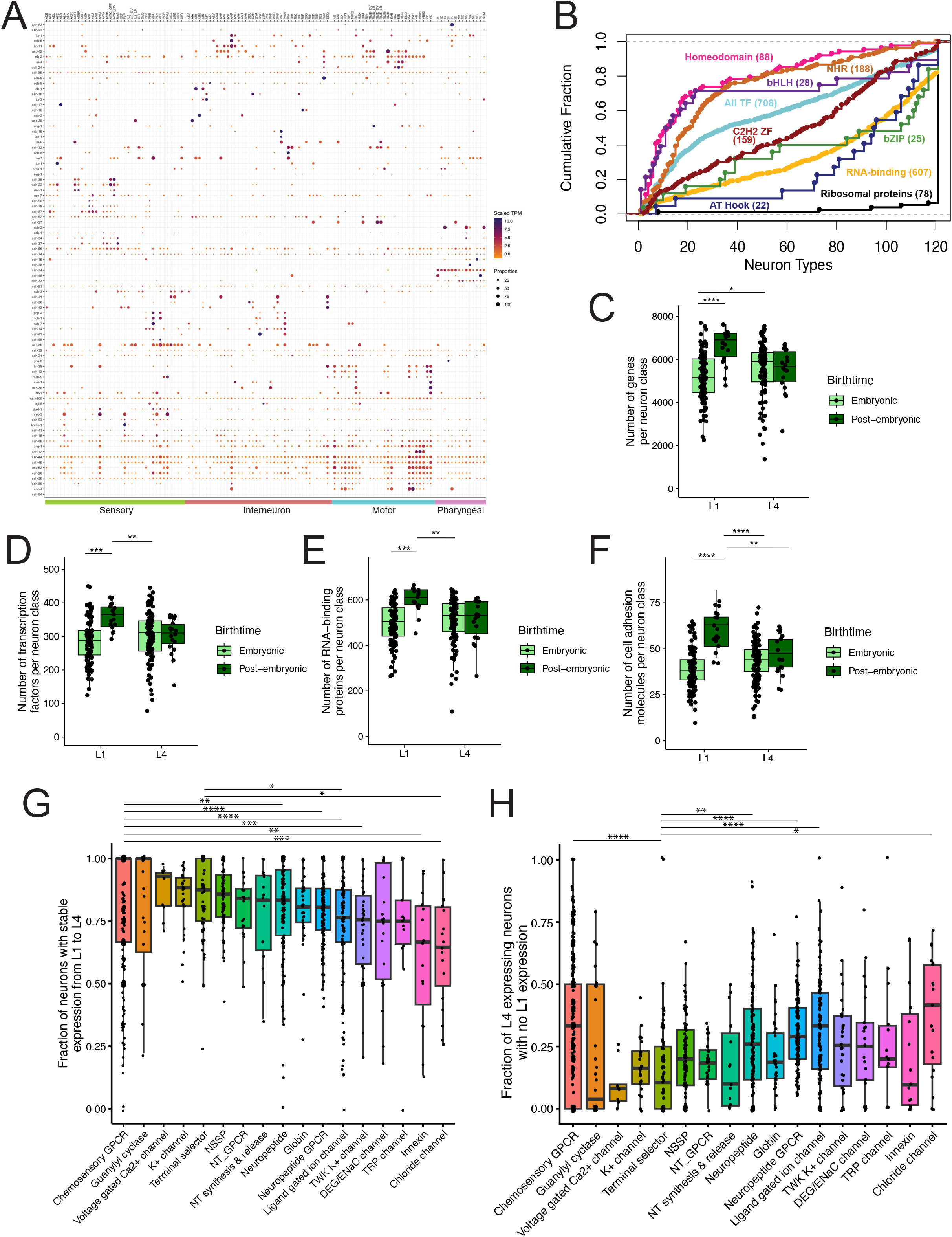
Expression of TFs, RNA-binding proteins and cell adhesion molecules in the larval nervous system. A) Clustered dot plot of L1 homeodomain TF expression across 121 neuron types grouped by functional category. Dots are colored by scaled TPM value and sized by proportion of TF-expressing cells. B) Cumulative distribution of TF families at L1, with the number of genes expressed from each family in parentheses. C) Total number of genes expressed at threshold 2 in each neuron class at the L1 (left) and L4 (right) stages. Neuron types are divided by birthtime. D-F) Quantification of gene families at threshold 2 for D) TFs, E) RNA-binding proteins, and F) cell adhesion molecules. Groups were compared using linear models with birthtime, stage, and number of cells per neuron class as covariates and Tukey p-value adjustment for multiple comparisons of estimated marginal means. G) Box and jitter plots of gene expression stability in embryonic neurons from L1 to L4 for neuronal gene families. H) Box and jitterplots of the fraction of L4-expressing neurons with no expression at L1 for neuronal gene families. For G-H), Kruskal-Wallis tests followed by Dunn’s multiple comparison test. * p < 0.05, ** p < 0.01, *** p < 0.001, **** p < 0.0001. See also Figure S7 and Tables S16, S17.

We detected more genes per neuron class in postembryonic neurons compared to embryonic neurons in the L1, but not the L4 (Figure 4C). We noted that TFs, RNA binding proteins and cell adhesion proteins were similarly more numerous in postembryonic neurons (Figure 4D-F), a finding that mirrors cadherin expression across larval development^38^. GPCR neuropeptide receptors and metabotropic receptors showed the opposite pattern, with fewer genes expressed in postembryonic neurons compared to embryonic neurons in the L1 (Figure S7C-D). Ribosomal genes and ligand-gated ion channels showed no differences (Figure S7E-F). These results point to gene families upregulated in newborn L1 neurons to promote post-mitotic differentiation versus earlier generated embryonic neurons. Conversely, fewer GPCRs and metabotropic receptors in postembryonic neurons in the L1 could indicate immature signaling function.

### Stability of gene expression between the L1 and L4 nervous systems

Nervous system development involves conserved core features, such as neurotransmitter usage and characteristic anatomy. We therefore investigated whether L1 stage expression of gene families with neuronal function was preserved in the L4 (see Methods). Expression of chemosensory GPCRs, guanylyl cyclases, voltage-gated Ca2+ channels, K+ channels and terminal selector TFs was highly stable from L1 to L4 (Figure 4G). We also measured the appearance of new expression at the L4 stage. Terminal selector TFs, guanylyl cyclases, voltage-gated Ca2+ channels, neurotransmitter-related genes and innexins had relatively few instances of newly gained expression from L1 to L4. Chemosensory GPCRs, neuropeptides, neuropeptide receptors, ligand-gated ion channels, and chloride channels frequently demonstrated newly gained expression (Figure 4H). These data indicate that as neurons mature, early chemosensory GPCR expression patterns remain largely stable while additional patterns emerge. Neuropeptides, neuropeptide receptors and ligand-gated ion channels feature relatively higher rates of both loss of early expression and gain of new expression. Overall, these data demonstrate the stability of several families of neuronal genes, including terminal selectors and classical neurotransmitter systems, in the embryonically born nervous system across larval development.

### Differential gene expression across specific neuron types during development

*C. elegans* larval development features changes in anatomy^4,6,7,8,39,40^, connectivity^9–11,41^, locomotor behavior^12,13^ and odor preferences^14,15^. We therefore performed differential gene expression analysis between the embryo and the L1 for 89 neuron classes and between the L1 and L4 stages for 121 neuron classes (See Tables S8-S9). Between the embryo and L1, we detected 54,757 instances of differential expression (i.e., a gene showing differential expression between ages in a given neuron) featuring 5941 DEGs (Figure 5A, Table S10). Between L1 and L4, we detected 36,350 instances of differential expression featuring 5810 DEGs (Figure 5B, Table S11).

**Figure 5.**
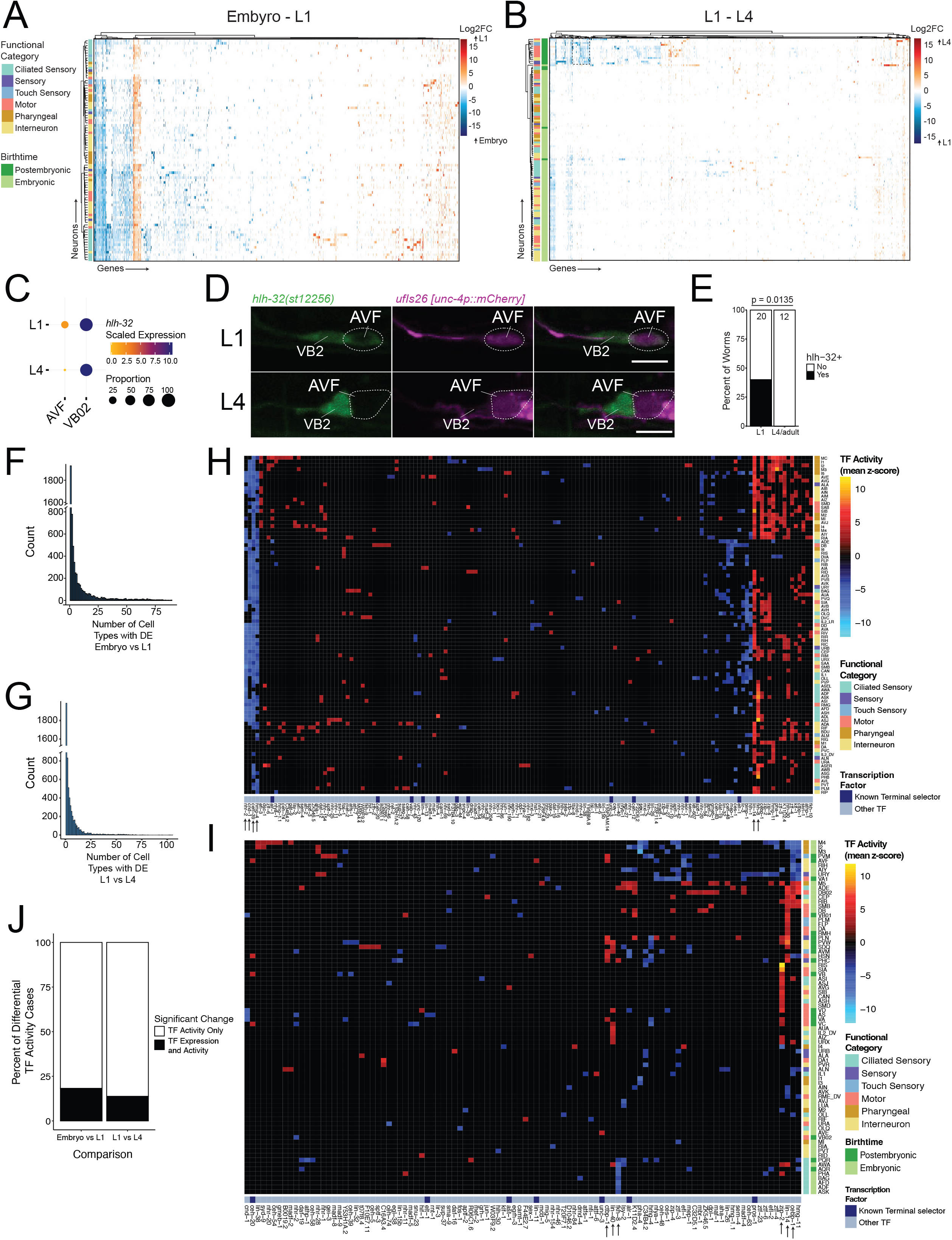
Thousands of genes are differentially expressed across larval development. A-B) Clustered heatmaps of log2 fold change (log2FC) for DEGs between A) embryonic and L1 stages and B) L1 and L4 stages. Positive log2FC values (orange/red) reflect higher gene expression in the older stage; negative log2FC values (blue) reflect higher expression in the younger stage. Neuronal functional categories and birthtimes are depicted as left Y-axis labels. C) Dotplot of scRNA-seq data for *hlh-32* in AVF and VB2 in L1 and L4. D) Confocal micrographs showing endogenously GFP-tagged *hlh-32* in AVF in L1, but not L4 vs stable expression in VB2. Scale bar = 5 µm. E) Quantification of *hlh-32*::GFP expression in AVF in L1 and L4/adults, Fisher’s Exact Test. F-G) Histograms of neuron classes in which a gene exhibited significant differential expression for the embryo vs L1 (F) and the L1 vs L4 (G) comparisons. H-I) Heatmaps showing TFs (columns) with significant activity changes across neurons (rows) between the H) embryonic and L1 stages and I) the L1 and L4 stages. Colors indicate directionality of change in TF regulon expression (red/yellow denotes increased expression of a TF regulon in the older stage, blue denotes decreased expression in the older stage). Functional categories are color coded (right), and TFs with known roles as terminal selectors (in any neuron) are denoted in dark blue (bottom). Black arrows (bottom) mark TFs with significant changes across many neurons. J) Fraction of instances of differential TF activity which also showed differential TF expression (black) or which lacked significant differential TF expression (white). See also Figures S8-S12, Tables S8-S15.

We found significant concurrence with genes previously reported to differ between L1 and L4^12^, particularly among genes with higher expression at L4 (Figure S8A). Discrepancies could be due to disparate timing for L1 imaging and scRNA-seq experiments or to differences in the perdurance of mRNA vs GFP-tagged protein reporters^42–44^. In addition, we independently validated a select number of instances of differential expression for which CRISPR/Cas9 engineered endogenous reporter alleles were available. We focused on DEGs between L1 and L4 in neurons with unknown function (CAN) or that undergo substantial postembryonic development (HSN), and on TFs and neuropeptides, which showed some of the largest fold changes (Tables S10, S11; Figure 5C-E, Figure S8B-F). For example, we confirmed decreased expression of the basic helix-loop-helix TF *hlh-32* in AVF in the L4 vs L1 (Figure 5C-E) and additional neuropeptide gene differential expression across a range of neuron classes (Figure S8B-D, Figure S9A-F).

To identify patterns across the nervous system and the genome, we clustered DEG profiles for both neurons and genes (Figure 5A-B). Our analysis revealed four major conclusions. First, we noted that transcripts encoding most ribosomal proteins, including *rps-4*, showed broad transient upregulation in neurons from embryo to L1 (Figure S10A-B), with significant downregulation by L4 (Figure S11A-B). Four ribosomal genes (*rpl-7A, rps-16, rpl-25.1, rpl-27*) showed a distinct pattern, with little change from embryo to L1, and widespread higher expression in L4 neurons (Figure S12A). We imaged endogenously tagged versions of three ribosomal proteins (RPL-25.1, RPL-7A, RPS-4) (Figure S12B-D). RPL-7A protein levels moderately decreased from L1 to L4, which differs from the pattern of *rpl-7A* transcripts (Figure S12B). However, RPS-4 and RPL-25.1 protein levels changed consistently with transcripts (Figure S12C-D). These data suggest highly dynamic regulation of protein translation during *C. elegans* nervous system development, consistent with reports from other organisms^45–47^.

Second, most neurons downregulate a shared set of genes in their early post-mitotic state. A set of 45 genes showed higher expression in the embryo compared to the L1 in embryonic neurons, (Figure S10A, see Methods) and higher expression in the L1 than L4 among postembryonic neurons (Figure 5B, dashed box, Figure S11A-B). These genes encode cell adhesion molecules, proteins involving in protein folding and processing, TFs, chromatin modifiers, histones and several genes with various or uncharacterized functions (Table S12). These genes also significantly changed along pseudotime in newborn neurons (Tables S7, S12). The earliest generated postembryonic neurons, AQR and PQR, have similar levels of these genes between L1 and L4, suggesting downregulation occurred by the late L1 period. These data highlight the cellular processes involved in early neuronal differentiation.

The third major conclusion is that subsets of functionally related neurons downregulate distinct sets of genes during early maturation. In both age comparisons, ciliated sensory neurons largely clustered together. Two gene clusters with shared patterns of higher expression in the embryo vs L1 across many ciliated sensory neurons (Figure S10A, dark blue) were enriched in genes involved in non-motile cilia formation and function (Figure S10B). An additional subset of genes showed a more restricted pattern of elevated expression in the L1 vs L4 only among postembryonic ventral cord motor neurons. Genes involved in mRNA splicing, the proteasome and chromatin modification were significantly enriched in this group (Figure S11A, B, magenta).

The fourth major principle is that maturation largely entails cell-type specific differential expression. Several pieces of evidence support this conclusion. Most genes showed significant differential expression in a small number of neuron classes (Figure 5F-G, median number of neurons with DE per gene = 3). Only five genes, *lep-5, pgal-1, ctc-3, hsp-70, pab-1*, showed broadly higher expression (>50% of neuron classes) in the L4 compared to L1. Notably, the non-coding RNA *lep-5* showed the broadest upregulation, consistent with its established role in promoting the larval to adult transition and sexual maturation^48,49^. Additionally, we found a cluster of genes (Figure S10A) with higher expression in the L1 vs embryo in selected ciliated sensory neurons. This cluster showed significant enrichment of insulin-related genes, *irld* family genes (denoted as “receptor L domain”) and seven transmembrane domain G-protein coupled receptors (Figure S10B).

A substantial number of genes showed differential expression in more than one neuron class. These genes could exhibit three possible patterns: 1) consistently higher expression in the younger stage, 2) consistently higher expression in the older stage, or 3) mixed changes, with higher expression in the younger stage for some neuron classes but higher expression in the older stage in others. Between the embryo and L1, ∼80% of genes with DE in > 1 class (3192/4011) consistently changed expression in the same direction across neurons (‘coherent’ genes). Between L1 and L4, 53% (2014/3823) of genes consistently changed expression in the same direction. The remaining genes (819/4011, 20%, in embryo–L1 and 1809/3823, 47%, in L1-L4) exhibited changes in opposite directions depending on the neuron class (‘incoherent’ genes). For example, the cysteine synthase enzyme *cysl-1* was higher in the embryo in 17 neuron classes, but higher in the L1 stage in 17 other neuron classes. Incoherent genes exhibited only slightly lower absolute fold-change values compared to coherent genes (embryo vs L1: incoherent median log2FC: 3.40, coherent median log2FC: 3.73, p-value: 6.9 × 10^−10^, Figure S10C; L1 vs L4: incoherent median log2FC: 2.18, coherent median log2FC: 2.4; p-value 4 × 10^−7^, Figure S11C). We identified several incoherent genes from both comparisons with log2 fold changes > 4 in different directions across cells (Table S13). These results highlight the cell-type specificity of neuronal maturation.

### Differential gene expression is correlated with neuronal maturation

We found medians of 571 DEGs per neuron class between the embryo and L1 (Figure S10D-E) and 227 DEGs between L1 and L4 (Figure S11D-E). No significant differences were seen in the number of DEGs between functional categories of neurons for the embryo-L1 comparison (Figure S10D-E). In the L1-L4 comparison, pharyngeal neurons had fewer DEGs than all other neuron categories (Figure S11D-E), even when postembryonic neurons were excluded. Postembryonic neurons exhibited significantly more DEGs between L1 and L4 than embryonic neurons (Figure S11F-G) likely due to neuronal maturation from the L1 to L4 stage.

The embryonic neuron HSN shows the third highest number of DEGs, likely due to delayed maturation of HSN compared to other embryonic neurons^50,51^. HSN neurons are generated in the embryo, but do not acquire neuronal features until L4 when they innervate newly generated vulval muscle^50^. The postembryonic VC neurons also differentiate and synapse with vulval muscle in late L4^4,24^. Consistent with delayed differentiation, the HSN and VC populations show significantly weaker expression of pan-neuronal marker genes vs other neurons at the L1 stage (Figure S8G-H). Along with genes showing higher expression in HSN at the L4 stage (Table S11), we also detected downregulation of genes upon HSN maturation. Among the 193 genes expressed at higher levels in HSN in the L1, we identified several genes that indicate possible roles of HSN in the early larval nervous system. At L1, but not L4, HSN expressed transcripts of the polycistronic innexin genes *inx-12* and *inx-13* (Table S11). These innexins are primarily expressed in non-neuronal tissues, including the epidermis, and their expression in HSN could facilitate communication between HSN and epidermis in early larval development. We also detected higher transcript levels in HSN at the L1 stage for genes involved in neuronal signaling, including the neuropeptide *srlf-1* (Figure S8B, E) and the putative acetylcholine-gated chloride channel *lgc-47* (Table S11). Several TFs were expressed in HSN in L1 but not L4, including *ham-2, bed-3* and the SOX family TF *sem-2*. Many of these same genes are also expressed at higher levels in L1 in the VC neurons. Notably, in the postembryonic M lineage, *sem-2* antagonizes terminal differentiation of muscle cells^52^ which suggests that *sem-2* may be acting to pause full neuronal maturation in HSN and VC.

### Gene regulatory network analysis to identify TFs driving developmental changes in gene expression

To identify putative transcriptional regulators driving differential expression across development, we used gene regulatory network (GRN) analysis. For our approach, we used recently described gene regulatory networks (GRNs)^53^ which assign likely targets to TFs based on ChIP-Seq experiments, DNA-binding motifs, and TF-gene promoter interactions from yeast one-hybrid assays^53–55^. We reasoned that it may be possible to infer changes in TF activity from changes in expression of its putative targets^56^ (see Methods). We identified 966 cases of significantly altered TF activity for 151 distinct TFs (Figure 5H, Table S14) and 355 instances of altered activity for 102 distinct TFs between L1 and L4 (Figure 5I, Table S15). Several of these instances were consistent with experimental work, including increased activity from embryo to L1 of FKH-8 in ciliated neurons^57^ (Figure 5H), decreased activity from embryo to L1 of the proneuronal factor NeuroD homolog CND-1 in many neuron classes^58–60^ (Figure 5H) and increased expression of LIN-14 targets from L1 to L4^20,61–69^.

Our GRN analysis yielded three main principles. First, most TFs exhibiting significant age-related changes in activity likely function in restricted subsets of neurons. TFs showed significant activity changes in a median of two neurons in the embryo-L1 and 1.5 neurons for the L1-L4 comparison, consistent with the differential gene expression results (Fig 5A, B). We did identify a subset of TFs with broad changes in activity (e.g., *hmg-4, nhr-2* and *ceh-39,* black arrows in Figure 5H-I). As these TFs most often showed decreases in activity (i.e., decreases in target expression), their loss of activity may be involved in the shared patterns of gene downregulation in early postmitotic neurons described above.

Second, our results support the idea that terminal selector regulatory activity is stable in post-mitotic neurons. Nearly all known terminal selector TFs are included in the *Cel*EsT GRNs, but these TFs comprise only a small fraction of the TFs with significant activity among DE genes (Figure 5H-I). Most neurons do not show evidence of significant changes in terminal selector TF activity across the development stages we compared. These observations are consistent with the stability of terminal selector TF expression patterns (Figure 4), and with the model that terminal selectors establish and maintain neuronal gene expression throughout the life of a neuron^70^.

Third, the changes in TF activity are not strongly coupled to significant changes in the expression of the TFs themselves. Most TFs with differential activity in a given neuron did not exhibit differential expression in that neuron (Figure 5J). This finding raises the possibility that post-transcriptional regulation of TF activity may be involved in nervous system development. Expression-based GRN analysis has limits, including incomplete coverage of TFs and TF-target assignments. Additionally, many target genes are listed in the regulons of multiple TFs. Therefore, if a group of co-regulated genes share common regulatory networks within the GRNs, it may result in significant putative activity changes for multiple TFs. These caveats notwithstanding, the detection of several instances of known TF regulation suggests that GRN analysis provides plausible hypotheses for TFs that may coordinate maturation in the *C. elegans* nervous system.

### Neuropeptidergic networks in the L1 larval stage

Neuropeptide-encoding genes were significantly enriched in the differentially expressed genes throughout larval development (Figure S10B, Figure S11B) and exhibited some of the highest log fold changes (Tables S10, S11). These neuropeptides, along with their receptors, form “wireless” networks^16,71–73^ of communication between neurons that are often unconnected by wired chemical synapses. We therefore wondered if developmentally-regulated differences in neuropeptide gene expression between L1 and L4 animals led to differences in the structure of their neuropeptide networks.

Using the same approach used in our earlier work^72^, we analyzed the L1 neuropeptide network (see Methods, Figure 6A-C, Figure S13A-C) and compared it to the L4 network. We limited our comparative analysis to the 291 neurons present at both the L1 and L4 stages and initially used the most stringent gene expression threshold. In general, the architecture of the neuropeptidergic network in the L1 resembled that of the L4. In both stages the network was very dense, with a decentralized organization, a defined mesoscale structure consisting of communities with similar input patterns, and specialized peptidergic hubs with high neuropeptide connectivity but modest synaptic connectivity. However, at the level of individual connections, we observed many cases where changes in neuropeptide expression altered peptidergic signaling networks. Across the 88 neuropeptide precursor (NPP)-GPCR pairs, 39% (35 pairs) showed expression changes leading to alterations in network edges (Fig 6A-C, Figure S14). Between 77% and 79% of neuron-to-neuron connections (for short- vs mid-range networks) were conserved across the two stages. For the mid-range network, 25% of conserved connections used identical NPP-GPCR pairs, 63% share at least 1 common NPP-GPCR pair, and 12% were topologically conserved but involved different molecular pairs (Figure 6B); similar percentages were seen for networks limited to short-range signaling or with a less stringent expression threshold (Figure S15). The number of conserved connections was significantly greater (p-value = 0.01) than expected by random chance (Figure 6B). L1 networks exhibited a modest 2–3% increase in overall connectivity compared to those in the L4 (Figure 6C).

**Figure 6.**
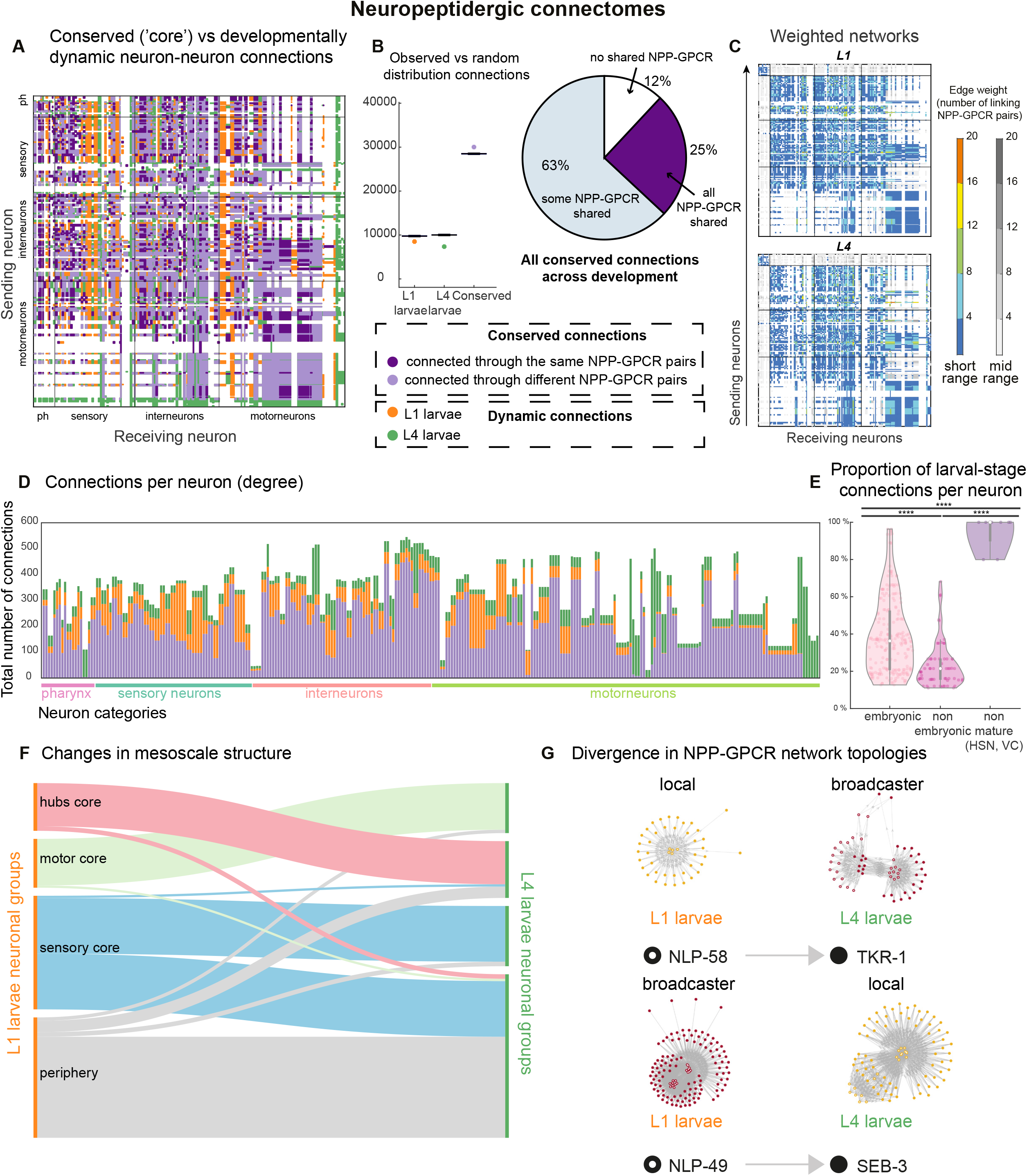
Developmental alterations in neuropeptide networks. A) Neuropeptidergic connectome (mid-range) showing the developmental pattern of connections between sending (y-axis) and receiving neurons (x-axis). Core (conserved) connections and developmentally dynamic connections are color-coded. Neurons are grouped by functional category (ph = Pharyngeal). B) Left: box plot showing observed value of stage specific connections (L1 and L4) and conserved connections as colored points vs. boxplots showing distribution of values for 100 random networks. Significance was established with a two-tailed test. Right: pie chart showing the percentage distribution of the conserved connections between NPPs and GPCRs forming the core connectome. 12% of those connections do not share a common NPP-GPCR pair across development. The other 88% share at least one NPP-GPCR pair (64%) or all NPP-GPCR pairs (23%). C) Weighted neuropeptidergic connectomes, colored by the number of NPP-GPCR pairs. D) Total number of connections (y-axis, mid-range networks) per neuron across development (x-axis) grouped by functional categories. Bars are color-filled according to the subsets of core and developmentally dynamic connections (See key in B). E) Proportion of developmentally dynamic neuropeptide connections (y-axis) per neuron classified by birthtime. Kruskal-Wallis test with Dunn’s correction, \*\*\*\**P* < 0.0001. F) Alluvial plot displaying the changes in mesoscale structure across development. Functional mesoscale groups at the L1 stage (left) vs the L4 stage (right). The same 4 groups (hubs core, sensory core, motor core, periphery) are conserved but some individual neurons change between groups. G) Network topologies of the indicated NPP/GPCR pairs. Empty circles represent NPPs, filled circles represent GPCRs. Local networks display restricted NPP and GPCR expression (≤50 neurons), pervasive networks display broad NPP and GPCR expression (>50 neurons), broadcaster networks display restricted NPP but broad GPCR expression. NLP-58 and NLP-49 = NPPs; TKR-1 and SEB-3 = GPCRs. See also Figures S13-S17 and Table S18.

We next investigated the importance of individual neurons in the L1 neuropeptidergic network. We compared the highly connected neuropeptidergic hub neurons from the L4 stage^72^ to those in the L1. Most of these peptidergic hub neurons are embryonically derived and exhibit high degree at the L1 stage, with a connectivity correlation >0.8 across stages (Figure S16A-B). These data support the notion that these neuropeptide hubs form the scaffold of the neuropeptide connectome, with their organizational role established from birth. Outside of this conserved core, stage-specific connections emerged, particularly in neurons undergoing significant developmental changes. Postembryonically born neurons like SDQ, VC, PVM, and AVM showed >80% stage-specific neuropeptide connectivity (Figure 6D-E, Figure S16C). Neuropeptide connectivity in these cases was restricted in the earlier stage by shorter processes (Figure S13B) as well as by large variations in gene expression between L4 and L1 stages. Among embryonically developed neurons, connectivity divergence was more prevalent in head neurons, such as OLQ, RMD, IL1 and CEP (Figure 6D; Figure S16C) linked to mechanosensation, a modality known to undergo functional refinement during development^74–76^.

Stage-specific changes in neuropeptide expression also altered the mesoscale structure of the neuropeptide connectome. We showed previously that the neurons in the L4 network can be sorted into four distinct communities based in their input patterns (see Methods, Figure 6F). While this structure was conserved in the L1, the neurons that cluster in each community changed. For example, the HSN and AVM neurons were in the peripheral group at L1 but in the hubs core in L4, reflecting their maturation (Figure 6F; Table S18; Figure S17). Conversely, the OLQ neurons were in the sensory core in L1, but peripheral in L4, suggesting age-related reduction in functional connectivity (Figure 6F; Table S18; Figure S17). To independently assess the community structures in L1 and L4 animals, we used a block modelling approach^77,78^, which resulted in nearly identical neuronal communities as the t-SNE/PCA approach (see Methods, Figure S17). Overall, both methods indicated a conserved mesoscale network structure, with some neurons changing community membership due to developmental reconfiguration of incoming connections.

The network topologies for individual neuropeptide-receptor pairs also underwent changes. We previously classified these networks as either “local” (restricted expression of both peptide and receptor), “broadcaster” (restricted peptide but broad receptor expression), “integrative” (broad peptide but restricted receptor expression) or “pervasive” (broad peptide and receptor expression). We observed some networks expanded their scope during development; for example, the *nlp-58/tkr-1* tachykinin-like homologue showed a “local” topology at L1 but a “broadcaster” topology at L4 (Figure 6G). In contrast, some pairs showed reduced scope, such as *nlp-49/seb-3*, which shifted from a “broadcaster” network at L1 to a more “local” network at L4, reflecting more restricted expression of both ligand and receptor (Figure 6G).

Neuropeptide expression is also sexually dimorphic in the *C. elegans* nervous system^79,80^ as well as highly divergent between *Caenorhabditis* species^18^. Collectively, these data identify neuropeptidergic signaling as a dynamic feature of nervous system function across age, sex and evolution.

### Identifying molecular determinants of neuronal membrane contact

We next sought to use the L1 transcriptomic data to identify candidate molecular determinants of nerve ring structure and connectivity. We detected expression of 298 transmembrane, GPI-anchored or secreted proteins (i.e. cell adhesion molecules, CAMs)^81–85^ in the L1 nervous system. Membrane contact is a prerequisite for synapses, and the extent of membrane contact can be predictive of synaptic connectivity^24,86,87^. We catalogued gene enrichment based on membrane contact for each of the 80 neurons in the L1 nerve ring. We detected 62 instances of gene enrichment in either contacted or non-contacted cells (Table S19). For example, the cadherin *hmr-1* and the immunoglobulin superfamily adhesion molecule *syg-1* were significantly enriched in cells that contact AVK vs cells with no AVK membrane contact. Conversely, the transmembrane low density lipoprotein receptor gene T13C2.6/*gldi-2* and the tetraspanin *tsp-21* were enriched in cells not contacting AVK (Figure 7A-B). Notably, both HMR-1 and SYG-1 have documented binding interactions with proteins whose transcripts are expressed in AVK (*hmr-1* and *sax-3*, respectively^85,88^), suggesting that these molecules may promote the membrane contacts of AVK.

**Figure 7.**
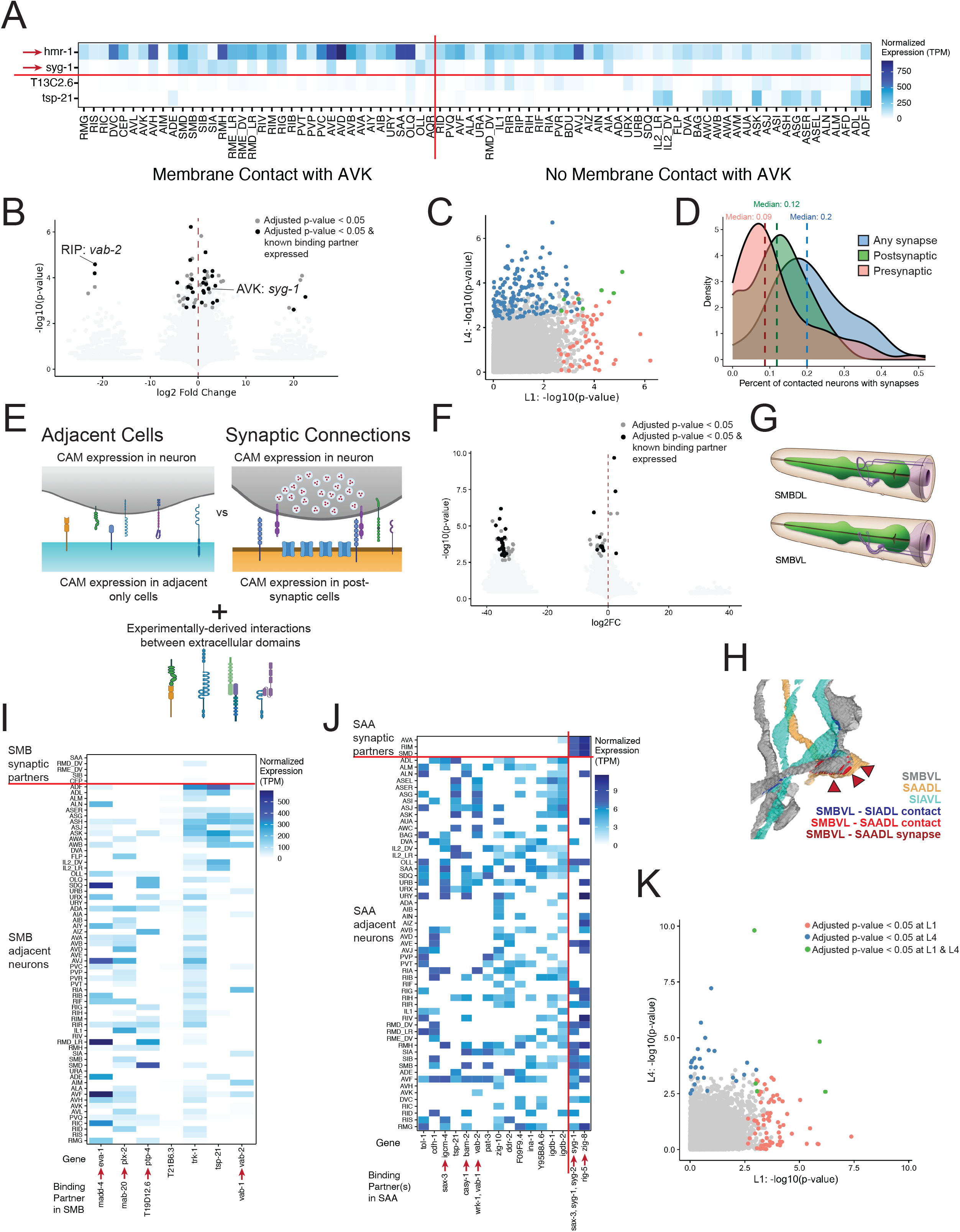
Mapping secreted and cell surface proteins for neurite fasciculation and synaptic connectivity. A) Enrichment of *hmr-1* and *syg-1* in neurons contacting AVK (left) vs neurons not contacting AVK (right). T13C2.6 and *tsp-21* (bottom) are enriched in neurons not contacting AVK. B) Volcano plot of CAM enrichment based on membrane contact. Each dot represents a gene tested for a given neuron. Log2FC > 0 = enrichment in neurons with membrane contact. Dark grey dots denote significant enrichment. Black dots denote significant enrichment in which at least one known binding partner is expressed in the tested neuron. C) Scatterplot showing adjusted p-values for each gene/neuron combination testing enrichment in membrane contacted vs non-contacted cells in L1 (x-axis) and in L4 (y-axis). Genes only detected at only one age are not shown. (Pink) Significant enrichment in L1, (Blue) Significant enrichment in L4. (Green) Significant enrichment in both stages. D) Density plot of the fraction of membrane-contacted neurons with synaptic connections in the L1 nerve ring. (Blue) Fraction of membrane-contacted neurons with any synaptic connections (Green) Fraction of neurons that receive postsynaptic output. (Pink) Fraction of membrane-contacted neurons that provide presynaptic input. E) Top: CAM expression in cells with membrane contact only (left) and in cells with synaptic connections (right). Bottom: Known CAM binding partners. F) Volcano plot for enrichment of CAMs based on synaptic connectivity. Each dot represents one gene/neuron comparison. Log2FC < 0 = enrichment in non-synaptic adjacent cells; log2FC > 0 = enrichment in synaptically-connected cells. Dot colors as in panel B. G) SMB neurons that innervate the nerve ring, from WormAtlas. H) EM reconstruction (NeuroScan) of late L1 nerve ring featuring three neurons: SAADL (orange) has both membrane contact (red patches) and synaptic contacts (red triangles) with SMBVL (gray), whereas SIAVL (light blue) has only membrane contact (dark blue) with SMBVL. I) Expression (TPM) of 7 CAMs enriched in SMB-non-synaptic adjacent cells (below) compared to in SMB postsynaptic partners (above). J) Expression of 14 genes (left) enriched in SAA-non-synaptic, adjacent neurons and 2 genes (right) enriched in SAA-synaptic partners. Red arrows in A, I, J denote genes encoding proteins with known binding partners expressed in the indicated neuron. K) Scatterplot of p-values for each gene-neuron combination testing enrichment in synaptically connected cells in L1 (x-axis) and in L4 (y-axis). Genes detected at only one age are not shown. Colors as in panel C. All p-values adjusted using Benjamini-Hochberg procedure. See also Figure S18 and Tables S19-S21.

We analyzed CAM expression in the L4 nerve ring to determine if enrichment in contacted or non-contacted cells is maintained across larval development. Surprisingly, of the 150 cases of enrichment in the L4, only seven transcripts were also significantly enriched in contacted vs non-contacted neurons at the L1 stage (Figure 7C). The distinct patterns of gene enrichment in contacted vs non-contacted cells at L1 and L4 may reflect differences in genes involved in establishing vs maintaining membrane contact in the *C. elegans* nervous system^67,89,90^.

### Identifying determinants of connectivity from a neuron-specific transcriptome

Electron micrograph reconstructions of the nerve ring at L1 revealed that a neuron is synaptically connected to a limited fraction of cells with which it has membrane contact (Figure 7D). Although previous work determined that neuron adjacency in the nerve ring is predictive of connectivity, this correlation is selective for neuronal contacts that exceed a minimum threshold^91^. We reasoned that molecular mechanisms that either promote or inhibit the formation of synapses between neurons may be distinct from those that govern membrane contact (Figure 7E). We therefore sought to identify CAMs differentially expressed in synaptically connected neurons vs neurons with only membrane contacts (see Methods).

We identified 75 cases of enriched expression patterns when comparing non-synaptic adjacent neurons vs postsynaptic outputs of a given neuron (Figure 7F), many involving genes with known binding partners^85^. Most of the significant cases (70/75) involved enrichment of genes in non-synaptic adjacent cells. For example, in the L1, the four SMB neurons contact 67 neuron types but only synapse onto five (Figure 7G-H)^6,24,86^. We detected significant enrichment of seven genes in SMB-adjacent, non-synaptic neurons compared to SMB-postsynaptic partners (Figure 7I, Table S20), including the semaphorin receptor *plx-2* and ephrin *vab-2* which have known binding partners expressed in SMB (*mab-20* and *vab-1,* respectively). Semaphorins and ephrins can mediate intercellular repulsion^92–96^, suggesting that *mab-20/plx-2* and/or *vab-2/vab-1* signaling may antagonize the formation of SMB synaptic output.

The SAA neurons showed the highest number of enriched genes (sixteen), with 14 enriched in non-synaptic, adjacent cells, and two transcripts enriched in synaptically connected cells (Figure 7J). This includes enrichment of *zig-8* in SAA postsynaptic partners and expression of the ZIG-8 interacting Ig-domain IgLON molecule *rig-5* in SAA. Notably, RIG-5 interaction with ZIG-8 specifies connections in the ventral nerve cord^97^. In addition, closely related IgLONs in *Drosophila melanogaster*, DIPs and Dprs, promote selective adhesion in the developing nervous system^98–101^.

The analysis of each neuron and its presynaptic inputs in the L1 revealed 247 instances of significant gene expression enrichment in a presynaptic partner vs adjacent cells featuring 94 genes across 29 neuron classes (Figure S18A, Table S21). Only one instance involved a gene selectively enriched in synaptic partners: *wrk-1* is enriched in the presynaptic partners of RMH relative to solely adjacent neurons. The AIY neuron provides an example of genes enriched in non-synaptic, adjacent cells (Figure S18B-D). We noted that transcripts for a given gene may be enriched in the synaptically connected cells for one neuron, but in adjacent, non-synaptic partners of a different set of neurons. For example, in addition to its enrichment in presynaptic inputs to RMH, *wrk-1* is enriched in adjacent, non-synaptic partners of four neurons: DVA, URX, RIG and SIA. A given CAM could differentially affect connectivity via interactions with distinct molecules on adjacent cells. Most genes with enrichment in adjacent, non-synaptic partners were enriched in the adjacent cells of only a small number of neuron classes (median = 2 neuron classes). The genes showing the broadest enrichment patterns in adjacent cells were significant in fewer than 10 classes. These results are consistent with a model in which individual CAMs may regulate the connectivity of a small subset of neuron classes.

We note that the limited number neurons in the synaptically-connected group constrains the power of our statistical test. For example, several genes are expressed in a subset of presynaptic inputs to AIA and in none of its non-synaptic, adjacent neurons (Figure S18E). Similar cases can be identified by high log fold changes (Tables S20, S21) and provide attractive hypotheses for molecules that may drive synaptic specificity in individual neuronal circuits.

Most (80%) synapses added during larval development involve the strengthening of existing connections; only 20% are established between previously unconnected neurons^6^. Mechanisms that sustain synaptic specificity throughout development may differ from pathways that define initial connections^90,102–104^. We therefore compared gene enrichment among synaptic or adjacent partners between the L1 and L4 stages. Only a limited number of instances showed significant enrichment at both ages (Figure 7K, Figure S18F). This observation suggests that distinct molecular mechanisms may underlie the initial patterning of synapses and subsequent maintenance.

Our scRNA-seq data can be explored at the CengenApps: L1: https://cengen.shinyapps.io/L1app; updated L4 data: https://cengen.shinyapps.io/L4app. We expect that these data will inspire future studies of genes, neurons and circuits, and will allow for mechanistic insight into neuronal development and maturation across an entire nervous system.

### Limitations of the study

Profiles of 13 under-represented neuron classes are likely incomplete. Our scRNA-seq method fails to capture non-polyadenylated transcripts and many non-coding RNAs. Some DEGs between embryonic and L1 neurons and between L1 and L4 neurons may result from different methods of isolation for embryos and larvae or differences in cellular responses to dissociation between stages. As the nerve ring is primarily formed during late embryogenesis, transiently expressed molecular determinants may not be detectable by mid-late L1. In addition, some of the enrichments we identified may not be causal.

## RESOURCE AVAILABILITY

### Lead contact

Requests for further information and reagents should be directed to the lead contact, Seth Taylor.

### Materials availability

Strains used in this study can be obtained from the *Caenorhabditis Genetics Center* or upon request from the lead contact.

### Data and code availability

- Single-cell RNA-seq data are publicly available at GEO under the accession number GSE310667.
- scRNA-seq data are available at the L1 CengenApp (https://cengen.shinyapps.io/L1app) or the L4 CengenApp (https://cengen.shinyapps.io/L4app).
- Imaging files can be accessed at Mendeley Data with at doi: 10.17632/nkt7gd9hm4.1.
- Original code is available at github: https://github.com/cengenproject and on Zenodo at doi: https://doi.org/10.5281/zenodo.23753627.
- Additional information required to reanalyze our data is available from the lead contact upon request.

## Acknowledgements

We thank the CeNGEN Advisory Board for guidance. O.H. is an Investigator with the Howard Hughes Medical Institute. FACS (Flow Cytometry Shared Resource, supported by Ingram Cancer Center [P30 CA68485], DDRC [DK058404]), scRNA-seq (VANTAGE, supported by CTSA [5UL1 RR024975-03], Ingram Cancer Center [P30 CA68485], Vision Center [P30 EY08126], and NIH/NCRR [G20 RR030956]), and confocal imaging (Cell Imaging Shared Resource [NIH CA68485, DL20593, DK58404, DK59637, and EY08126]) were performed at Vanderbilt. Strains were provided by the CGC (NIH P40 OD010440A). All research at the Department of Psychiatry in the University of Cambridge is supported by the NIHR Cambridge Biomedical Research Centre (NIHR203312) and the NIHR Applied Research Collaboration East of England. The views expressed are those of the author(s) and not necessarily those of the NIHR or the Department of Health and Social Care. This work was funded by the Research Foundation Flanders (FWO) grant G0B5322N, the Baillet Latour Fund, and NIH (R01NS100547 to M.H., O.H., D.M.M., R01 NS110391 to O.H., and R01NS113559, R01NS106951 to D.M.M).

## Author contributions

Conceptualization: S.R.T., M.H., O.H., D.M.M. III; Methodology: S.R.T., E.V., D.M.M. III; Formal Analysis: S.R.T., C.O., A.A., E.G., E.C., A.W., L.R.S., S.G.; Investigation: S.R.T., C.O., R.M., L.R.S., S.T.B., D.H., A.R., J.P., G.V., G.R.A., D.M.M., M.M., M.E; Software: A.W., S.R.T.; Supervision: S.R.T., I.B., P.V., W.R.S., O.H., D.M.M. III; Funding acquisition: M.H., O.H., D.M.M. III, S.K.; Writing – original draft: S.R.T., C.O., D.M.M. III; Writing – review & editing: S.R.T., D.M.M. III, M.H., O.H.

## STAR Methods

### EXPERIMENTAL MODEL AND STUDY DETAILS

#### *C. elegans* growth and maintenance

All animals used were *C. elegans* hermaphrodites ranging from the L1 to adult stages. Animals were cultured at 20° C unless otherwise described and maintained on NGM agar plates seeded with OP50-1 bacteria or on 8P agar plates seeded with NA22 bacteria.

#### Strain List

The strains used in this study are described in Table S1.

### METHOD DETAILS

#### Mesh Synchronization

To obtain synchronized cultures of mid-late L1 animals without subjecting them to L1 arrest, which alters gene expression^105^, we scaled up worm cultures by chunking ¼ of a 60 mm plate of recently starved worms onto a 100 mm plate seeded with OP50-1. After culturing for 2-3 days at 20 C, 1/5 of each plate was chunked onto 8P nutrient agar 150 mm plates seeded with *E. coli* strain NA22. Worms were cultured at 20 C for 3-4 days. Approximately ten 150 mm plates were used for each strain to be sorted (see below). Five 150 mm plates were used for sorting controls. We scraped worms and eggs off 150 mm plates with a cell scraper and pelleted by centrifugation at 150 rcf for 2.5 minutes. Embryos were isolated by hypochlorite treatment of the pellet and floatation in 30% sucrose. Embryos were washed 2x in M9 and placed on a nylon mesh grid with 10-micron or 18-micron pores. Embryos will not pass through the pores of the mesh, but hatched L1 larvae are able to crawl through. The mesh was placed on M9 containing NA22 bacteria, and embryos were allowed to hatch at room temperature. Larvae collected in the first 1.5 hours after completion of hypochlorite treatment were discarded, and the mesh grid was placed over a fresh collection media of M9 with NA22 bacteria for 2-4 hours. Hatched L1s were pelleted by centrifugation and plated onto fresh 8P x 150 mm plates with NA22 and incubated at 20° C for 12-17 hours. This timetable yields larval populations synchronized to within 14-20 hours after hatching. Where possible, we estimated the developmental stage of cultures by counting fluorescent protein expression in postembryonic neurons that are born during the first larval stage (L1). For example, with strain OH10689, which features rab-3::2xNLS-TagRFP in all neuronal nuclei, we counted the number of RFP+ neurons in the ventral nerve cord. For strains without fluorescent labeling of easily identifiable postembryonic neurons, we cultured worms for similar durations of time post hatching. Estimates of developmental staging are found in Table S2.

#### Dissociation

Single cell suspensions were obtained with modifications from prior work^16,106–109^ as follows. Synchronized larvae were collected and separated from bacteria by washing twice with ice-cold M9 and centrifuging at 150 rcf for 2.5 minutes. Washed larvae were transferred to a 1.6 mL centrifuge tube and pelleted at 16,000 rcf for 1 minute. 100-250 μL pellets of packed worms were treated with 500 μL of SDS-DTT solution (20 mM HEPES, 0.25% SDS, 200 mM DTT, 3% sucrose, pH 8.0) for 2 minutes. Following SDS-DTT treatment, worms were washed five times by diluting with 1 mL egg buffer and pelleting at 16,000 rcf for 30 seconds. Worms were then incubated in pronase (15 mg/mL, Sigma-Aldrich P8811, diluted in egg buffer) for 23 minutes on a rocking nutator. During the pronase incubation, worms were mechanically disrupted by pipetting through a P1000 pipette tip for four sets of 80 repetitions. The status of dissociation was monitored under a fluorescence dissecting microscope at 5-minute intervals. The pronase digestion was stopped by adding 750 μL L-15 media supplemented with 10% fetal bovine serum (L-15-10), and cells were pelleted by centrifuging at 530 rcf for 5 minutes at 4° C. The pellet was resuspended in L-15-10, and single cells were separated from whole worms and debris by centrifuging at 150 rcf for 2 minutes at 4° C. The supernatant was passed through a 35-micron filter cap into a 5 mL round bottom tube. The pellet was resuspended a second time in L-15-10, spun at 150 rcf for 2 minutes at 4° C, and the resulting supernatant was added to the collection tube.

#### Fluorescent-Activated Cell Sorting

Fluorescence Activated Cell Sorting (FACS) was performed on a BD FACSAria™III Cell Sorter equipped with 70-micron diameter nozzles^16,26,109^. DAPI was added to the sample (final concentration of 1 μg/mL) to label dead and dying cells. We used a variety of fluorescent reporter strains to label subsets of neuronal cells (Table S2). For example, we used an *acr-2::GFP* reporter (strain CZ631) to target ventral nerve cord cholinergic motor neurons and an *unc-47::GFP* reporter (strain EG1285) to target GABAergic neurons. N2 worms lacking fluorescence and single-color controls (for intersectional labeling strategies and dual-color sorts) were used to set gates to exclude auto-fluorescent cells and to compensate for spillover between fluorescent channels. In three experiments, single-cell suspensions from separate strains were combined prior to FACS: (1) NC3583, NC2957; (2) OH16003, EG1285, ZM5488 and *uIs152*; (3) CZ631, NC3523, PS3504. Cells were sorted under the “4-way purity” mask. Cells expressing the desired fluorescent proteins were collected into 400 μL of chilled (4° C) L-15 media containing 33% FBS (L-15-33) and stored on ice. We concentrated the collected cells by centrifuging at 500 rcf for 5 minutes at 4° C. We removed all but ∼30-50 μL of the supernatant, resuspended the cells and counted fluorescent cells on a hemocytometer. Cell suspensions loaded into the 10X Chromium Controller ranged in concentration from 350-1000 cells/uL.

From one preparation of sorted cells, we generated both a live-cell sample and a methanol-fixed sample after sorting. This preparation contained strains NC3583 and NC2957. The live cell sample was prepared as described above. After concentrating the sample, we had ∼117 μL of sample at ∼1000 cells/μL. We submitted 22 μL for the live-cell sample. The remaining amount was fixed in methanol by adding 1 mL of ice-cold methanol drop-wise while gently vortexing the cells, followed by an additional 3 mL of ice-cold methanol. The cells were stored at −20° C for 22 days. Fixed cells were pelleted by centrifugation at 1000 rcf for 12 min at 4° C. We removed 4.05 mL of the supernatant. The fixed cells were suspended in 400 μL of egg buffer with 1% BSA and 2.5 U/μL RNAse inhibitor (NEB M0314L). Resuspended cells were concentrated by centrifuging at 800 rcf for 12 min at 4° C. 350 μL of the supernatant was removed and cells were gently resuspended. An aliquot was taken, DAPI was added (to concentration of 1 μg/mL), cells were counted on a hemocytometer based on the DAPI signal and then loaded onto the 10X Chromium Controller.

#### Single-cell RNA-Sequencing

Each sample (targeting 10,000 cells per sample) was processed for single cell 3’RNA sequencing utilizing the 10X Chromium system. Libraries were prepared using P/N 1000121 following the manufacturer’s protocol. The libraries were sequenced using the Illumina NovaSeq 6000 with 150 bp paired end reads. Real-Time Analysis software (RTA,version 2.4.11; Illumina) was used for base calling and analysis was completed using 10X Genomics Cell Ranger software (v6.0.0), and a custom reference genome based on Wormbase release WS273^110^ with extended 3’ UTR sequences^16^.

#### Data Analysis

Data analysis downstream of CellRanger was performed in R. We used the DropletUtils R package (version 1.6.1) function EmptyDrops^111^ to distinguish cells from empty droplets. We used a threshold of 50 UMIs for all samples except sample 8672-ST, for which we set the threshold at 90 UMI due to apparently higher background contamination, as evidenced by a pronounced plateau when plotting the number of UMIs in each barcode. Contamination from ambient RNA was estimated and corrected using the SoupX R package (version 1.6.1)^112^. The parameters and genes used to estimate the amount of contamination for each sample are listed in Table S2. We performed quality control with the scater package (1.24)^113^, using the percentage of UMIs mapping to the mitochondrial transcripts nduo-1, nduo-2, nduo-3, nduo-4, nduo-5, nduo-6, ctc-1, ctc-2, ctc-3, ndfl-4, atp-6, ctb-1, MTCE.7, and MTCE.33. The individual datasets for each sample were then combined and further processed using the monocle3 package (version 1.2.9)^17,22,114,115^. Genes detected in fewer than 5 cells were removed, and data were log-normalized. We performed dimensionality reduction using PCA (using 100 PCs, based on an elbow plot showing the variance explained by each principal component), followed by UMAP (with umap.min_dist = 0.3, umap.n_neighbors = 75). We performed clustering on the cells with both the monocle3 “cluster_cells” algorithm on UMAP coordinates using the default leiden settings (res = 1e-3), and with the Seurat “FindClusters” algorithm using the principal components. The two approaches yielded similar results.

#### Annotation of cell types

The majority of the cells were found in discrete groups in the UMAP embeddings (see Figure 1B, Figure 1D, Figure S3A), and these discrete groups were assigned cell type identities based on the expression of known cell-type specific genes from previous single-cell datasets and from the literature^16,25,26,116^. HSN and VC, though lacking pan-neuronal markers at the L1 stage, were annotated based on the combination of known gene expression (Table S4, Figure S2). HSN was identified by the co-expression of *egl-5*, *unc-86* and *ham-2* (Figure S2A-D). VC was identified by the co-expression of *ham-2, ceh-6,* and *lin-39* (Figure S2E-F).

Some cell groups in the global UMAP expressed markers of progenitor populations (e.g., cell cycle genes) and postmitotic neurons. We generated sub-UMAPs of these populations and could identify separate populations from the T, Q and P lineages based on known TF expression (Table S4). We then generated subsets for each lineage. We used both UMAP and PHATE embeddings to visualize the expression of cell-type specific genes for each lineage. The UMAP projection of the Q lineage displayed distinct AVM and PVM populations, whereas the PHATE projection did not. For the T lineage, we found that both UMAP and PHATE displayed separate progenitor and post-mitotic neuron populations, but the PHATE projection showed a more continuous pattern with branching reflecting the known lineage divisions. For the P lineage, the UMAP reductions featured structure that reflected the known lineage branches and divisions based on independent expression data. PHATE reductions of these same datasets lacked this structure. Therefore, we selected to present UMAP projections for the Q and P lineages and PHATE projections for the T lineage.

The T.p-derived neurons and some of their progenitors express the POU-domain TF *unc-86*^116–118^ and the posterior Hox genes *php-3* and *nob-1*^116,117^. We annotated multiple clusters as T.p lineage neuronal progenitors based on co-expression of *cdk-1*, *unc-86*, *nob-1* and *php-3* (Figure S4C, data not shown). Several TFs were expressed in specific subsets of progenitor cells, including the pro-neural ATOH1 bHLH TF homolog *lin-32*. We validated this finding *in vivo* with an endogenously tagged *lin-32* GFP reporter that is initially expressed in T.pp, and later in its descendants, T.ppa (PVW) and T.ppp (Figure S4E-F).

The EVX1/2 TF homolog *vab-7* is expressed in PVW and in a distinct progenitor cluster that gives rise to PHC (Figure S4G). We confirmed expression of the VAB-7 protein in these lineages with an CRISPR/Cas9 engineered GFP-tagged *vab-7* reporter allele that is detected in PVW, its sister (the T.ppp neuroblast), and daughter cells, including PHC (Figure S4 G-H). We used these combined results to annotate the T.pp lineage (Figure 2D). Additionally, we detected a distinct group of cells among T-lineage progenitors with strong *unc-86* and *egl-1* expression (data not shown). This cluster had few other strongly enriched genes beyond *egl-1*, and fewer genes and UMIs than other T-lineage cells. Given the role of *egl-1* in apoptosis^119,120^, we annotated this cluster as T.pppp, which undergoes apoptosis during normal development.

P cell-derived progenitors were identified by expression of cell cycle genes (e.g., *cdk-1*)^121–123^ (Figure S4K) vs post-mitotic cells that express markers (e.g., *sbt-1*)^124,125^ for mature neurons (Figure S4L). Notably, we observe strong concordance with a set of 49 TFs previously assigned to postembryonic neuronal progenitors on the basis of GFP reporter analysis^126^ (Figure S5O). Specific progenitor types (e.g., Pn.aa, Pn.ap) were annotated by expression of known marker genes (Figure 2K, L, O; Table S4). Notably, P-lineage clusters form continuous trajectories in the UMAP embedding with the earliest progenitors at one end of the projection and post-mitotic neurons at the other (Figure 2I-J). This embedding reflects the pattern of known cell divisions and suggests transcriptional changes as cells divide and mature. We noted two distinct sets of P-lineage cells that appeared to correspond to P0-P1 and P2-P12 lineages. To confirm the identities of P0-P1 and P2-P12 derived cells, we examined Hox genes, which distinguish subclasses of adult motor neurons. Adult motor neurons derived from P0-P1 (e.g., VA1, VD1, VB2, VB1) do not express Hox genes (with the exception of AS1 and VD2, which both express an endogenous *ceh-13* reporter^26^), whereas other P-lineage-derived motor neurons express at least one of the Hox genes *ceh-13, lin-39,* or *mab-5*^26^. On that basis, we confirmed that P0-P1 derived cells in our dataset lack Hox gene expression (Figure S5C-F), whereas P2-P12 derived cells show consistent expression of Hox genes (Figure S5G-J). The K cell and DVB were not identified in the data, likely due to a lack of labeling in the mid-to-late L1 stage. The DVB neuron does not begin expressing *unc-47* or pan-neuronal markers until mid-L2 or later, after the developmental stages we sampled^127^.

We were unable to identify subclasses of several ventral nerve cord motor neurons classes (VA, VC, VB, AS) which have been previously described in L4 or adult datasets^16,26^. A substantial proportion of marker genes enriched in these subclasses in the adult^26^ were detected in the L1 data (Table S5), sometimes in a similar fraction of cells. For example, *C24H12.1* is detected in 2.1% of adult VA neurons, where it is highly enriched in adult VA2 (Table S5). Single-molecule FISH at the L2 stage^26^ showed the same pattern of VA2 enrichment. In the L1 VA cluster, *C24H12.1* is detected in 4.5% of the VA cells, but these are distributed throughout the VA population in UMAP space (Figure S5N). Many other subclass-specific markers were not detected at L1 or were present at much lower levels (Table S5).

#### Reanalyzing published embryonic data

We re-analyzed single-cell data from *C. elegans* embryonic cells to obviate potential artifactual results arising from differences in analysis pipelines^22^. We downloaded fastq files from the SRA archives for GSE126954. This series includes data from 15 separate 10X data sets. We processed these files through CellRanger (v8.0.0), using the same custom WS273 reference genome used for the L1 data. A different reference genome was used in the original analysis of these data. We then processed the raw feature x barcode matrices using EmptyDrops. In the original analysis, the authors initially used stringent UMI count thresholds to distinguish cells from empty droplets. We reasoned that this approach may have excluded many neurons, including possible additional classes not originally annotated. From the processed embryonic data available at https://cello.shinyapps.io/celegans/, we extracted the unique cell barcode IDs and corresponding metadata from each individual experiment. We generated new filtered feature x barcode matrices that retained all the same cell barcodes from the original analysis as well as cell barcodes detected with our EmptyDrops method. We then used SoupX, scater and monocle3 to further process the data. Details for SoupX contamination correction of embryonic datasets can be found in Table S9.

We then merged the individual datasets and performed dimensionality reduction. We used pan-neuronal markers *sbt-1* and *egl-21*^16,17,22^ to identify post-mitotic neuronal populations. We subset clusters containing neurons and repeated dimensionality reduction, using 50 principal components and umap.min_dist = 0.3, umap.n_neighbors = 75). Several discrete populations in the UMAP embeddings could be identified as anatomical neuron classes. Other populations contained original cell type annotations (from the source data) for multiple cell classes or cell-type specific markers of multiple neuron classes from reference datasets. We subset these clusters and re-ran dimensionality reduction and subsequent embedding to increase resolution. In some cases, we reduced either the number of PCs calculated (based on the elbow plots displaying the variance explained) or the umap.min_dist and umap.n_neighbors parameters. In each round of subsetting, cells were clustered and cluster markers were identified using the monocle3 “top_markers” function. Cluster markers were compared to expression of L4 neuronal data using the CeNGENApp and to expression atlases of homeodomain TFs^116,128^ to annotate cell identity. In addition to clustering with previously identified embryonic cell types, newly detected embryonic neurons also appeared as distinct clusters in the UMAP. Using gene expression patterns from previously published single-cell datasets and the literature, we annotated most of these clusters. In each case, annotation of a distinct cell type was never solely made based on separation in UMAP space; rather, it was determined by clear correspondence of marker genes to known patterns of expression in either the literature or the L4 neuronal reference. In total, we annotated 92 neuronal classes. Of these, 89 types had a matching type in the L1 data. The three classes that did not match were instances in which subclasses were detected in L1 that could not be unambiguously distinguished in the embryonic data (AWC, RMD, and RME in embryo; AWC^ON/^AWC^OFF^, RMD LR/RMD DV, and RME LR/DV in L1). Our reanalysis identified 28 additional neuronal classes that were not conclusively annotated in the original embryonic data.

#### Pseudotime analysis and differential expression

We calculated pseudotime values for post-mitotic neurons. We subset data into groups of immediate neuronal progenitors and their daughter cells (e.g., P0.p, VA1 and VD1; Q.pa, AVM/PVM and SDQ). We performed dimensionality reduction using PCA (12 PCs) and clustered the cells on the PCs. We used the principal components and the clustering results as input to the slingshot R package^129^, with the cluster corresponding to the progenitor population as the starting point. The output of slingshot includes pseudotime values for individual cells. We then used the associationTest function from the tradeSeq R package^130^ to test for genes that change with respect to pseudotime. In some cases, the extreme edges of pseudotime contained few cells, which can lead to unstable and inaccurate model fitting. We therefore used the cells within the 0.01 - 0.99 quantiles for each neuron type as input to the tradeSeq fitGAM function. In the fitGAM function we used nknots = 4 and in the associationTest function we used nPoints = 8, contrastType = “end” and inverse = “Chol.” All other values were run at default settings. We performed the association test on each post-mitotic neuron independently. We calculated adjusted p-values using the Bonferroni method within each neuron’s analysis and retained only genes with adjusted p-values < 0.05.

We performed this analysis for all 16 postembryonically derived neurons. We also calculated pseudotime (as above) and the association test for 15 embryonic neuron classes which had > 100 cells and an annotated immediate progenitor class from our re-analysis of the Packer et al (2019) datatset. The embryonic neurons tested were: ASJ, AUA, AFD, RMD, ASH, RIB, ASG, AWA, ADF, AWB, RIV, AVB, RIA, RIC, and RIM. Cells that were present in the original Packer data have an estimated embryonic time value “raw.embryo.time.” Our re-analysis added additional cells which lack this value. For cells that have both raw.embryo.time and pseudotime values, we performed Pearson correlation.

#### Comparison of larval vs embryonic neuronal progenitors and newborn neurons

We calculated aggregate pseudobulk TPM expression profiles for each progenitor and postmitotic neuron class used in the pseudotime analysis. For this analysis, we included an additional set of embryonic motor neuron progenitors and neurons (DA, DB, DD neurons). These cell types were not suitable for the pseudotime analysis due to uncertain annotations in the early post-mitotic states (a large cell population annotated as DA_DB in the original paper contains markers of both the DA and DB neurons and could not be disentangled). Distinct DA and DB populations were annotated, but only from late embryonic timepoints. We generated the aggregate TPM value by normalizing UMI counts for each cell by a size factor (the monocle3 software package generates this as the UMI count per cell divided by the geometric mean of UMI counts for all cells), taking the mean normalized expression of each gene across the single cells corresponding to a cell type, and rescaling the resulting values to sum to 1,000,000^22^. We performed pairwise Spearman’s correlations between cell types and clustered the resulting correlation matrix using the pheatmap R package, with the distance method = “correlation” and clustering method “ward.D2.” We then used the igraph R package^131–133^ to construct a force-directed network based on pairs with Spearman’s rho correlations of 0.7 and above.

#### Reannotation of L4 single-cell data

During annotation of the L1 single cell data, we identified subsets of cells for several neuron types that showed high expression of genes involved in unfolded protein responses (UPR). We annotated these separately from cells without high levels of UPR-related genes and excluded them from further analysis to avoid expression artifacts likely resulting from the dissociation procedure. For direct comparisons with the L1 data, we also removed cells with high UPR responses from the previously obtained L4 data set^16^. Additionally, both the current L1 single-cell dataset and recently published data from adult animals^26^ resolved some cell types that were not originally identified in the L4 data set. Using marker genes described in the L1 and adult datasets, we were able to separate the DD and VD neurons and identify a distinct VA1 population in the L4 data.

#### Differential Gene Expression

Differential gene expression analyses between stages were completed using the Wilcoxon rank-sum test in Seurat (v4.4.0 for P lineage comparisons; v5.3.0 for L1 vs embryo, L1 vs L4)^134–137^. For the L1 to L4 comparisons, monocle3 cell_data_set objects containing the L1 and L4 neuronal single-cell datasets were combined into one object. This object was then converted into a Seurat object. 121 neuronal cell types were present in both stages. For each cell type, the Seurat object was subset to include only that cell type and the data were then normalized using the Seurat function NormalizeData with default parameters. The FindMarkers function was used to identify differentially expressed genes between L1 and L4 cells. The following parameters were used: ident.1 = “L4,” ident.2 = “L1,” logfc.threshold = 0, min.pct = 0. With these parameters, positive average log2 fold change values indicate higher expression in L4, whereas negative log2 fold change values indicate higher expression in L1. The results from all 121 separate comparisons were compiled into one table. We filtered the results to retain comparisons based on four criteria: 1) an adjusted p-value of < 0.05; 2) an absolute average log2 fold change of >= 1; 3) the gene must have been detected in > 10% of the cells in the higher-expressing stage; 4) the gene must have been detected in > 5 cells in the higher-expressing stage. Additionally, we identified 23 genes whose quantitative expression is unreliable in either the L1 or L4 data as their genomic loci were overexpressed in the transgenic reporter strains (Table S8). These genes were also removed from the differential expression results.

We used the pheatmap R package^138^ to visualize differential expression results. We removed genes that did not show any significant differential expression, resulting in 5810 genes. For visualization, we show the average log2 fold change values for these genes across all neurons. The embryo to L1 differential gene expression analysis was performed in the same way between 89 neuron classes present in both datasets. For this comparison, negative log2 fold change values indicate higher expression in the embryo, and positive values indicate higher expression in L1.

We used the following criteria to identify genes broadly downregulated in early postmitotic neurons from both comparisons: genes had to be significantly differentially expressed in more than 10 neuron classes between L1 and L4, with at least 75% of those cases higher at L1 and they had to also be differentially expressed in > 40 neuron classes between the embryo and L1 with higher expression in the embryo in at least 75% of the cases. Forty-five genes met these criteria. We used linear regression including functional categories or neuron birthtimes and the number of cells present at each age as covariates to account for sampling differences.

For each differential expression comparison within the Q, T and P-lineages, Seurat objects of each lineage dataset was subset to include only the two cell types to be compared. The FindMarkers function was used to identify differentially expressed genes between the two cell types, with parameters min.logFC.threshold = 0 and min.pct = 0. For parent-daughter cell comparisons, ident.1 was set to the parent cell and ident.2 was set to the daughter cell, so positive log2FC changes indicate higher expression in the parent. For sister-sister cell comparisons, ident.1 was set to the anterior sister and ident.2 was set to the posterior sister, so positive log2FC changes indicate higher expression in the anterior sister. The results were filtered on 4 criteria: (1) adjusted_p_value < 0.05; (2) log2FC > 1; (3) gene detected in 10% cells in the higher expressing cell type; (4) and gene detected in >5 cells in the higher expressing cell type.

For comparisons involving VD, we combined VD, VD2, VD12, and VD13 cells. For comparisons involving the P0.aa-AVF-VB02 and P1.aaa-AVF-VB01 sub-lineages, we combined VB01 and VB02 cells. We used the pheatmap R package for visualization of differential expression results. For Figure 3K, we filtered the P-lineage differential expression for TFs differentially expressed between at least one pair of directly related cell types under the above parameters. We show single-cell scaled TPM values for each gene across a subset of P-lineage cell types.

#### Gene Set Enrichment Analysis

Gene set enrichment was performed in R using Wormcat 2.0 annotations^139^. The whole genome annotations (version from Nov 11, 2021) were downloaded from www.wormcat.com and the wormcat R package was installed from GitHub. The background gene set for genes from the embryo vs L1 comparison consisted of the 17626 genes detected (at least one UMI in at least six cells) in the merged embryo-L1 dataset. The background gene set for genes from the L1 vs L4 comparison consisted of the 18330 genes detected in the merged L1-L4 neuronal datasets. We used Bonferroni corrected p-values and calculated a fold-enrichment from the background set using the formula^140^:

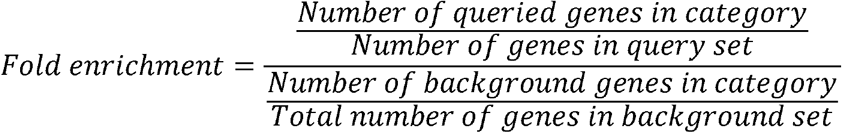

#### Gene Regulatory Network Analysis

We used the *Cel*EsT gene regulatory network (GRN)^53^ to estimate TF activity from the differential expression analyses between the embryo and L1 stages and between the L1 and L4 stages. This resource constructed networks consisting of a TF and its predicted targets for 487 TFs. The TF-predicted target relationships were determined from a combination of DNA-binding motif analysis, ChIP-seq and enhanced yeast one-hybrid (eY1H) screens, with an average of 829 targets per TF. The rationale behind this approach is that the activity of a TF can be inferred from the collective changes in the expression of that TF’s targets. For each neuron in each age comparison, the average log2 fold-change for all genes was used as input with the *Cel*EsT GRN, which contains data for 487 TFs, for the decouple function in the decoupleR R package^141^. We used the multivariate linear model (‘mlm’). We calculated Benjamini-Hochberg corrected p-values to control for multiple comparisons within each neuron. We retained only those TFs with adjusted p-values < 0.05. We also filtered our results to retain TFs that were detected in the neuron of interest in at least one of the ages of the comparison.

#### Thresholding

To distinguish true gene expression from noise, we used a thresholding procedure^16^ on pseudobulk aggregated data for each neuronal cell type based on the proportion of cells in each neuron type in which a gene was detected. We performed this thresholding on the reannotated L4 data and the L1 dataset separately. We first calculated aggregate statistics for each neuron type, including the proportion of cells expressing each gene and a normalized TPM expression value^22^. This TPM value is calculated by normalizing UMI counts for each cell by a size factor (the monocle3 software package generates this as the UMI count for each cell divided by the geometric mean of total UMI counts for all cells), taking the mean normalized expression of each gene across the single cells corresponding to a cell type, and rescaling the resulting values to sum to 1,000,000^22^.

In exploratory analyses, we tested thresholding gene expression based on several criteria, including hard cutoffs of either UMI counts or the proportion of cells in which a gene was detected, as well as dynamic thresholding that considered each gene independently. We compared each thresholded dataset to a ground truth dataset^16^ consisting of 160 genes with known expression patterns across the nervous system. We generated an L1 ground truth dataset (Table S16) which differed slightly from the previously used L4 ground truth dataset due to developmental changes in innexin expression as described^142^. The L4 ground truth was the same as previously used. For a wide range of threshold values (272 possible thresholds, ranging from 0 – 1.0), we generated 5,000 stratified bootstraps of the ground truth genes using the R package boot^143,144^, and computed true positive (TPR), false positive (FPR) and false discovery rates (FDR). We estimated 95% confidence intervals with the adjusted percentile (BCa) method. We used the dynamic thresholding procedure, based on the highest proportion of cells expressing each individual gene, since it performed best based on TPR, FPR and FDR when compared to the ground truth. This approach results in proportion threshold values that are calculated for each individual gene^16^.

For each gene, we determined the cell type which had the highest proportion of cells expressing that gene. Thresholds were set as fractions of the highest proportion for each gene. We performed this thresholding for the L1 dataset using the L1 ground truth and generated updated thresholded datasets for the reannotated L4 data using the L4 ground truth. For the L4 data, we used threshold values of 0.02, 0.04, 0.09 and 0.15 as previously described^16^. We selected four threshold levels of varying stringency for the L1 data, at 0.025, 0.04, 0.08, and 0.13 *of the highest proportion* of cells in which a gene was detected. These thresholds were selected to match the TPR, FPR, and FDR values from the L4 data (Table S17).

At each dynamic threshold level, for each gene the TPM value was set to 0 for each neuron type in which the proportion of cells expressing the gene was below the dynamic threshold for that gene. For example, in the L1 data, the VA1 neuron cluster had highest proportion of cells with transcripts for the TF *unc-4*, at 90.7% of the cells, giving proportion thresholds of 0.02265, 0.03624, 0.07248, and 0.11778. Thus, at the lowest stringency (threshold 1), the *unc-4* TPM value was set to 0 in all cell types in which it was detected in fewer than 2.265% of the cells, whereas at the highest stringency (threshold 4), *unc-4* TPM value was set to 0 in all neuron types in which it was detected in fewer than 11.778% of cells. For the posterior Hox gene *php-3*, the neuron class PHC had the highest proportion of positive cells, with 42.3% of PHC cells having *php-3* transcripts. Proportion thresholds for *php-3* were 0.0106 (i.e., 0.025 * 0.423), 0.01692, 0.03384 and 0.05499. In the case of *php-3*, at threshold 1, all neuron types in which *php-3* was detected in fewer than 1.06% of the cells were set to 0, and at threshold 4, all neuron types in which *php-3* was detected in fewer than 5.499% of cells were set to 0. For all neuron classes in which the proportion of cells expressing a gene was above the threshold, the continuous TPM values were retained.

Thresholds and stats for each age:

L4 data

| Threshold | Fraction of highest proportion | TPR | FPR | FDR |
|---|---|---|---|---|
| 1 | 0.02 | 0.853 | 0.118 | 0.197 |
| 2 | 0.04 | 0.810 | 0.0880 | 0.145 |
| 3 | 0.09 | 0.695 | 0.0458 | 0.105 |
| 4 | 0.15 | 0.578 | 0.0301 | 0.085 |

L1 data

| Threshold | Fraction of highest proportion | TPR | FPR | FDR |
|---|---|---|---|---|
| 1 | 0.025 | 0.853 | 0.120 | 0.202 |
| 2 | 0.04 | 0.807 | 0.092 | 0.170 |
| 3 | 0.08 | 0.689 | 0.055 | 0.127 |
| 4 | 0.13 | 0.577 | 0.037 | 0.103 |

Our ground truth dataset contained several genes with independently documented expression in all neuron classes as well as several genes not expressed in any neuron class. The dynamic thresholding procedure did not perform well compared to ground truth for these two categories of genes, as the dynamic approach assumes that a gene is expressed in some cell classes and absent in others. We therefore set two initial filters that precluded some genes from the dynamic thresholding. First, we retained the TPM values in all cells for genes that were detected in ≥1 % of the cells in every neuron class. 474 genes met this criterion in the L1 dataset, and 183 genes met this criterion in the L4 dataset. Second, genes that were detected in ≤2% of the cells in every neuron class were removed from the thresholded neuronal data; 5,345 genes met this criterion in the L1 dataset, and 4,109 genes met this criterion in the L4 dataset. Many of these genes were detected in much higher proportions in non-neuronal cells. The raw detected expression values of these genes in neuron classes can be found in the unthresholded datasets. The two hard thresholds of detection in ≥ 1% of all cells in every neuron class and detection in ≤ 2% of cell in every neuron class were selected empirically based on the results of comparing thresholding with several possible threshold values to the ground truth. We confirmed that no genes in our L1 and L4 data sets met the critierion of being detected in ≥1% but ≤2% of the cells in every neuron class.

We also performed dynamic thresholding for the newly annotated embryonic data. We used the same thresholds as for the L1 data because we lack an equivalent ground truth dataset for the embryonic nervous system for calculating independent TPR, FPR and FDR values.

#### L1 stability

We calculated the stability of gene expression from L1 to L4 using binarized expression data from threshold 2. We assessed the stability of expression for each gene as follows:

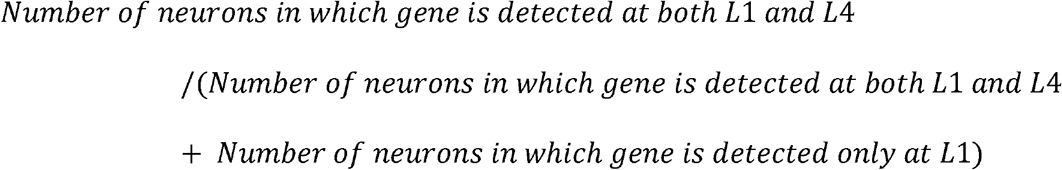

To account for the appearance of expression at L4 as well, we also calculated the fraction of neurons expressing a gene at L4 in which the gene was not detected at L1. Genes were grouped into functional gene families based on the literature, known function and sequence homology^83^.

#### Microscopy

For confocal imaging, worms were mounted on slides containing agarose pads (2-5%) and immobilized with levamisole (17 mM). Images were acquired on a Nikon A1R or Olympus FV1000 laser scanning confocal microscope, with 40X or 60X oil immersion objectives.

##### Validation of differential expression between L1 and L4

For imaging analysis of neuropeptide reporters in the L1 and L4 stage, well-fed, gravid adults were treated with a hypochlorite solution. Between 20 to 30 eggs were transferred to a standard OP50 seeded plate. Animals were bleached and eggs seeded 19h before imaging for L1 stage and 48h before imaging for the L4 stage. In some experiments, L1 and L4 worms identified by morphology and picked off mixed stage, well-fed plates. Worms were immobilized with 100 mM sodium azide (NaN_3_) and placed on 5% agarose pads on glass slides. Imaging was conducted at the L1 or L4 stage. Images were captured using a wide-filed microscope (Axio Imager Z2) at 40X magnification. Image processing and the fluorescence intensity measurements were carried out using Fiji^145^. We combined CRISPR/Cas9 engineered reporter alleles for neuropeptide genes with the NeuroPAL landmark strain *(otIs669)* to determine in which neuron types they are expressed^146^. Neuron types were identified by the NeuroPAL colour code and the location of their nuclei compared to other neurons. Comprehensive guides for identifying neurons using NeuroPAL are available at https://www.hobertlab.org/neuropal/. In cases in which NeuroPAL was not used for neural ID, the expression of the reporter in a neuron was determined by assessing the presence of “speckles” in the cell of interest^147^. Anatomical position within a specific ganglion was used to further specify the final ID.

##### In vivo quantification of ribosomal protein expression

Ribosomal protein encoding genes *rps-4, rpl-7A* and *rpl-25.1* were tagged endogenously with *C. elegans* codon optimized mEosEM at their C-terminus by CRISPR/Cas9 genome editing as described^148^. Briefly, single stranded repair templates encoding mEosEM flanked with 40 bp homology arms were co-injected with pre-complexed Cas9 RNP targeting the respective locus. Worms at the L1 or L4 stage were immobilized with 50mM sodium azide on 5% agarose pads and imaged on a Zeiss LSM980 laser scanning confocal microscope (40x objective) with the 488nm laser and full spectral range of the GaAsP-PMT detector at 800V for fluorescence and transmitted light PMT detector for Nomarski co-imaging. Regions corresponding to indicated cell types were outlined and measurements were restricted to cytoplasmic areas using the “threshold” function in Fiji before pixel intensity was quantified.

#### Neuropeptide Network analysis

##### Neuropeptide network construction, with no spatial filtering

The neuropeptide networks were constructed as previously described^72^. Briefly, we considered a directed connection between neurons i and j whenever neuron i expressed a neuropeptide precursor gene (NPP) and neuron j expressed the cognate G-protein coupled receptor (GPCR). Spatial constraints on peptidergic signaling were incorporated subsequently (see next section). We used a published dataset of neuropeptide-GPCR interactions^71^ as the source of peptide-receptor cognate pairs.

To identify neurons expressing various neuropeptides and receptors, we used the expression data presented in this article, which includes 45 NPPs and 47 GPCR genes encoding neuropeptide receptors that are expressed in both L1 and L4 larvae. These correspond to 88 validated neuropeptide-receptor (NPP-GPCR) biochemical interactions. For each of the neuropeptide-receptor pairs (n), we defined an adjacency matrix A^(n)^ whose entries A^(n)^(i, j) equal 1 if neuron i expresses the relevant NPP and neuron j expresses the corresponding GPCR, and 0 otherwise. These adjacency matrices were built using a binarized, thresholded version of the expression data for the 291 neurons shared between the L1 and L4 larval stages. Two different gene expression thresholds were used as shown in Figure S14, with both thresholds yielding similar results. The L4 network included 49 NPP genes and 51 GPCRs^16,71,72^. The diffusion range was established by defining a matrix of neuronal proximity, in which we used electron microscopy (EM) data of the L4 nervous system to assign neurons to specific process bundles^24^. The L1 network included 45 NPPs and 47 GPCRs expressed in both the L1 and L4 stages, forming 88 validated ligand-receptor pairs with high-potency interactions (EC_50_ ≤ 500 nM); transcripts for the neuropeptides *flp-23*, *nlp-22*, *nlp-27*, *nlp-25* and the receptors *npr-41, dmsr-16, gnrr-3,* and *dmsr-11* were detected in the nervous system at the L4 but not at the L1 stage. We used neuronal proximity information from EM data from L1 animals 16 hours post-hatching to define diffusion range^6,9^ (Figure S13A-B). We made model networks based on short-range diffusion, in which neuropeptide signaling occurs only between neurons with processes in the same process bundle (∼10nm), and mid-range diffusion, which allows signaling between neuron processes in the same anatomical area (head, midbody, tail) (<50nm).

As described above, each NPP-GPCR interaction defines a separate binary network. The overall neuropeptide adjacency matrix, A, was obtained by summing these individual interaction networks, resulting in a weighted directed network in which the weight of each edge represents the number of distinct neuropeptide-receptor interactions connecting the corresponding pair of neurons.

For the across-development network analysis, we constructed several separate networks, as illustrated in Figure 6A. The first network contains all binary connections at L1, whereas the second network contains all binary connections at the L4 larval stage. The third network contains only binary connections that are present at both L1 and L4 developmental stages. Finally, the fourth network contains only those connections that are conserved between L1 and L4 and are supported by the identical combination of NPP-GPCR interactions in both stages.

##### Neuropeptide network spatial constraining

Neuropeptide signaling was locally thresholded to filter out connections between neurons anatomically distant from each other. A developmental-stage dependent matrix of anatomical proximity was constructed using electron microscopy data of the nervous system of the *C. elegans* L1 larvae (16h after hatching)^6,9,84^ for the L1 matrix, and separately using electron microscopy data of the nervous system of *C.elegans* at L4 larvae (as shown in L.Ripoll-Sánchez et al. 2023) for the L4 matrix. These data were used to create a table of locations for each neuronal process, identifying 27 different neuronal process bundles in the *Caenorhabditis* nervous system as previously defined^26^. This classification was then used to filter out neuropeptidergic connections based on putative signaling ranges in the two developmental stages. The mid-range stringency thresholding used in this analysis allows connections between neurons with neuronal processes in the same anatomical area (∼50 μm distance): head (including pharynx and the ventral cord neurons in the ventral ganglion), midbody and tail. The differences in size between both developmental stages imply that the mid-range, short-range and long-range distances were different, but the area they connected is consistent for both developmental stages.

##### Degree

Edge counts and adjacency matrices were all computed using binary, directed versions of the networks. These networks were used to compute degree using the Brain Connectivity Toolbox^149^ for MATLAB. Degree is defined as the number of edges connected to a given node. In-degree is the number of incoming connections to a given node and out-degree is the number of outgoing connections.

##### Identifying mesoscale organization

To identify mesoscale organization in the neuropeptide network, we followed our previous approach^72^. For each developmental stage, each neuron was represented by its incoming connectivity profile, corresponding to a column of the weighted adjacency matrix A. Thus, while the mathematical object A remains the weighted adjacency matrix of the neuropeptide network, it is interpreted here as a collection of neuronal connectivity profiles, with each column representing a high-dimensional feature vector describing the incoming neuropeptide connections of a single neuron. These high-dimensional connectivity profiles were first standardized (z-score normalization, mean =0, standard deviation =1). We then performed, in parallel, principal component analysis (PCA) and dimensionality reduction using t-distributed stochastic neighbor embedding (t-SNE) on the standardized connectivity profiles.

We identified clearly distinct groupings of neurons in both PCA and t-SNE embeddings (Figure S17A-B, D-E). Neurons were then assigned to candidate clusters based on their positions in the embedded space. For t-SNE, the cluster assignments remained consistent across multiple t-SNE iterations. Similarly to t-SNE, neurons were visually assigned to candidate clusters based on their positions in the PCA embedding. Candidate groupings from t-SNE were cross-checked with groupings based on PCA. Most group assignments were consistent across t-SNE and PCA clustering; ambiguities were resolved by comparing t-SNE to PCA embeddings. Finally, the rows and columns of the adjacency matrix were reordered according to the identified clusters, revealing block structure in the reordered adjacency matrix and facilitating visualization of mesoscale organization in the neuropeptide network.

As a further independent validation of the mesoscale groupings, a probabilistic block model was fit directly to each neuron’s incoming connectivity profile. Binary incoming connections were weighted using an inverse document frequency (IDF)-style transform:

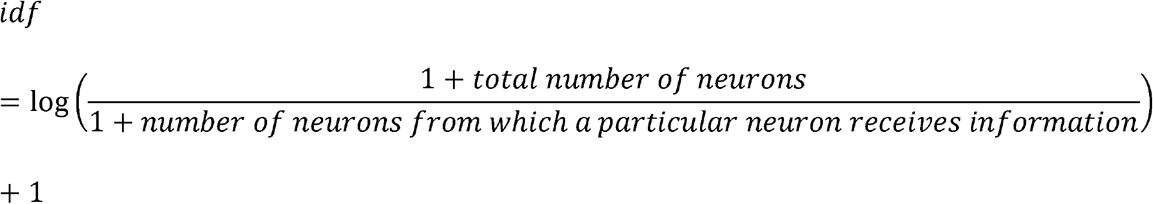

to down-weight inputs from broadly-connecting source neurons, and the resulting profiles were L2-normalized and projected into a low-dimensional space using truncated singular value decomposition (n_components = 12 for L4, n_components = 8 for L1). A diagonal-covariance Gaussian mixture model was then fit to this embedding (k = 4 components, n_init = 10 for L4, n_init = 15 for L1, random_state = 42), assigning each neuron to one of k blocks based on maximum posterior probability, with this probability retained as a measure of assignment confidence. Block assignments were compared, neuron by neuron, against the groups identified from the t-SNE and PCA embeddings.

The block model and the t-SNE/PCA approaches gave the same group assignments for every neuron in L4. For the L1 dataset, three neurons (PQR, ALNL, ALNR) were grouped in the periphery in the block model rather than the motor (PQR) or sensory (ALNL, ALNR) groups assigned by t-SNE and PCA methods. Visual inspection of the incoming connectivity profiles of these neurons clearly corresponded to the motor and sensory groups respectively, supporting the t-SNE and PCA group assignments (Fig. 6; Figure S17C, F). This discrepancy reflects the sparser, less redundant connectivity of the L1 network relative to L4: with fewer informative connections per neuron at this earlier developmental stage, the dimensionality-reduction step underlying the block model is more susceptible to attenuating subtle, group-defining connectivity signals for a small subset of neurons.

##### Quantification and statistical analysis

Statistical data is reported in the main text, figures, and tables as noted. Significance adheres to the common standard, after adjusting for multiple testing, of p < 0.05. The symbols *, **, ***, **** refer to p < 0.05, 0.01, 0.001, and 0.0001, respectively. Not significant is described as n.s. The n for each statistical test is described in each figure.

#### Connectome comparison

##### Standardizing chemical synaptic and membrane contact matrices

The EM data of larval nerve rings^6^ provide resolution at the level of single individual cells (e.g., ASIL and ASIR). To match the resolution of the scRNA-seq expression data, we summed the number of synaptic connections and the physical membrane contacts of the neurons within each transcriptionally defined class (ASIL + ASIR for the ASI neuron). We did this independently for larval datasets from late L1 through adult (datasets 4 – 8 and series JSH and N2U in source publications)^6,150^. To account for the variability between animals, we focused on connections present in multiple datasets. For the L1 dataset, we used the summed synapse counts and membrane contact data from dataset 4 (∼16 hours after birth, in late L1) but only kept those synapses and membrane contacts that were also present in dataset 5 (∼23 hours after birth, L2 stage).

For the L4 comparison, we used the numbers from dataset 8 (∼50 hours after birth) but only kept synapses that were also present in dataset 7, and both the JSH and N2U reconstructions^150^ and membrane contacts that were also present in the JSH and N2U series (membrane contact data are not available for dataset 7^6^).

We generated a list of 443 genes that encode proteins with extracellular domains from several published resources^81,83,85^. These include genes encoding transmembrane cell surface proteins, predicted GPI-anchored proteins and secreted proteins. 298 of these were detected in nerve ring neurons at L1. 313 of these were detected in nerve ring neurons in the updated L4 data. Binding data for extracellular domains was taken from Table S3 in Nawrocka, et al^85^ and from Wormbase and the Alliance for Genome Resources^151^. We used the SimpleMine tool in the Alliance for Genome Resources website to search for curated Protein-Protein interactions between the genes in our CAM list. We then used the Wormbase interaction data to identify publications supporting these interactions and manually curated interactions that included the extracellular domains to exclude binding of intracellular regions.

##### Membrane contact enrichment

For each neuron (‘tested neuron’) with contacts in the nerve ring (80 neurons at L1, 84 at L4), we separated all other neurons into two categories based on whether they had any membrane contact with the tested neuron. We performed unpaired two-sample Welch’s t-tests (assuming unequal variances) in R on the mean expression of each gene in contacted neurons versus the mean expression in non-contacted neurons. We used the threshold 2 TPM expression values for this analysis. We also generated a t-statistic representing the magnitude and direction of enrichment (positive values indicate enrichment in contacted cells, negative values indicate enrichment in non-contacted cells), p-value, the fold change of the means and log2 transformed fold changes as a measure of effect size. In calculating log2 fold changes, we added a value of ‘1e-5’ to the mean of each group to prevent undefined values when one mean was equal to zero. We generated adjusted p-values for multiple comparisons across genes in each neuron using the “p.adjust” function in R with the Benjamini-Hochberg method. We also determined for each gene whether any known binding partners were expressed in the tested neuron.

##### Synaptic partner gene enrichment

For each tested neuron, we split all other cells having membrane contact with the tested neuron into one of two categories: 1) synaptic partners or 2) non-synaptic, but adjacent cells. We performed this operation separately for presynaptic inputs and postsynaptic outputs for each tested neuron. We restricted this analysis to neurons that had at least three presynaptic inputs or postsynaptic outputs. We used the threshold 2 TPM normalized expression values for this analysis. We performed unpaired Welch’s t-tests on the mean expression of each gene in the synaptically-connected neurons versus the non-synaptic, adjacent neurons. We generated fold change, log2 transformed log fold change, t-statistics, p-values and Benjamini-Hochberg adjusted p-values for each tested gene/tested neuron combination. We then assessed whether a tested gene had any known binding partners in the tested neuron. We performed this analysis separately for the data from L1 and L4.

### QUANTIFICATION AND STATISTICAL ANALYSIS

The statistical analyses performed are described in the relevant Method details sections of the STAR Methods for each analysis. They are also summarized in the respective figure legends. In the case of single-cell RNA sequencing analyses, n represents individual single cells. For microscopic analyses, n represents individual animals. Statistical analyses were performed in R.

### ADDITIONAL RESOURCES

Resources are listed under the Data and code availability section of the results.

