## Supplemental Figures for "A gene expression atlas of a juvenile nervous system"

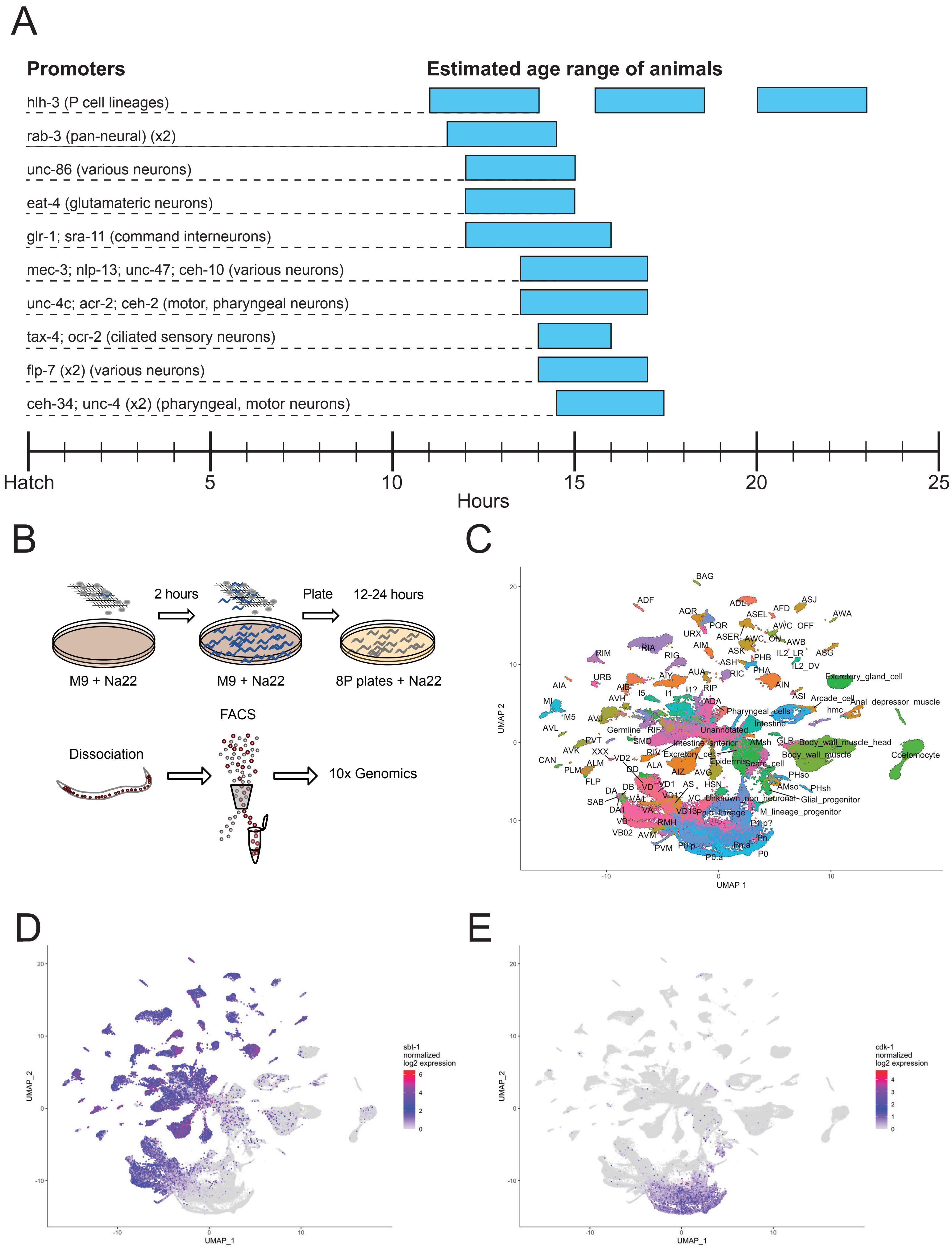
A
Promoters
Estimated age range of animals
hlh-3 (P cell lineages)
rab-3 (pan-neural) (x2)
unc-86 (various neurons)
eat-4 (glutamateric neurons)
glr-1; sra-11 (command interneurons)
mec-3; nlp-13; unc-47; ceh-10 (various neurons)
unc-4c; acr-2; ceh-2 (motor, pharyngeal neurons)
tax-4; ocr-2 (ciliated sensory neurons)
flp-7 (x2) (various neurons)
ceh-34; unc-4 (x2) (pharyngeal, motor neurons)
Hatch
5
10
15
20
25
Hours
B
2 hours
Plate
12-24 hours
M9 + Na22
M9 + Na22
8P plates + Na22
FACS
Dissociation
10x Genomics
C
BAG
ADF
ADL
ASJ
AQR
AFD
AWA
ASEL
PQR
AWC_OFF
URX
ASER
AWC_ON
AWB
AIM
RIA
ASK
IL2_LR
ASG
RIM
RIG
PHB
ASH
IL2_DV
AUA
RIC
PHA
AIY
Excretory_gland_cell
URB
AIA
AIB
I2
I1?
RIP
AIN
AVH
I5
I1
ASI
Arcade_cell
MI
ADA
Anal_depressor_muscle
M5
Pharyngeal_cells
hmc
AVJ
Intestine
AVL
Germline
RIF
Unannotated
PVT
SMD
Intestine_anterior
AMsh
GLR
Body_wall_muscle_head
AVK
XXX
RIV
Excretory_cell
Body_wall_muscle
Coelomocyte
CAN
VD2
Epidermis
Seam_cell
PLM
ALM
ALA
DD
AIZ
AVG
PHso
FLP
VD
VD1
AS
HSN
AMso
PHsh
DA
DB
VD12
VC
Unknown_non_neuronal
Glial_progenitor
SAB
VA1
DA1
VA
VD13
Pn.p_lineage
M_lineage_progenitor
VB
RMH
P1.p?
VB02
AVM
Pn
PVM
P0.p
Pn.a
P0.a
P0
UMAP 2
UMAP 1
D
UMAP_2
UMAP_1
sbt-1
normalized
log2 expression
E
UMAP_2
UMAP_1
cdk-1
normalized
log2 expression

**Figure S1. Single-cell RNA sequencing of L1 larvae, related to Figure 1.** A) Neuron-specific fluorescent reporter strains and developmental ages sampled for scRNA-Seq. The rab-3, flp-7 and ceh-34; unc-4 reporter strains were used for two experimental samples each. B) Schematic of mesh protocol used to generate large synchronous cultures (top) and use of FACS to enrich for targeted cell populations (bottom). C-E) UMAPs of all 161,562 cells, colored by C) cell type annotation, D) expression of the pan-neural gene sbt-1 marking post-mitotic neurons, and E) expression of the cell cycle gene cdk-1 marking progenitor cells.

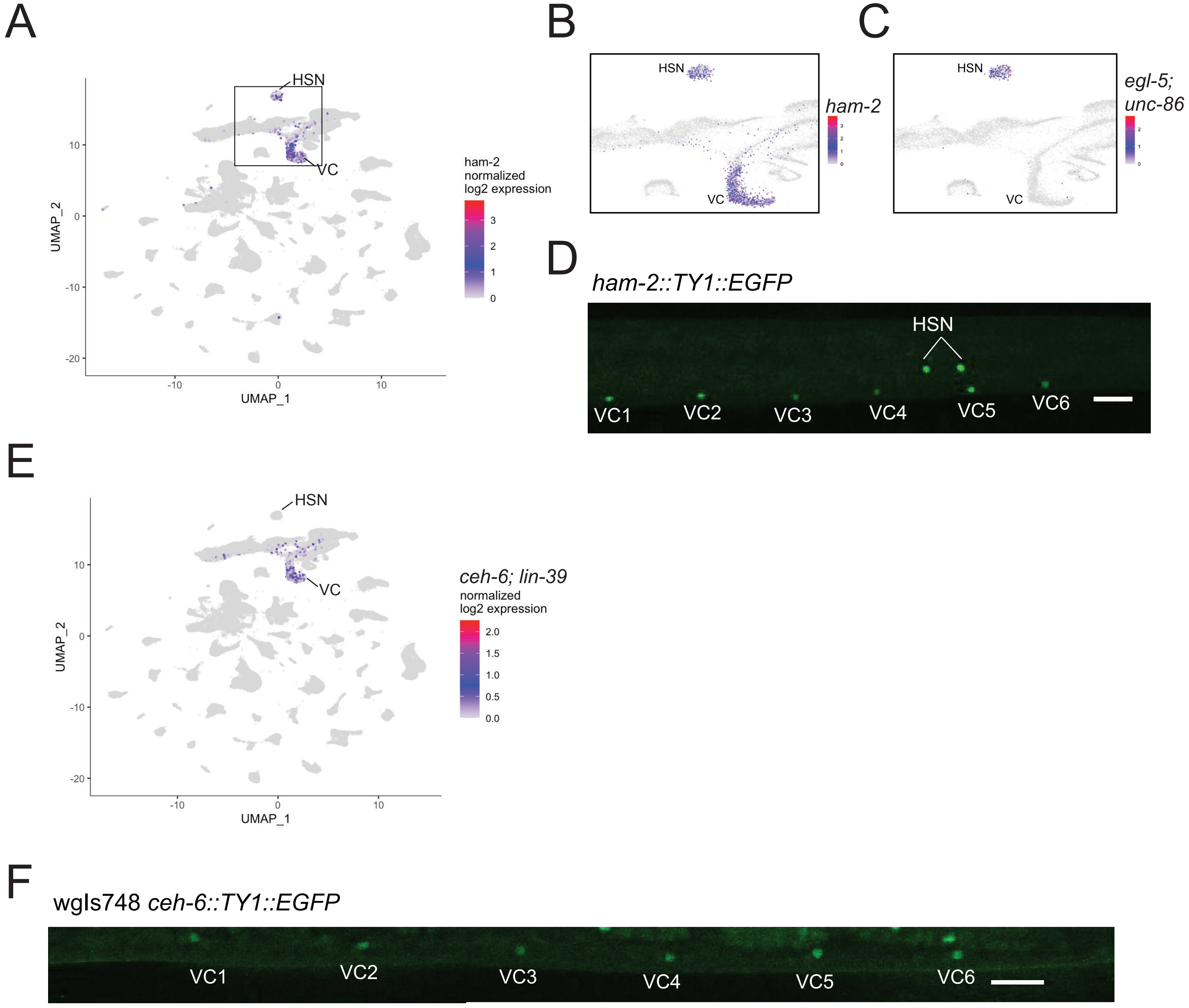


**Figure S2. Annotation of HSN and VC neurons in L1, related to Figure 1.** A) UMAP of ham-2 expression, which is restricted to HSN and VC. B-C) Enlargement of the boxed region in panel A, colored by B) expression of the TF ham-2 and C) co-expression of egl-5 and unc-86, which uniquely marks HSN. D) Confocal micrograph of a ham-2 fosmid reporter in late L1 larvae shows expression in VC neurons in the ventral cord and in the HSN neuron pair. E) The VC neuron cluster uniquely shows co-expression of the TFs ceh-6 and lin-39. F) Confocal micrograph of a ceh-6 fosmid reporter in late L1. Expression is restricted to the six VC neurons in the ventral nerve cord. Scale bars = 10 μm.

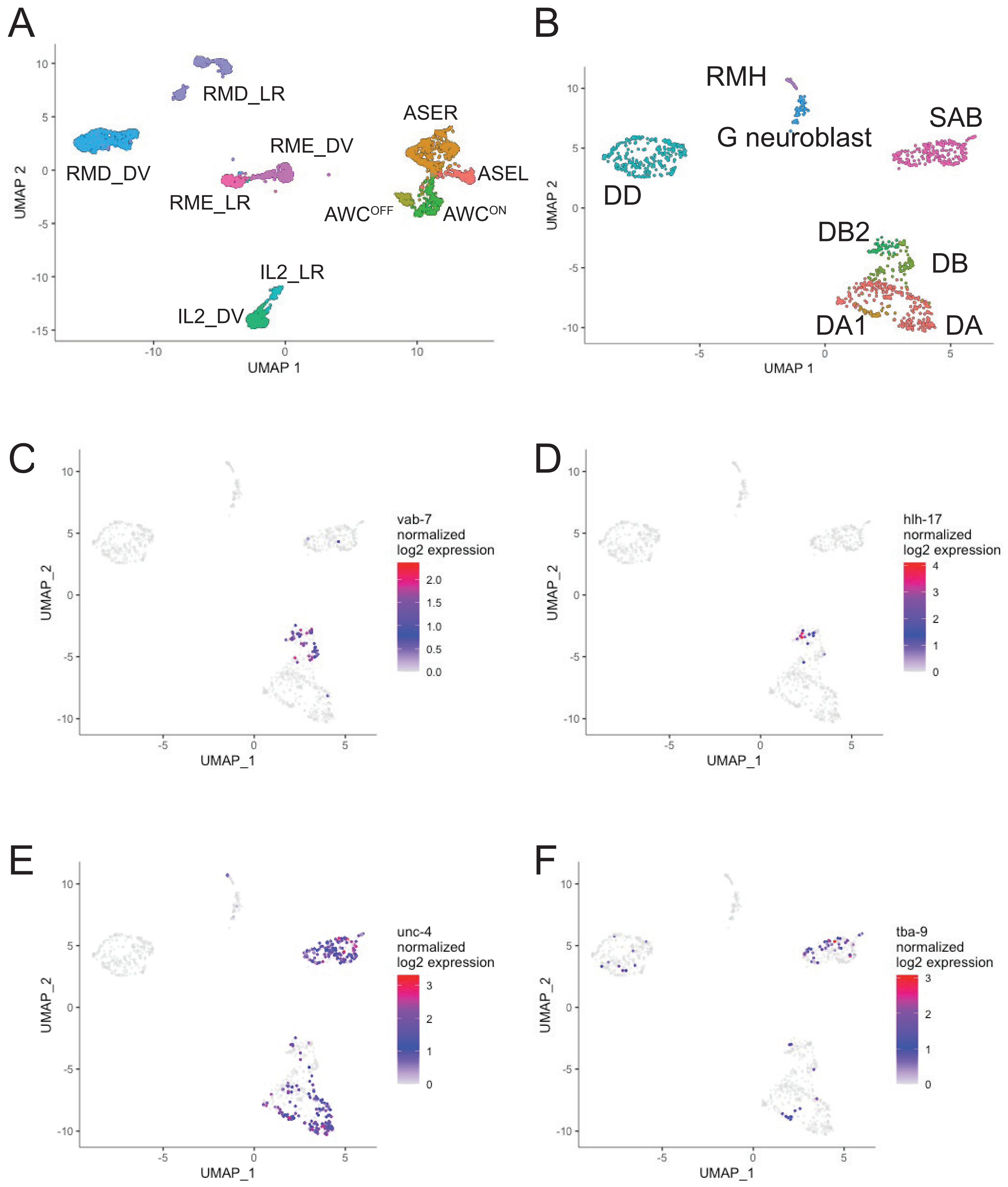
A
RMD_LR
RMD_DV
RME_DV
RME_LR
ASER
ASEL
AWC$^{OFF}$
AWC$^{ON}$
IL2_LR
IL2_DV
UMAP 1
UMAP 2
B
RMH
G neuroblast
SAB
DD
DB2
DB
DA1
DA
C
vab-7 normalized log2 expression
D
hlh-17 normalized log2 expression
E
unc-4 normalized log2 expression
F
tba-9 normalized log2 expression
UMAP_1
UMAP_2

**Figure S3. Neuronal subclasses are present in the L1, related to Figure 1.** A) Sub-UMAP showing separation of neuronal subclasses for RMD, RME, IL2, ASE, and AWC. B) Sub-UMAP showing embryonic motor neurons that include the newborn RMH cluster, the G neuroblast, DD, SAB, and subclasses of the DA and DB motor neurons. C) Sub-UMAP showing expression of the DB class marker vab-7. D) Sub-UMAP showing expression of the DB2-specific marker hlh-17. E) Expression of the DA and SAB class marker unc-4. F) Expression of the tubulin tba-9, which is restricted to DA1 among DA neurons26.

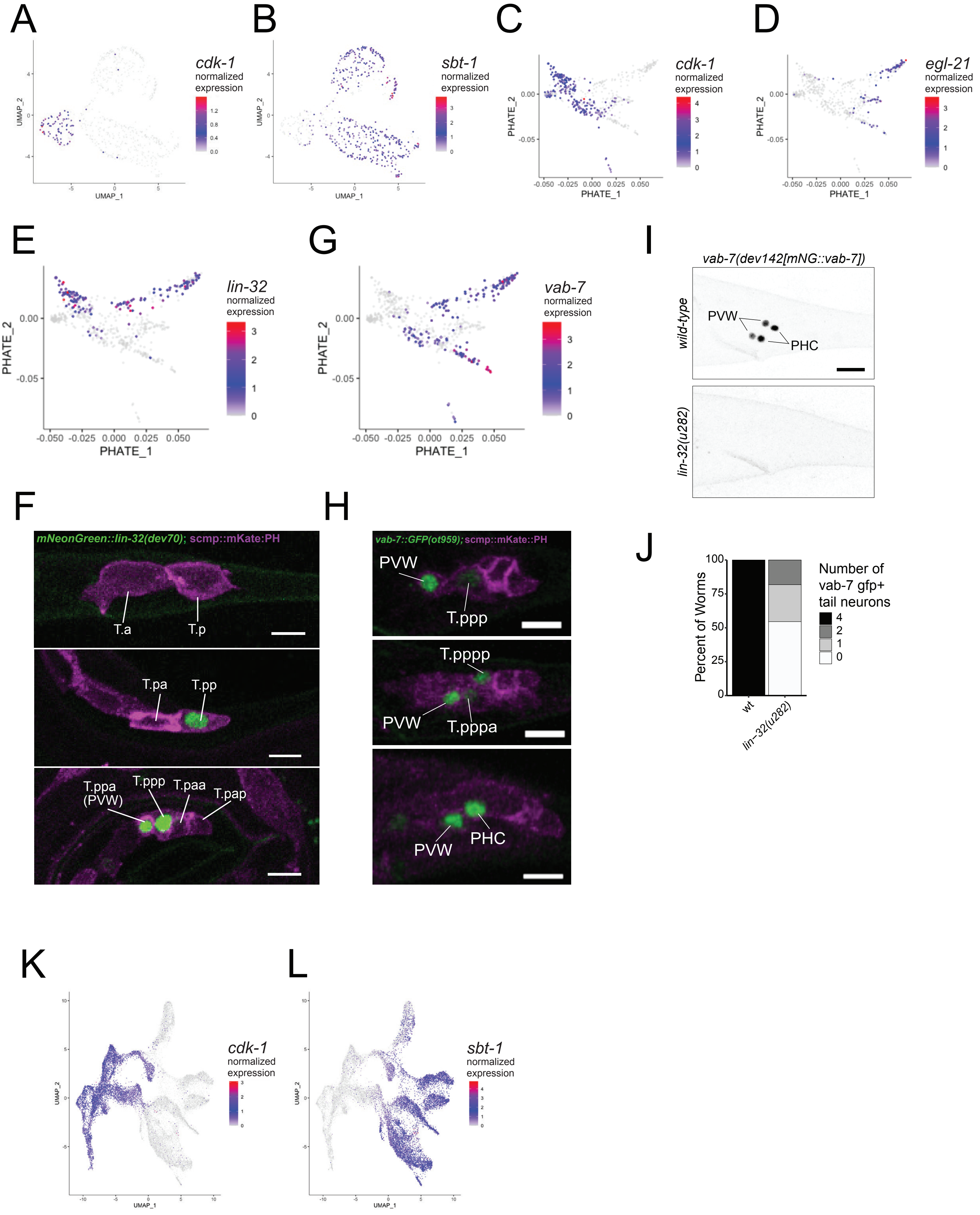
A
cdk-1 normalized expression
UMAP_1
UMAP_2
B
sbt-1 normalized expression
C
cdk-1 normalized expression
PHATE_1
PHATE_2
D
egl-21 normalized expression
E
lin-32 normalized expression
G
vab-7 normalized expression
I
vab-7(dev142[mNG::vab-7])
wild-type
PVW
PHC
lin-32(u282)
F
mNeonGreen::lin-32(dev70); scmp::mKate:PH
T.a
T.p
T.pa
T.pp
T.ppa (PVW)
T.ppp
T.paa
T.pap
H
vab-7::GFP(ot959); scmp::mKate::PH
PVW
T.ppp
T.pppp
T.pppa
PHC
J
Percent of Worms
Number of vab-7 gfp+ tail neurons
4
2
1
0
wt
lin-32(u282)
K
cdk-1 normalized expression
L
sbt-1 normalized expression

**Figure S4. Q, T and P lineage progenitors and neurons in the L1, related to Figure 2.** A-B) Sub-UMAPs of Q.pa lineage, colored by A) expression of the cell cycle gene cdk-1 marking the Q.pa progenitor and B) the neuropeptide processing gene sbt-1 in post-mitotic Q-derived neurons. C-D) PHATE plots of the T-lineage showing C) expression of cdk-1, which marks neuronal progenitors and D) egl-21 in three branches corresponding to postmitotic neurons. E) PHATE plot showing expression of lin-32 in the T.pp lineages. F) Confocal micrographs of an endogenous mNeonGreen::lin-32 reporter allele and a membrane bound mKate expressed in the T lineage by a seam cell promoter. G) PHATE plot showing expression vab-7 in the T.pp lineages. H) Confocal micrographs showing endogenously tagged vab-7::GFP reporter in combination with membrane bound mKate in the T-lineage. Expression of vab-7 begins in PVW (T.ppa), its sister T.ppp, and the T.ppp descendants, including PHC. I) An endogenous GFP reporter of vab-7 is expressed in the PVW and PHC neuron pairs in the tail in wild-type (top) but not lin-32(u282) animals (bottom). J) vab-7 expression is decreased in PVW and PHC in lin-32 mutants. K) P-lineage UMAP colored by expression of the cell cycle marker cdk-1 and (L) colored by expression of the post-mitotic neuron marker sbt-1. Fisher’s Exact Test, p = 2.243x10-7. Scale bars = 10 μm.

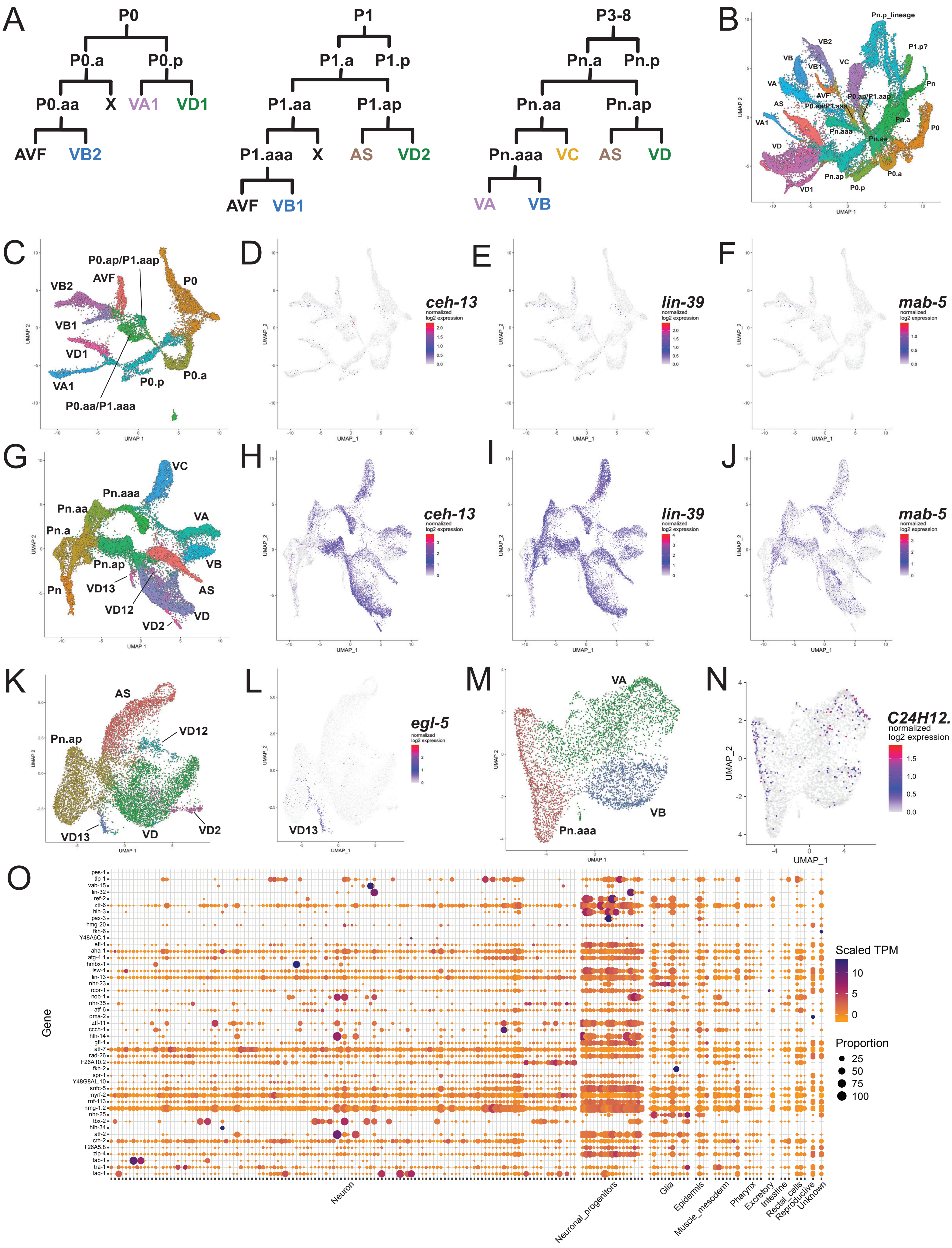

A
P0
P0.a
P0.p
P0.aa
X
VA1
VD1
AVF
VB2
P1
P1.a
P1.p
P1.aa
P1.ap
P1.aaa
AS
VD2
VB1
P3-8
Pn.a
Pn.p
Pn.aa
Pn.ap
Pn.aaa
VC
VD
VA
VB
B
Pn.p_lineage
P1.p?
Pn
P0.ap/P1.aap
P0.aa/P1.aaa
Pn.aaa
Pn.aa
P0
P0.a
P0.p
UMAP 1
UMAP 2
C
D
ceh-13
normalized log2 expression
E
lin-39
F
mab-5
G
VD13
VD12
H
I
J
K
Pn.ap
L
egl-5
M
N
C24H12.1
UMAP_1
UMAP_2
O
Gene
pes-1
tlp-1
vab-15
lin-32
ref-2
ztf-6
hlh-3
pax-3
hmg-20
fkh-6
Y48A6C.1
efl-1
aha-1
atg-4.1
hmbx-1
isw-1
lin-13
nhr-23
rcor-1
nob-1
nhr-35
atf-6
oma-2
ztf-11
ccch-1
hlh-14
gfl-1
atf-7
rad-26
F26A10.2
fkh-2
spr-1
Y48G8AL.10
snfc-5
myrf-2
rnf-113
hmg-1.2
nhr-25
tbx-2
hlh-34
atf-2
crh-2
T26A5.8
zip-4
tab-1
tra-1
lag-1
Scaled TPM
Proportion
25
50
75
100
Neuron
Neuronal_progenitors
Glia
Epidermis
Muscle_mesoderm
Pharynx
Excretory
Intestine
Rectal_cells
Reproductive
Unknown

**Figure S5. Distinguishing P0-P1 and P2-P12 derived progenitors and neurons, related to Figure 2.** A) Diagrams of the P0, P1 and P3-P8 lineages, which adopt related but distinct patterns of division for a subset of cells. B) Sub-UMAP of the P0-P12 lineages, colored by cell type. C-F) Sub-UMAPs of P0-P1 progenitors and neurons, colored by C) cell type, expression of the Hox genes ceh-13 (D),, lin-39 (E), and mab-5 (F). G-J) Sub-UMAPs of P2-P12 progenitors and neurons, colored by cell type (G), expression of the Hox genes ceh-13 (H), lin-39 (I) and mab-5 (J). K) Sub-UMAP of P.ap progenitors, VD neurons, and AS neurons, showing the VD subclasses VD2, VD12, and VD13. L) Sub-UMAP as in panel K, colored by expression of the Hox gene egl-5. M) Sub-UMAP of P.aaa progenitors, VA neurons, and VB neurons, showing lack of distinct VA and VB subclasses. N) Sub-UMAP of P.aaa lineage colored by the expression of the VA2 and VB3 specific marker C24H12.1. O) Dotplot showing the expression of 49 TFs previously assigned to post-embyronic neuronal progenitors across all scRNA-seq cell types.

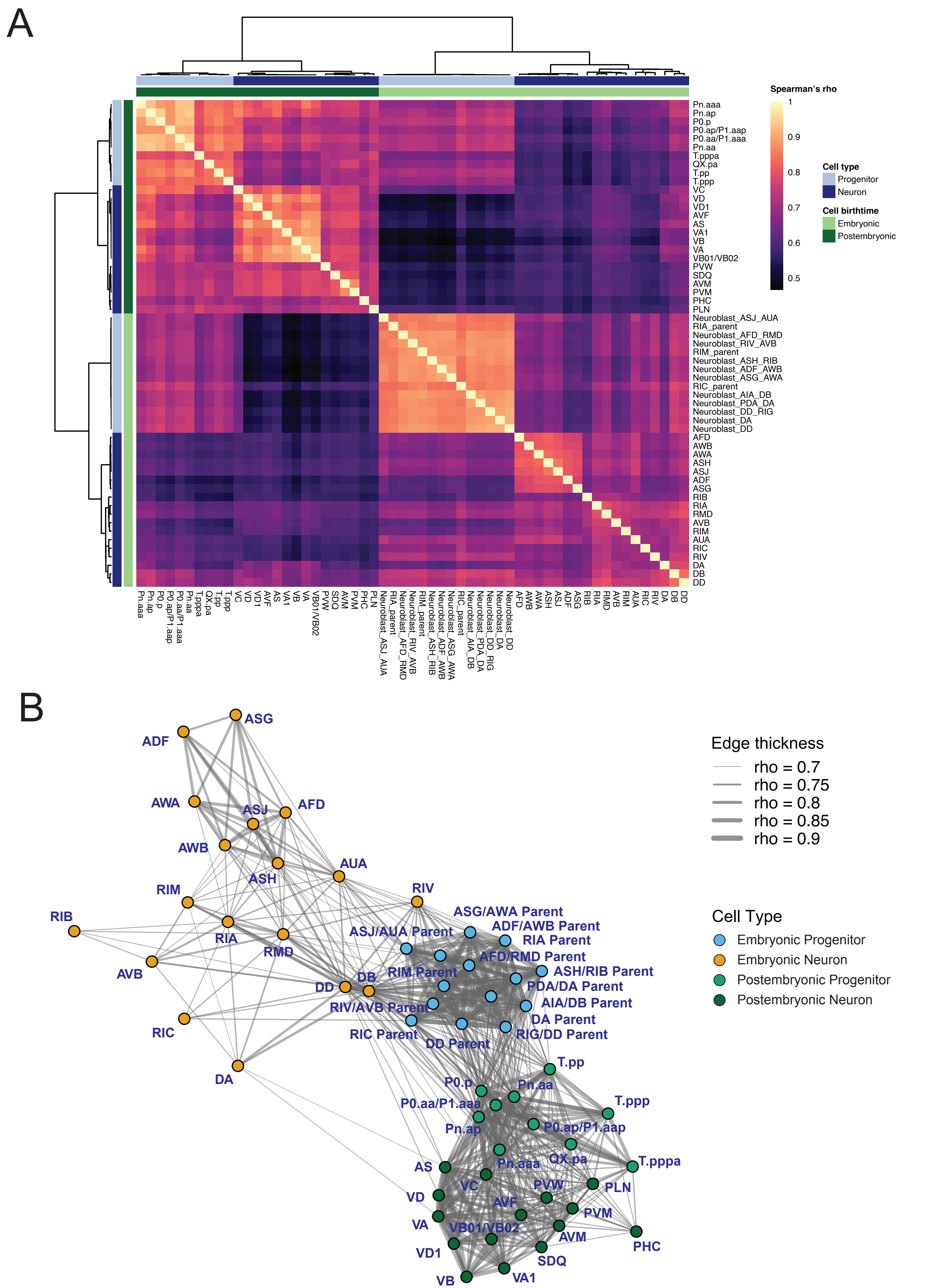

A
Spearman's rho
1
0.9
0.8
0.7
0.6
0.5
Cell type
Progenitor
Neuron
Cell birthtime
Embryonic
Postembryonic
Pn.aaa
Pn.ap
P0.p
P0.ap/P1.aap
P0.aa/P1.aaa
Pn.aa
T.pppa
QX.pa
T.pp
T.ppp
VC
VD
VD1
AVF
AS
VA1
VB
VA
VB01/VB02
PVW
SDQ
AVM
PVM
PHC
PLN
Neuroblast_ASJ_AUA
RIA_parent
Neuroblast_AFD_RMD
Neuroblast_RIV_AVB
RIM_parent
Neuroblast_ASH_RIB
Neuroblast_ADF_AWB
Neuroblast_ASG_AWA
RIC_parent
Neuroblast_AIA_DB
Neuroblast_PDA_DA
Neuroblast_DD_RIG
Neuroblast_DA
Neuroblast_DD
AFD
AWB
AWA
ASH
ASJ
ADF
ASG
RIB
RIA
RMD
AVB
RIM
AUA
RIC
RIV
DA
DB
DD
B
Edge thickness
rho = 0.7
rho = 0.75
rho = 0.8
rho = 0.85
rho = 0.9
Cell Type
Embryonic Progenitor
Embryonic Neuron
Postembryonic Progenitor
Postembryonic Neuron
ASG
ADF
AWA
ASJ
AFD
AWB
ASH
AUA
RIM
RIV
RIB
RIA
RMD
AVB
DD
DB
RIC
DA
ASG/AWA Parent
ADF/AWB Parent
ASJ/AUA Parent
RIA Parent
AFD/RMD Parent
RIM Parent
ASH/RIB Parent
PDA/DA Parent
RIV/AVB Parent
AIA/DB Parent
DA Parent
RIC Parent
DD Parent
RIG/DD Parent
T.pp
P0.p
Pn.aa
P0.aa/P1.aaa
T.ppp
Pn.ap
P0.ap/P1.aap
QX.pa
Pn.aaa
T.pppa
AS
VC
PVW
PLN
VD
AVF
PVM
VA
VB01/VB02
AVM
PHC
VD1
SDQ
VB
VA1

**Figure S6. Transcriptional similarity of neuronal progenitors and newborn neurons between larval and embryonic stages, related to Figure 3.** A) Heatmap of Spearman correlations between transcriptomes of neuronal progenitors and newborn neurons from larval and embryonic stages. Cell types were clustered using the “correlation” distance metric and “ward.D2” clustering method. Developmental stage and progenitor/postmitotic type are denoted by color bars above and to the left of the heatmap. B) Force-directed network of progenitors and newborn neurons. Colors denote developmental stage and progenitor/neuron identity. Edge thicknesses (Spearman’s rho coefficients > 0.7) show the strength of transcriptome similarity between cell types.

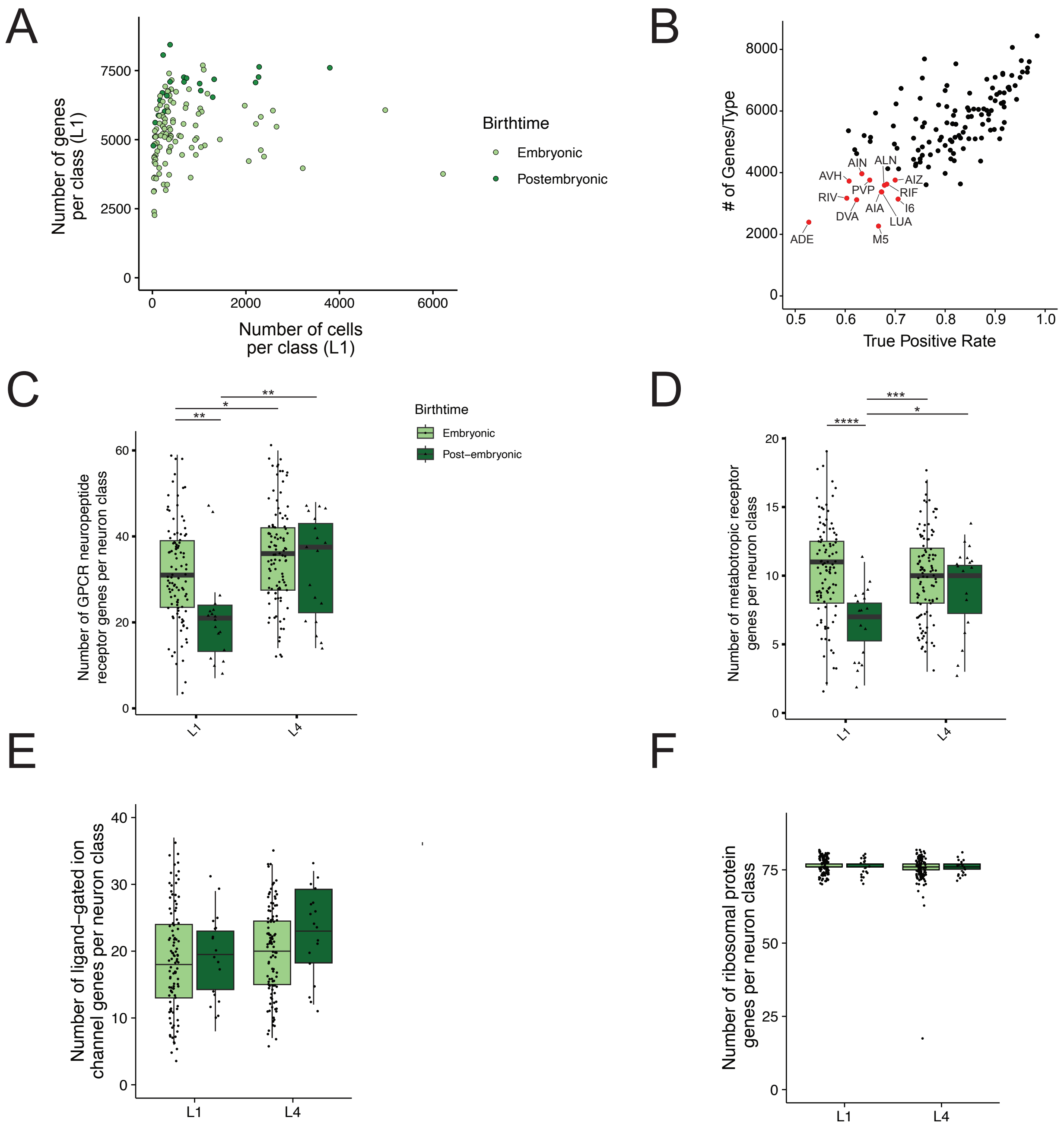
A
Number of genes per class (L1)
Number of cells per class (L1)
Birthtime
Embryonic
Postembryonic
B
### of Genes/Type
True Positive Rate
AIN
ALN
AVH
AIZ
PVP
RIF
RIV
I6
AIA
DVA
LUA
ADE
M5
C
Birthtime
Embryonic
Post−embryonic
Number of GPCR neuropeptide receptor genes per neuron class
L1
L4
D
Number of metabotropic receptor genes per neuron class
L1
L4
E
Number of ligand−gated ion channel genes per neuron class
L1
L4
F
Number of ribosomal protein genes per neuron class
L1
L4

**Figure S7. Expression of selected gene families in the larval nervous system, related to Figure 4.** A) Scatterplot showing the relationship between the aggregate number of genes detected in each neuron class (y axis) using threshold 2 and the number of cells captured per neuron class (x axis) in L1. B) Scatterplot showing true positive rate (TPR, x-axis) and number of genes detected on aggregate (y-axis) for each neuron class from threshold 2. Labeled neuron classes in red show the lowest TPRs and numbers of genes and are more likely to have false negatives. C-F) Combined boxplot and jitter plots showing the number of genes belonging to selected gene families detected per neuron class in L1 (left) and in L4 (right) for embryonic (light green) and postembryonic (dark green) neurons. Gene families: C) GPCR neuropeptide receptors, D) metabotropic neurotransmitter receptors, E) ligand-gated ion channels, and F) ribosomal proteins. All statistical tests were performed using linear models featuring the birthtime, stage, and number of cells per neuron class as covariates. Between group comparisons were performed on the estimated marginal means using the Tukey p-value adjustment for multiple comparisons. * $p$-value $< 0.05$, ** $p$-value $< 0.01$, *** $p$-value $< 0.001$, **** $p$-value $< 0.0001$.

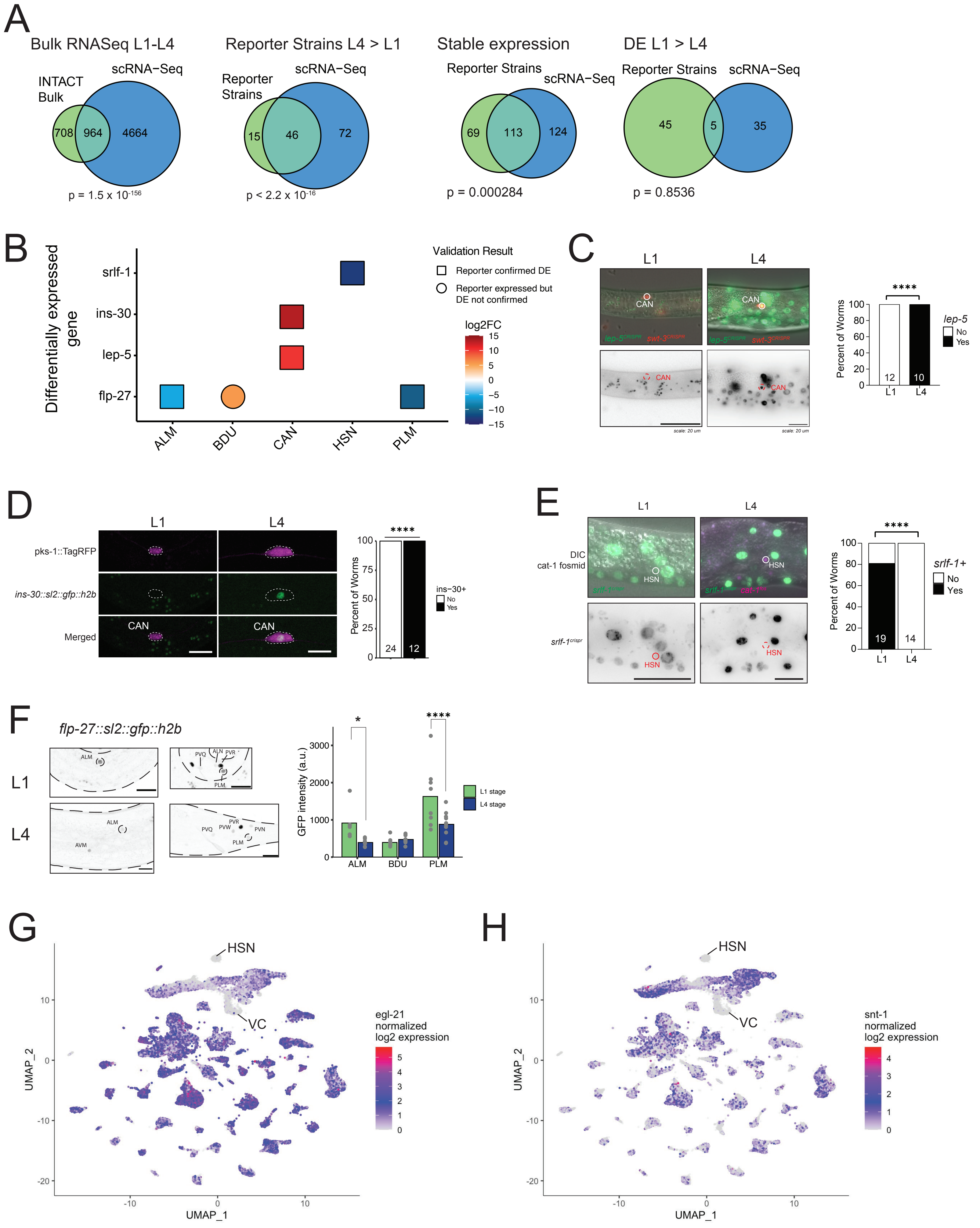
A
Bulk RNASeq L1-L4
INTACT Bulk
scRNA-Seq
708
964
4664
p = 1.5 x 10^-156
Reporter Strains L4 > L1
Reporter Strains
scRNA-Seq
15
46
72
p < 2.2 x 10^-16
Stable expression
Reporter Strains
scRNA-Seq
69
113
124
p = 0.000284
DE L1 > L4
Reporter Strains
scRNA-Seq
45
5
35
p = 0.8536
B
Differentially expressed gene
srlf-1
ins-30
lep-5
flp-27
ALM
BDU
CAN
HSN
PLM
Validation Result
Reporter confirmed DE
Reporter expressed but DE not confirmed
log2FC
15
10
5
0
−5
−10
−15
C
L1
L4
CAN
lep-5^CRISPR swt-3^CRISPR
scale: 20 um
Percent of Worms
lep-5
No
Yes
****
12
10
D
L1
L4
pks-1::TagRFP
ins-30::sl2::gfp::h2b
Merged
CAN
Percent of Worms
****
24
12
ins-30+
No
Yes
E
L1
L4
DIC
cat-1 fosmid
HSN
srlf-1^crispr
Percent of Worms
****
19
14
srlf-1+
No
Yes
F
flp-27::sl2::gfp::h2b
L1
L4
ALM
PVQ
ALN
PVR
PLM
AVM
PVW
PVN
GFP intensity (a.u.)
*
****
ALM
BDU
PLM
L1 stage
L4 stage
G
HSN
VC
UMAP_1
UMAP_2
egl-21
normalized
log2 expression
H
HSN
VC
UMAP_1
UMAP_2
snt-1
normalized
log2 expression

**Figure S8. Validation of differentially expressed genes from scRNA-Seq, related to Figure 5.** A) Venn diagrams showing overlap of differentially expressed genes (DEGs) between L1 and L4 in scRNA-Seq and data from Ref 12. There was significant overlap between scRNA-seq comparisons and bulk neuronal samples using INTACT (left), between scRNA-seq and reporter strain data for cases where expression was higher in L4 than L1 (middle left), and between scRNA-seq and reporter strain data for stable expression (middle right). There was less overlap for cases with higher expression in L1 than L4 (rightmost). One-sided Fisher's Exact Tests were used to test for significant overlap. B-F) Validation of individual DEGs between L1 and L4. B) Dotplot denoting a subset of DEGs between the L1 and L4 in specific neurons compared to fluorescent reporter strains. The color scale reflects the log2 fold change from scRNA-seq data (red, positive values = higher in L4; blue, negative values = higher in L1). The shape of the point reflects whether the fluorescent reporter was consistent with scRNA-seq data. Squares indicate the reporter showed the same differential expression as scRNA-seq. Circles indicate the reporter did not show differential expression. Some of these genes showed differential expression by scRNA-Seq in additional neuron classes which were not tested for validation. C) Left: Micrographs of an endogenous lep-5 GFP reporter in CAN in L1 and L4. The CAN cell body was labeled with an endogenous swt-3::RFP marker. Right: Quantification of lep-5 expression. D) Left: Confocal micrograph of an endogenous ins-30 GFP reporter in CAN in L1 vs L4. The CAN cell body was labeled with a pks-1::TagRFP transgene. Right: Quantification of ins-30 reporter expression between L1 and L4. Fisher's Exact Test. E) Left: Micrographs of srlf-1 expression in HSN (labeled by cat-1 reporter fosmid in L1 and L4. Right: Quantification of decreased srlf-1 expression in HSN from L1 to L4. Fisher's Exact Test. F) Left: Micrographs of an endogenous flp-27::GFP reporter in L1 (top) and L4 (bottom). Right: Quantification of flp-27::GFP intensity in ALM, BDU and PLM. ALM and PLM showed significantly decreased flp-27 reporter expression from L1 to L4, whereas BDU showed no change. Unpaired t-tests. G-H) UMAPs as in Figure 1E showing lack of expression of the pan-neuronal genes egl-21 (G) and snt-1 (H) in HSN and VC. * = p-value < 0.05, ** p-value < 0.01, *** p-value < 0.001, **** p-value < 0.0001. Scale bars in C, E = 20 μm, in D, F = 10 μm.

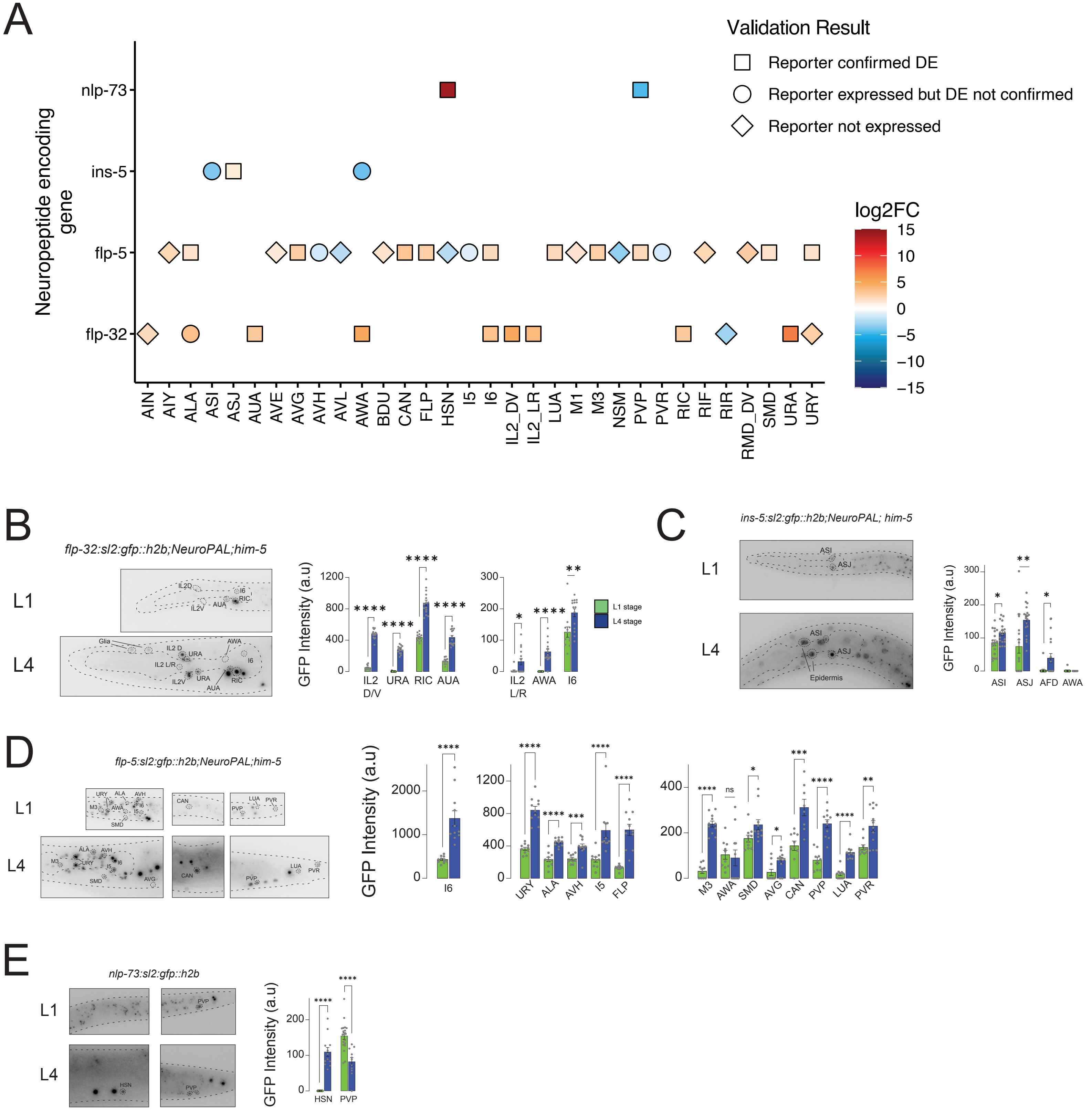
A
Validation Result
Reporter confirmed DE
Reporter expressed but DE not confirmed
Reporter not expressed
log2FC
15
10
5
0
−5
−10
−15
Neuropeptide encoding gene
nlp-73
ins-5
flp-5
flp-32
AIN AIY ALA ASI ASJ AUA AVE AVG AVH AVL AWA BDU CAN FLP HSN I5 I6 IL2_DV IL2_LR LUA M1 M3 NSM PVP PVR RIC RIF RIR RMD_DV SMD URA URY
B
flp-32:sl2:gfp::h2b;NeuroPAL;him-5
L1
L4
GFP Intensity (a.u)
IL2 D/V
URA
RIC
AUA
IL2 L/R
AWA
I6
L1 stage
L4 stage
C
ins-5:sl2:gfp::h2b;NeuroPAL; him-5
L1
L4
Epidermis
GFP Intensity (a.u)
ASI
ASJ
AFD
AWA
D
flp-5:sl2:gfp::h2b;NeuroPAL;him-5
L1
L4
GFP Intensity (a.u)
I6
URY
ALA
AVH
I5
FLP
M3
AWA
SMD
AVG
CAN
PVP
LUA
PVR
ns
E
nlp-73:sl2:gfp::h2b
L1
L4
GFP Intensity (a.u)
HSN
PVP

**Figure S9. Validation of differentially expressed neuropeptides, related to Figure 5.** A) Dotplot showing validation results for four endogenous neuropeptide reporters tested for differential expression between L1 and L4 in embryonically derived neurons (x- axis). All neurons with differential expression from scRNA-seq for these genes are shown. Color scale represents log2 fold change values from scRNA-seq data. Positive values (orange - red) = higher in L4, negative values (blue) = higher in L1. Shape represents the result of fluorescent reporter validation: squares indicate the reporter expression was consistent with scRNA-seq. Circles indicate the reporter was expressed in the given neuron but did not show differential expression between L1 and L4. Diamonds indicate the reporter was not detected in the neuron in either L1 or L4. B-F) Micrographs and quantification of reporter validation. Neurons with stable expression are not labeled in the micrographs for clarity. Postembyronically-derived neurons were not scored for validation due to lack of NeuroPAL coloring at L1. B) Left: Micrographs of flp-32 reporter expression in the head at L1 (top) and L4 (bottom). Right: quantification of GFP intensity in seven neuron classes. C) Left: Micrographs of ins-5 reporter expression in the head at L1 (top) and L4 (bottom). Right: quantification of GFP. The reporter showed a significant increase in AFD, the change in scRNA-seq data in AFD was not significant. D) Left: Micrographs of flp-5 reporter expression in the head, midbody and tail at L1 (top) and L4 (bottom). Right: quantification of GFP intensity. E) Left: Micrographs of nlp-73 reporter expression in the midbody and tail at L1 (top) and L4 (bottom). Right: quantification of GFP intensity. F) Left: Micrographs of nlp-64 reporter in L1 (top) and L4 (bottom). Right: quantification of GFP intensity in AVE and CAN. Mann-Whitney U tests were used for significance testing. * = p-value < 0.05, ** p-value < 0.01, *** p-value < 0.001, **** p-value < 0.0001.

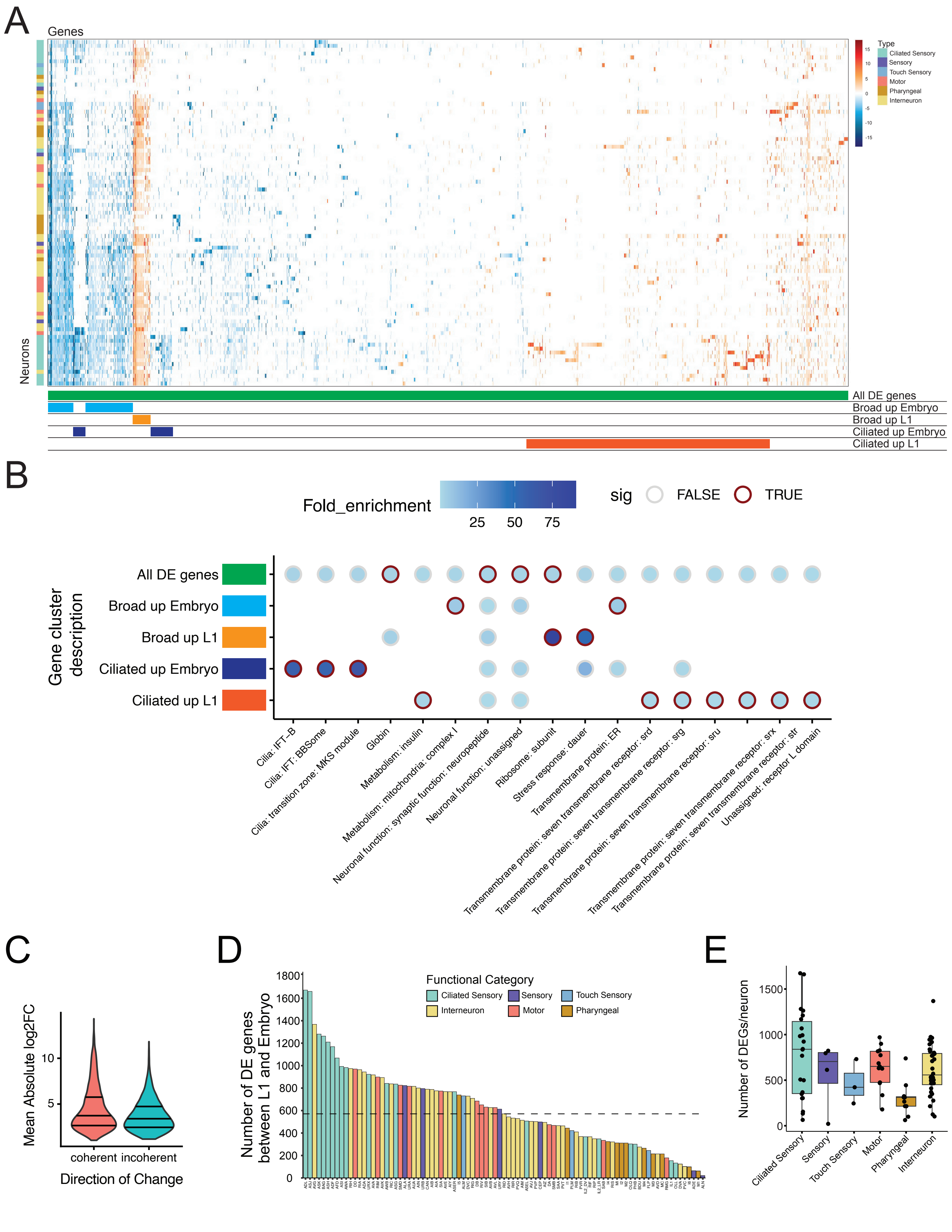

A
Genes
Neurons
Type
Ciliated Sensory
Sensory
Touch Sensory
Motor
Pharyngeal
Interneuron
All DE genes
Broad up Embryo
Broad up L1
Ciliated up Embryo
Ciliated up L1
B
Fold_enrichment
25
50
75
sig
FALSE
TRUE
Gene cluster description
All DE genes
Broad up Embryo
Broad up L1
Ciliated up Embryo
Ciliated up L1
Cilia: IFT-B
Cilia: IFT: BBSome
Cilia: transition zone: MKS module
Globin
Metabolism: insulin
Metabolism: mitochondria: complex I
Neuronal function: synaptic function: neuropeptide
Neuronal function: unassigned
Ribosome: subunit
Stress response: dauer
Transmembrane protein: ER
Transmembrane protein: seven transmembrane receptor: srd
Transmembrane protein: seven transmembrane receptor: srg
Transmembrane protein: seven transmembrane receptor: sru
Transmembrane protein: seven transmembrane receptor: srx
Transmembrane protein: seven transmembrane receptor: str
Unassigned: receptor L domain
C
Mean Absolute log2FC
10
5
coherent
incoherent
Direction of Change
D
Number of DE genes between L1 and Embryo
Functional Category
Ciliated Sensory
Sensory
Touch Sensory
Interneuron
Motor
Pharyngeal
1800
1600
1400
1200
1000
800
600
400
200
0
E
Number of DEGs/neuron
1500
1000
500
0
Ciliated Sensory
Sensory
Touch Sensory
Motor
Pharyngeal
Interneuron

**Figure S10. Ciliated neurons share differential expression between the embryo and L1, related to Figure 5.** A) Heatmap as in Figure 5A showing genes (columns) differentially expressed across the nervous system (neurons on y-axis). Neuron categories are depicted by the color scale on the left. Gene sets used for gene set enrichment analysis are depicted by the color bars on the bottom and labeled on the right. B) Wormcat enrichment analysis for five gene sets that showed clustered differential expression (labeled on left, corresponding to labels in panel A. Wormcat gene annotations are shown on the x-axis. The color scale represents fold enrichment over background, and categories with red borders were significant (Fisher's Exact Test with Bonferroni-corrected p-value $< 0.05$). Ciliated neurons shared patterns of higher expression in the embryo of genes (dark blue bars) related to cilia formation. C) Violin plot showing the mean absolute log2FC values for "coherent" and "incoherent" genes. D) Bar graph showing the number of DEGs per neuron class in the embryo vs L1 comparison, colored by functional category. The horizontal dashed line marks the median of 571 DEGs/neuron. E) Box and jitterplot showing the number of DEGs per neuron class in the embryo vs L1 comparison grouped by functional category. Linear regression with pairwise comparisons of estimated marginal means and Tukey's method for multiple comparisons.

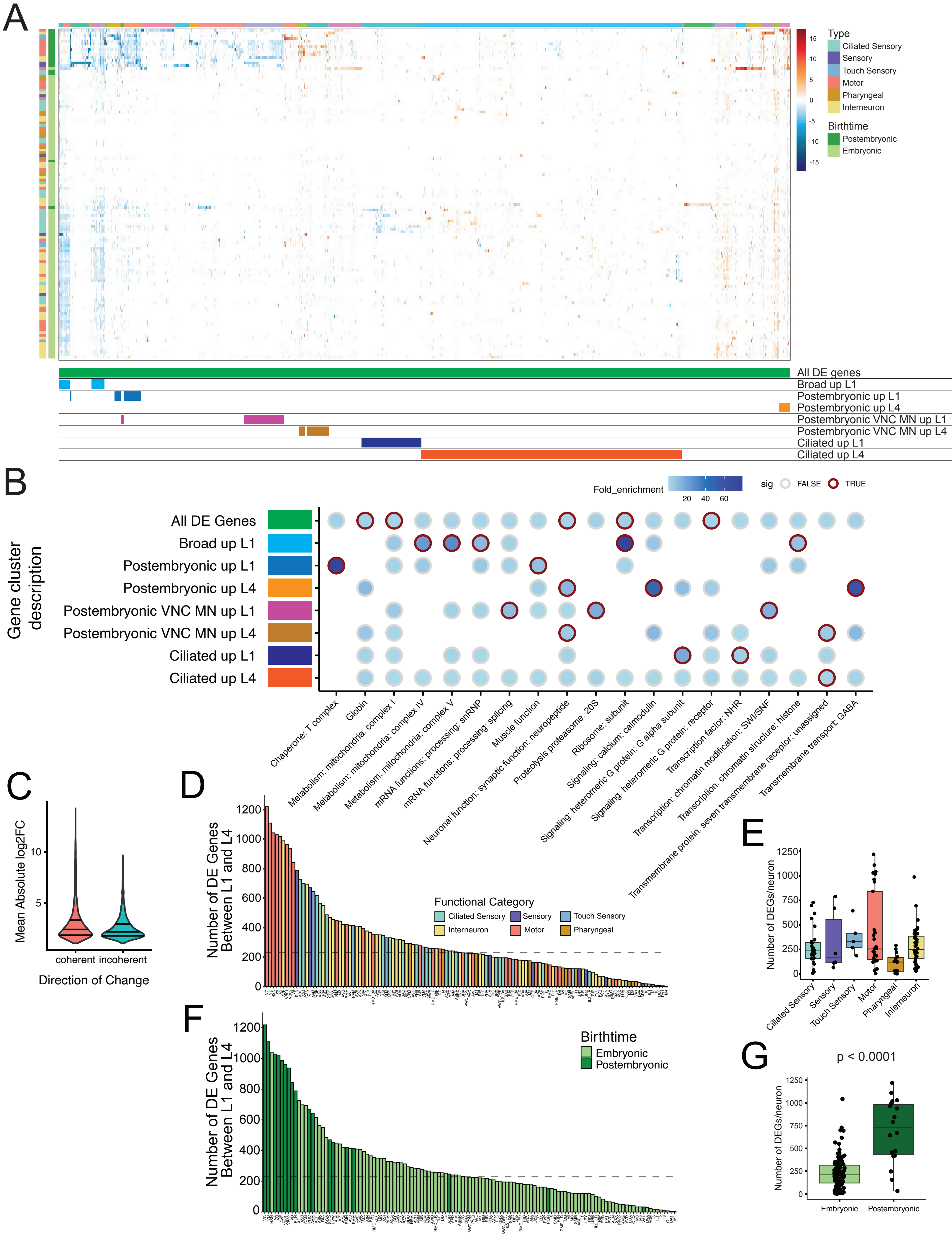

A
15
10
5
0
-5
-10
-15
Type
Ciliated Sensory
Sensory
Touch Sensory
Motor
Pharyngeal
Interneuron
Birthtime
Postembryonic
Embryonic
All DE genes
Broad up L1
Postembryonic up L1
Postembryonic up L4
Postembryonic VNC MN up L1
Postembryonic VNC MN up L4
Ciliated up L1
Ciliated up L4
B
Fold_enrichment
20
40
60
sig
FALSE
TRUE
Gene cluster description
All DE Genes
Broad up L1
Postembryonic up L1
Postembryonic up L4
Postembryonic VNC MN up L1
Postembryonic VNC MN up L4
Ciliated up L1
Ciliated up L4
Chaperone: T complex
Globin
Metabolism: mitochondria: complex I
Metabolism: mitochondria: complex IV
Metabolism: mitochondria: complex V
mRNA functions: processing: snRNP
mRNA functions: processing: splicing
Muscle function
Neuronal function: synaptic function: neuropeptide
Proteolysis proteasome: 20S
Ribosome: subunit
Signaling: calcium: calmodulin
Signaling: heteromeric G protein: G alpha subunit
Signaling: heteromeric G protein: receptor
Transcription factor: NHR
Transcription: chromatin modification: SWI/SNF
Transcription: chromatin structure: histone
Transmembrane protein: seven transmembrane receptor: unassigned
Transmembrane transport: GABA
C
Mean Absolute log2FC
10
5
coherent
incoherent
Direction of Change
D
Number of DE Genes Between L1 and L4
1200
1000
800
600
400
200
0
Functional Category
Ciliated Sensory
Sensory
Touch Sensory
Interneuron
Motor
Pharyngeal
E
Number of DEGs/neuron
1250
1000
750
500
250
0
Ciliated Sensory
Sensory
Touch Sensory
Motor
Pharyngeal
Interneuron
F
Number of DE Genes Between L1 and L4
1200
1000
800
600
400
200
0
Birthtime
Embryonic
Postembryonic
G
p < 0.0001
Number of DEGs/neuron
1250
1000
750
500
250
0
Embryonic
Postembryonic

**Figure S11. Characterization of differentially expressed genes between L1 and L4, related to Figure 5.** A) Heatmap as in Figure 5B showing differential gene expression between L1 and L4. Columns are genes, rows are neurons. Color scales on y-axis (left) denote functional groups and birthtimes. Hierarchical clustering grouped genes with similar patterns together. Color bars across columns indicate gene clusters. Colored bars below heatmap indicate gene sets that were used as input to Wormcat for gene set enrichment analysis, based on shared pattens of differential expression in multiple related neuron classes, as indicated by labels on the right. B) Gene set enrichment analysis for all DE genes (green, top row), or subsets of genes with shared differential expression across groups of neurons. Wormcat annotation categories are shown on the x-axis. Color scale reflects fold enrichment of genes in each category in the queried gene set over background (all genes detected by scRNA-seq). Dots with red borders were significantly enriched (Bonferroni-corrected p-value $< 0.05$). Significance was tested with Fisher's Exact Test followed by Bonferroni correction for multiple comparisons. C) Violin plot showing the mean absolute log2FC values for "coherent" and "incoherent" genes. D) Bar graph showing the number of DEGs/neuron class for the L1 vs L4 comparison, with neurons colored by functional category. The median of 227 DEGs/ neuron is depicted by the horizontal dashed line. E) Box and jitterplot of the number of DEGs/neuron grouped by functional category. F) Bar graph of DEGs/neuron for L1 vs L4 as in C but colored by birthtime (embryonic vs postembryonic). G) Box and jitterplot showing the number of DEGs/neuron grouped by birthtime. Postembryonically-derived neurons exhibited more DEGs/neuron than embryonically derived neurons. Linear regression with pairwise comparisons of estimated marginal means and Tukey's method for multiple comparisons.

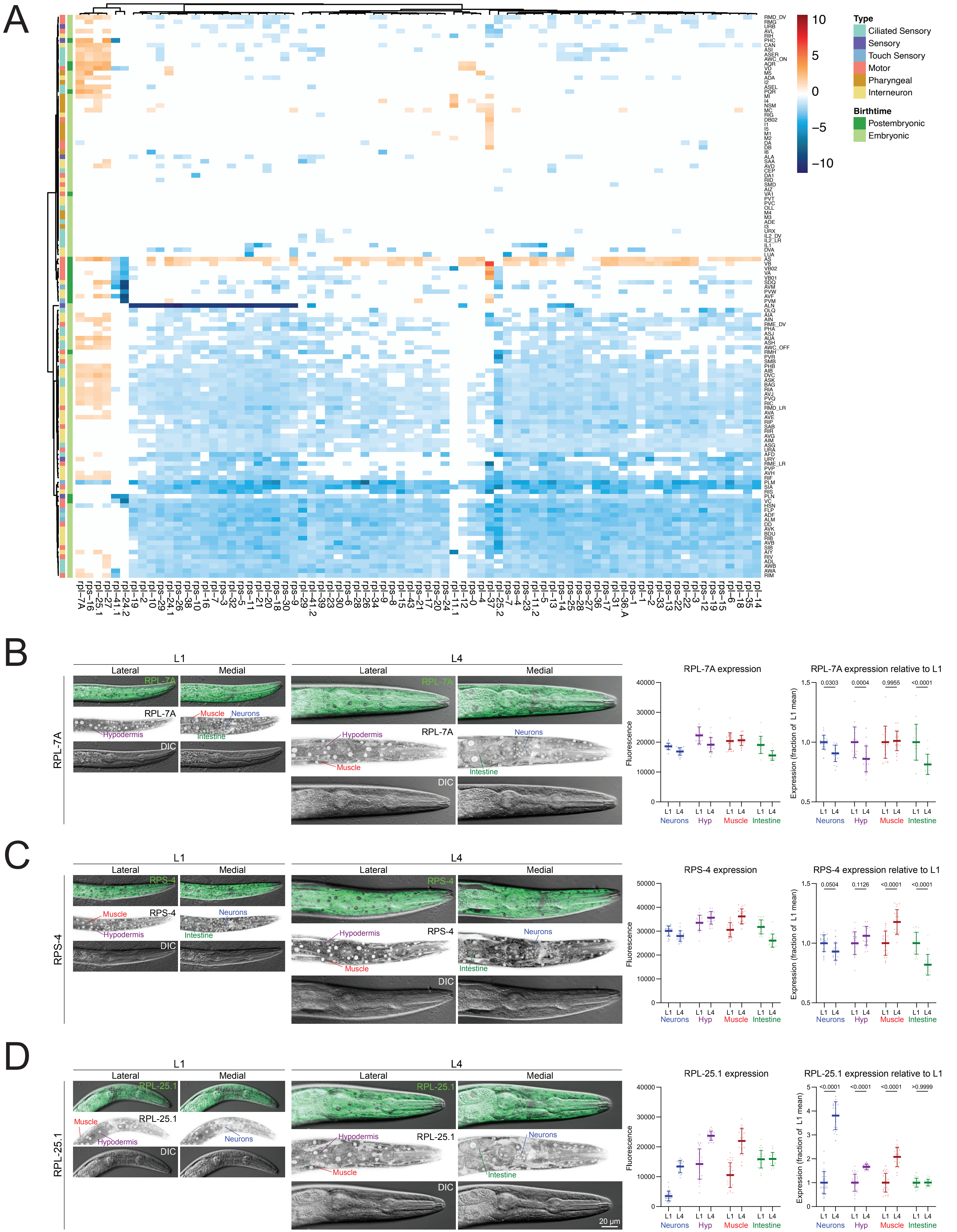
A
10
5
0
−5
−10
Type
Ciliated Sensory
Sensory
Touch Sensory
Motor
Pharyngeal
Interneuron
Birthtime
Postembryonic
Embryonic
RMD_DV RMG URB AVL RIH PHC CAN ASI ASER AWC_ON AQR VD M5 ADA I2 ASEL PQR MI I4 NSM MC RIG DB02 I1 I5 M1 M2 DA DB I6 ALA SAA AVD CEP DA1 RID SMD AIZ VA1 PVT PVC OLL M4 M3 ADE I3 URX IL2_DV IL2_LR IL1 DVA LUA AS VB VB02 VA VB01 SDQ AVM PVW AVF PVM ALN OLQ AIA AIN RME_DV PHA ASJ AUA ASH AWC_OFF RMH PVR SMB PHB AIB DVC ASK BAG RIA AVJ PVQ RIC RMD_LR AVA AVE RIP SAB RIR AVG AIM ASG URA AFD URY RME_LR PVP AVH RIF PLM SIA RIS PLN VC HSN FLP ADF ALM DD AVK BDU RIB AVB SIB AIY RIV ADL AWB AWA RIM
rpl−7A rps−16 rpl−25.1 rpl−27 rpl−41.1 rpl−24.2 rpl−19 rpl−2 rpl−10 rps−29 rpl−24.1 rps−26 rpl−38 rps−10 rpl−16 rpl−7 rps−3 rpl−32 rps−5 rps−11 rpl−21 rpl−20 rps−18 rpl−30 rps−9 rpl−29 rpl−41.2 rps−23 rpl−39 rps−30 rpl−23 rps−6 rpl−28 rpl−26 rpl−34 rpl−9 rps−8 rpl−15 rpl−43 rps−21 rpl−17 rps−20 rps−24 rpl−11.1 rpl−12 rps−0 rpl−4 rpl−37 rpl−25.2 rps−7 rps−4 rps−23 rpl−11.2 rpl−5 rpl−13 rps−14 rps−25 rps−28 rps−27 rpl−36 rps−17 rpl−31 rpl−36.A rps−1 rpl−1 rps−2 rpl−33 rps−13 rps−22 rpl−22 rpl−3 rps−12 rps−19 rps−15 rpl−6 rpl−18 rpl−35 rpl−14
B
RPL-7A
L1
L4
Lateral
Medial
RPL-7A
Muscle
Neurons
Hypodermis
Intestine
DIC
RPL-7A expression
Fluorescence
40000 30000 20000 10000 0
L1 L4 L1 L4 L1 L4 L1 L4
Neurons Hyp Muscle Intestine
RPL-7A expression relative to L1
Expression (fraction of L1 mean)
1.5 1.0 0.5
0.0303 0.0004 0.9955 <0.0001
C
RPS-4
RPS-4 expression
50000 40000 30000 20000 10000 0
RPS-4 expression relative to L1
0.0504 0.1126 <0.0001 <0.0001
D
RPL-25.1
RPL-25.1 expression
RPL-25.1 expression relative to L1
5 4 3 2 1 0
<0.0001 <0.0001 <0.0001 >0.9999
20 μm

**Figure S12. Genes encoding ribosomal proteins are broadly downregulated from L1 to L4, related to Figure 5.** A) Heatmap displaying the log2 fold change between L1 and L4 for ribosomal protein encoding genes (columns) with significant differential expression. Positive log2 fold change values (orange, red) reflect higher expression in L4, negative values (blue) reflect higher expression in the L1 stage. Color scales on y-axis (left) denote neuronal functional groups and birthtimes. B) Confocal micrographs showing RPL-7A::mEosEM (a green fluorescent protein) expression in hypodermis, muscle, intestine and neurons in the anterior regions of L1 (left) and L4 (center) larvae. Quantification (right) of fluorescence intensity shows a reduction in intensity among neurons ($p = 0.0303$), hypodermis and intestine between L1 and L4 stages. C) Confocal micrographs of RPS-4::mEosEM expression in anterior regions of L1 (left) and L4 (center). Quantification (right) of fluorescence intensity shows decreased expression in neurons, hypodermis and intestine, with increased expression in in muscle, from L1 to L4. D) Confocal micrographs showing RPL-25.1::mEosEM expression in hypodermis, muscle, intestine and neurons in the anterior regions of L1 (left) and L4 (center) larvae. Quantification (right) of fluorescence intensity shows significant increases in RPL-25.1 levels in neurons, muscle and hypodermis, but not intestine, between the L1 and L4 stages. 2-way ANOVA with Sidak's correction.

A

B

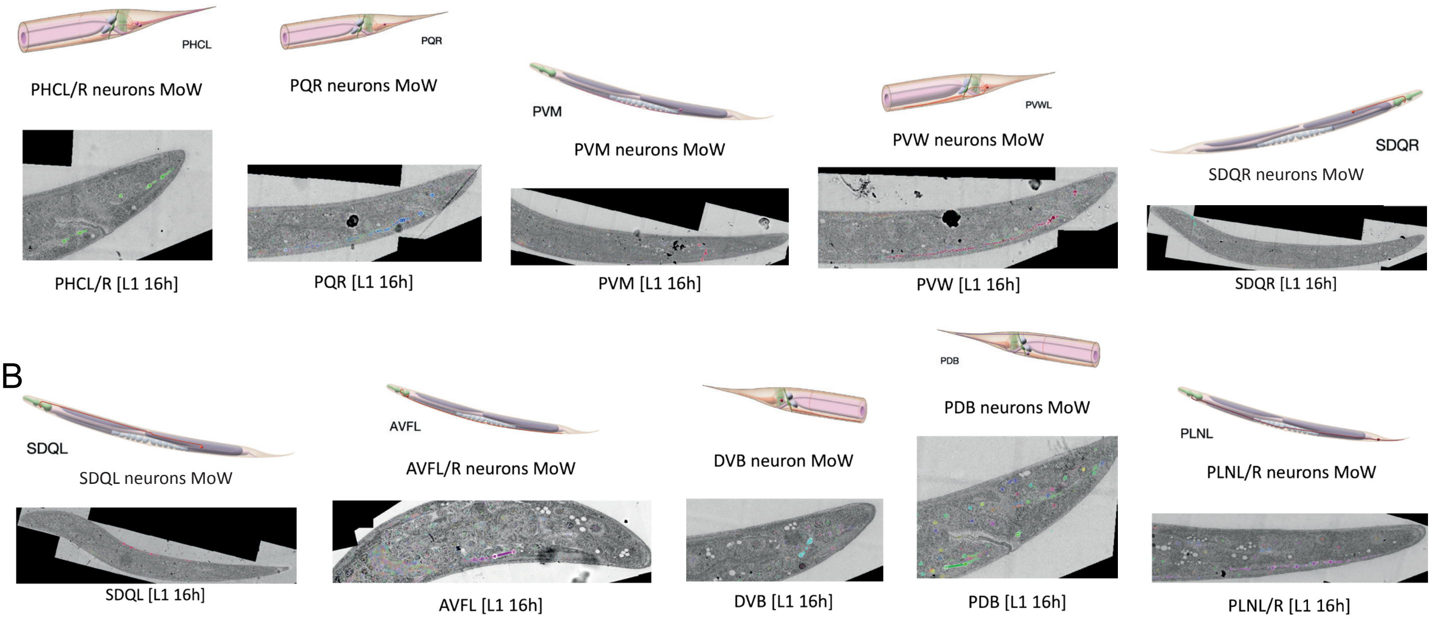


C

| | Embryo | L1 | L2 | L3 | L4 | Adult |
|---|---|---|---|---|---|---|
| Sex-shared neurons added | | AQR, PQR, AS, PVM, AVF, PVW, AVM, RMH, DVB, SDQ, PDB, VA, PHC, VB, PLN, VD | PDE, PVD, PVN, RMF | PDA (transdifferentiation) | | |
| Hermaphrodite specific neurons added/removed | CEMD/V Cell death | VC | | | HSN and VC terminal differentiation/ maduration | |

**Figure S13. Morphological description of neurons at the end of the first larval stage in C. elegans, related to Figure 6.** A-B) Morphological descriptions of neurons born during the first developmental stage (L1) are interpreted from annotated EM images of C. elegans 16h after hatching6. Shown are EM images of the last annotation for each neuron, representing the end of the neuron at this developmental stage. Neuron morphologies are compared to the fully matured adult stage descriptions published in the Mind of the Worm24 (MoW). Panel A features neurons that are fully matured at the end of the L1 larval stage. Panel B features neurons whose morphology is not fully matured at this stage. C) Table adapted from20. Neurons born and removed during each developmental stage, transdifferentiation events and differentiation/maturation of hermaphrodite neurons are listed.

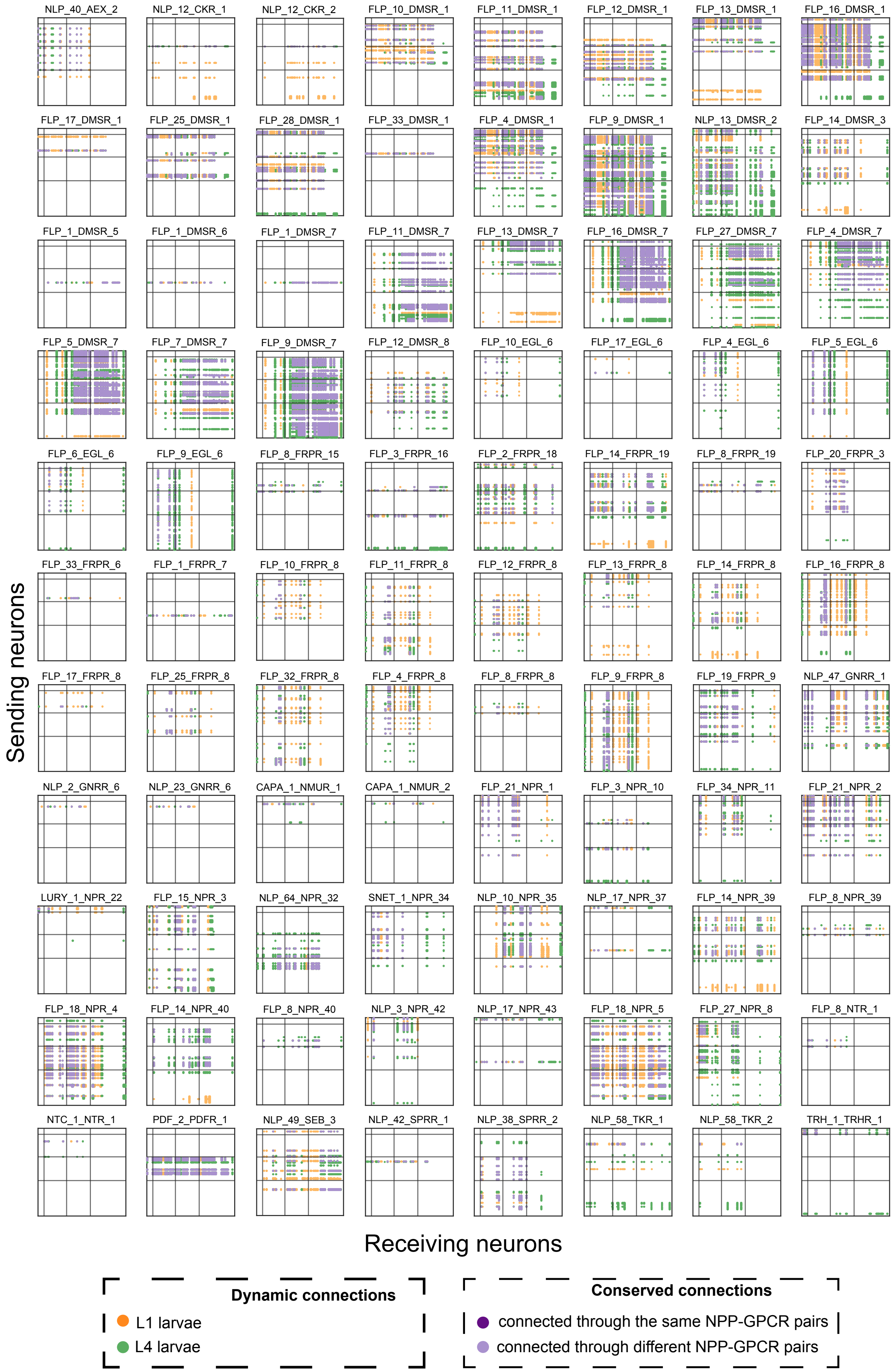

NLP_40_AEX_2
NLP_12_CKR_1
NLP_12_CKR_2
FLP_10_DMSR_1
FLP_11_DMSR_1
FLP_12_DMSR_1
FLP_13_DMSR_1
FLP_16_DMSR_1
FLP_17_DMSR_1
FLP_25_DMSR_1
FLP_28_DMSR_1
FLP_33_DMSR_1
FLP_4_DMSR_1
FLP_9_DMSR_1
NLP_13_DMSR_2
FLP_14_DMSR_3
FLP_1_DMSR_5
FLP_1_DMSR_6
FLP_1_DMSR_7
FLP_11_DMSR_7
FLP_13_DMSR_7
FLP_16_DMSR_7
FLP_27_DMSR_7
FLP_4_DMSR_7
FLP_5_DMSR_7
FLP_7_DMSR_7
FLP_9_DMSR_7
FLP_12_DMSR_8
FLP_10_EGL_6
FLP_17_EGL_6
FLP_4_EGL_6
FLP_5_EGL_6
FLP_6_EGL_6
FLP_9_EGL_6
FLP_8_FRPR_15
FLP_3_FRPR_16
FLP_2_FRPR_18
FLP_14_FRPR_19
FLP_8_FRPR_19
FLP_20_FRPR_3
FLP_33_FRPR_6
FLP_1_FRPR_7
FLP_10_FRPR_8
FLP_11_FRPR_8
FLP_12_FRPR_8
FLP_13_FRPR_8
FLP_14_FRPR_8
FLP_16_FRPR_8
FLP_17_FRPR_8
FLP_25_FRPR_8
FLP_32_FRPR_8
FLP_4_FRPR_8
FLP_8_FRPR_8
FLP_9_FRPR_8
FLP_19_FRPR_9
NLP_47_GNRR_1
NLP_2_GNRR_6
NLP_23_GNRR_6
CAPA_1_NMUR_1
CAPA_1_NMUR_2
FLP_21_NPR_1
FLP_3_NPR_10
FLP_34_NPR_11
FLP_21_NPR_2
LURY_1_NPR_22
FLP_15_NPR_3
NLP_64_NPR_32
SNET_1_NPR_34
NLP_10_NPR_35
NLP_17_NPR_37
FLP_14_NPR_39
FLP_8_NPR_39
FLP_18_NPR_4
FLP_14_NPR_40
FLP_8_NPR_40
NLP_3_NPR_42
NLP_17_NPR_43
FLP_18_NPR_5
FLP_27_NPR_8
FLP_8_NTR_1
NTC_1_NTR_1
PDF_2_PDFR_1
NLP_49_SEB_3
NLP_42_SPRR_1
NLP_38_SPRR_2
NLP_58_TKR_1
NLP_58_TKR_2
TRH_1_TRHR_1
Sending neurons
Receiving neurons
Dynamic connections
L1 larvae
L4 larvae
Conserved connections
connected through the same NPP-GPCR pairs
connected through different NPP-GPCR pairs

**Figure S14. Thresholded neuropeptidergic networks (mid-range) of the 88 NPP-GPCR pairs conserved between L1 and L4 larval stages, related to Figure 6.** The adjacency matrices display the developmental pattern of connections between sending neurons (y-axis) and receiving neurons (x-axis). Columns and rows are sorted by neuron class as in Figure 6 panel A. Core (conserved) connections and developmentally-dynamic connections are color-coded.

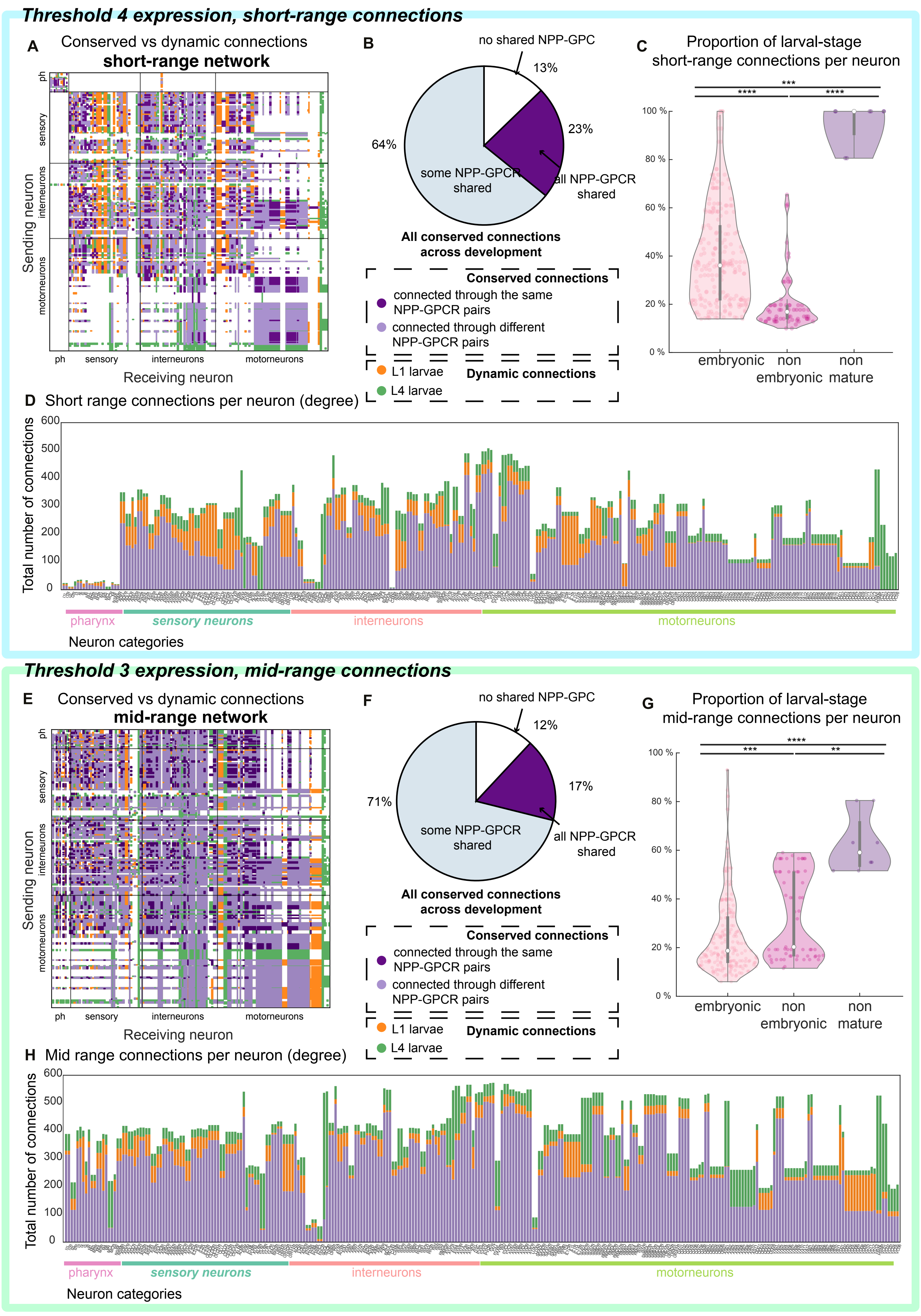

Threshold 4 expression, short-range connections
A
Conserved vs dynamic connections
short-range network
Sending neuron
ph
sensory
interneurons
motorneurons
Receiving neuron
B
no shared NPP-GPC
13%
23%
64%
some NPP-GPCR shared
all NPP-GPCR shared
All conserved connections across development
Conserved connections
connected through the same NPP-GPCR pairs
connected through different NPP-GPCR pairs
L1 larvae
L4 larvae
Dynamic connections
C
Proportion of larval-stage short-range connections per neuron
***
****
****
100 %
80 %
60 %
40 %
20 %
0 %
embryonic
non embryonic
non mature
D
Short range connections per neuron (degree)
Total number of connections
600
500
400
300
200
100
0
pharynx
sensory neurons
interneurons
motorneurons
Neuron categories
Threshold 3 expression, mid-range connections
E
Conserved vs dynamic connections
mid-range network
Sending neuron
ph
sensory
interneurons
motorneurons
Receiving neuron
F
no shared NPP-GPC
12%
17%
71%
some NPP-GPCR shared
all NPP-GPCR shared
All conserved connections across development
Conserved connections
connected through the same NPP-GPCR pairs
connected through different NPP-GPCR pairs
L1 larvae
L4 larvae
Dynamic connections
G
Proportion of larval-stage mid-range connections per neuron
****
***
**
100 %
80 %
60 %
40 %
20 %
0 %
embryonic
non embryonic
non mature
H
Mid range connections per neuron (degree)
Total number of connections
600
500
400
300
200
100
0
pharynx
sensory neurons
interneurons
motorneurons
Neuron categories

**Figure S15. Neuropeptide connectomes are robust across expression thresholds and physical distances, related to Figure 6.** A) Neuropeptidergic connectome (short-range) showing the developmental pattern of connections between sending neurons (y-axis) and receiving neurons (x-axis). The same structure and coloring scheme as Figure 6A were used. Threshold 4 expression data was used to construct this connectome. B) Pie chart showing the percentage distribution of the conserved connections between NPPs (Neuropeptide Precursor genes) and GPCRs (G-Protein Coupled Receptors) forming the core connectome. This chart corresponds to panel A connectome. Despite the increase in stringency of connections (short-range) we observe the same relative percentages as in the mid-range network (Figure 6). C) Proportion of developmentally dynamic neuropeptide connections (y-axis) per neuron for the connectome in panel A classified by embryonic development (colors and x-axis; "non-mature" refers to the HSN and VC neurons). Kruskal-Wallis test with Dunn's correction, ****$P < 0.0001$, ***$P < 0.001$, *$P < 0.025$. D) Total number of degrees (y-axis, short-range networks) in homologous neurons across development (x-axis) classified by functional categories for the network in panel A. E-H) Mid-range network characteristics with a relaxed expression threshold (Threshold 3). E) Neuropeptidergic connectome (mid-range) showing the developmental pattern of connections between sending neurons (y-axis) and receiving neurons (x-axis). The same structure and coloring scheme as panel A were used. Threshold 3 expression data was used to construct this connectome but, in this case, mid-range connections are present, as in Figure 6 panel A. F) Pie chart like panel B. This chart corresponds to panel E connectome. Despite the decrease in stringency of expression (threshold 3) we observe the same relative percentages as with threshold 4 (Figure 6). G) Proportion of developmentally dynamic neuropeptide connections (y-axis) per neuron for the connectome in panel E. Same as panel C. H) Total number of degrees (y-axis, mid-range networks) in homologous neurons across development (x-axis) classified by functional categories for the network in panel E.

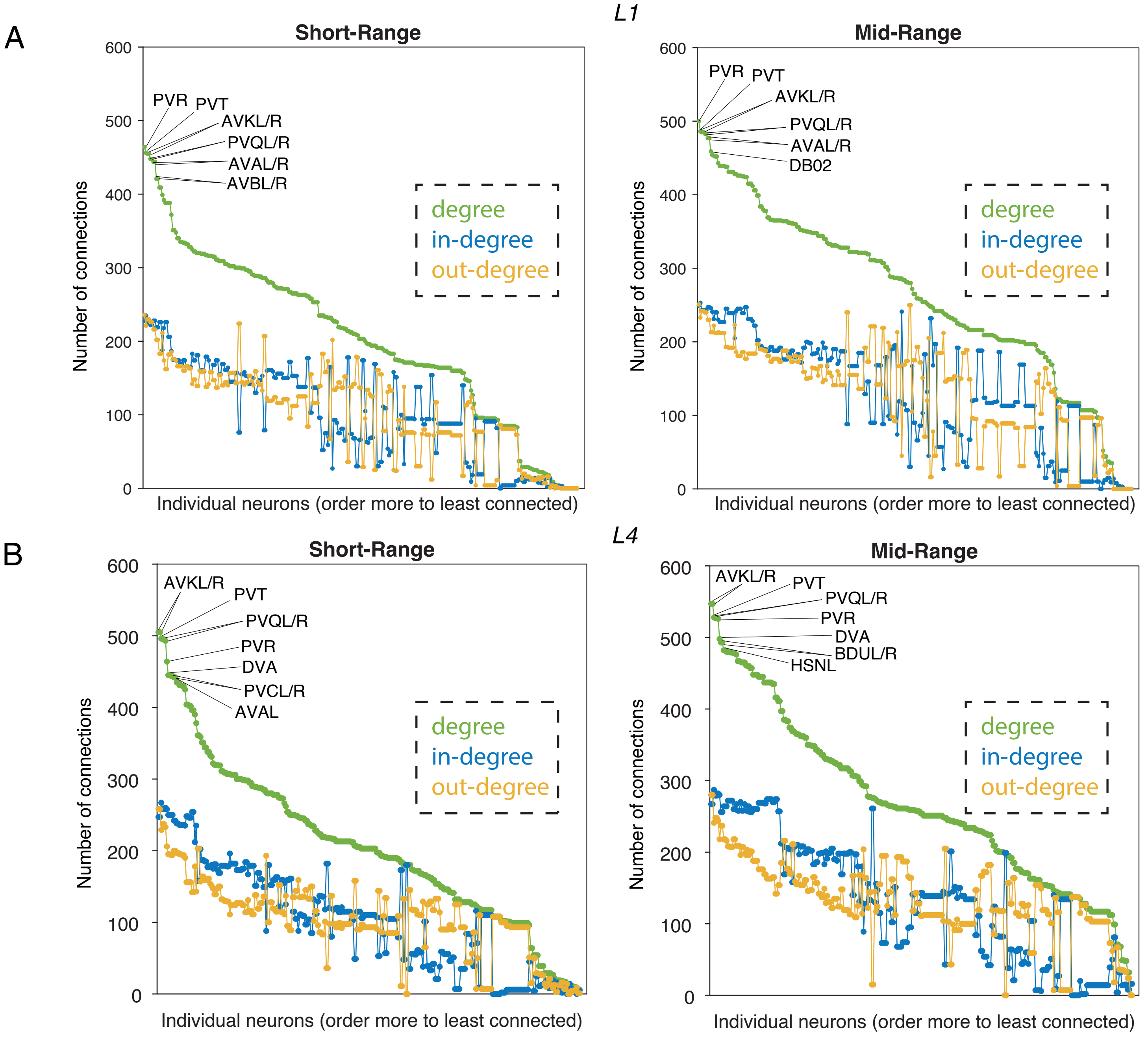


C

#### Convergent

| Neuron | Knorm | Kstd | L1 | L4 |
|---|---|---|---|---|
| PVR | 1 | 0 | 466 | 476 |
| PVT | 0.957739019 | 0.005914618 | 458 | 464 |
| PVQL | 0.938955347 | 0.004720072 | 450 | 463 |
| PVQR | 0.938955347 | 0.004720072 | 450 | 463 |
| AVKL | 0.916003105 | 0.032982389 | 457 | 445 |
| AVKR | 0.916003105 | 0.032982389 | 457 | 445 |
| AVAL | 0.869518765 | 0.034344818 | 446 | 433 |
| AVAR | 0.869518765 | 0.034344818 | 446 | 433 |
| DVA | 0.801926765 | 0.050487334 | 401 | 444 |
| AVBL | 0.797639485 | 0.020507453 | 423 | 419 |
| AVBR | 0.797639485 | 0.020507453 | 423 | 419 |
| AVEL | 0.728289654 | 0.023548703 | 390 | 415 |
| AVER | 0.728289654 | 0.023548703 | 390 | 415 |
| PVPL | 0.720591727 | 0.045928687 | 411 | 390 |
| PVPR | 0.720591727 | 0.045928687 | 411 | 390 |
| AVDL | 0.681992512 | 0.015567843 | 390 | 389 |
| AVDR | 0.681992512 | 0.015567843 | 390 | 389 |
| RMEV | 0.651209935 | 0.053231562 | 394 | 368 |
| RID | 0.642060086 | 0.001820876 | 374 | 382 |
| PVCL | 0.579399142 | 0.181177145 | 300 | 429 |

#### Divergent

| Neuron | Knorm | Kstd | L1 | L4 |
|---|---|---|---|---|
| HSNL | 0.166487079 | 0.507404484 | 86 | 430 |
| HSNR | 0.166487079 | 0.507404484 | 86 | 430 |
| AVM | 0.248365446 | 0.41351961 | 133 | 415 |
| SDQL | 0.004858004 | 0.394122762 | 4 | 272 |
| VC01 | 0 | 0.34151753 | 0 | 233 |
| VC02 | 0 | 0.34151753 | 0 | 233 |
| OLQDL | 0.166560131 | 0.280750642 | 304 | 126 |
| OLQDR | 0.166560131 | 0.280750642 | 304 | 126 |
| OLQVL | 0.166560131 | 0.280750642 | 304 | 126 |
| OLQVR | 0.166560131 | 0.280750642 | 304 | 126 |
| PVM | 0.004132043 | 0.264724352 | 5 | 187 |
| IL1DL | 0.118573646 | 0.254670793 | 265 | 104 |
| IL1DR | 0.118573646 | 0.254670793 | 265 | 104 |
| IL1L | 0.118573646 | 0.254670793 | 265 | 104 |
| IL1R | 0.118573646 | 0.254670793 | 265 | 104 |
| IL1VL | 0.118573646 | 0.254670793 | 265 | 104 |
| IL1VR | 0.118573646 | 0.254670793 | 265 | 104 |
| CEPDL | 0.104784951 | 0.251532688 | 255 | 96 |
| CEPDR | 0.104784951 | 0.251532688 | 255 | 96 |
| CEPVL | 0.104784951 | 0.251532688 | 255 | 96 |

**Figure S16. Neuropeptidergic hubs are largely conserved across age, related to Figure 6.** (A) Degree distributions of C. elegans L1 neural networks. In each case, degree (incoming plus outgoing connections) is shown in green, in-degree (incoming connections) in blue and out-degree (outgoing connections) in yellow. The 10 highest-degree hubs in each network are indicated. (B) Degree distributions of C. elegans L4 neural networks, displayed in the same way as panel A. (C) Top 20 neurons that show convergence (similar number of connections, Knorm) or divergence (different number of connections, Kstd) between L1 and L4 larval stages.

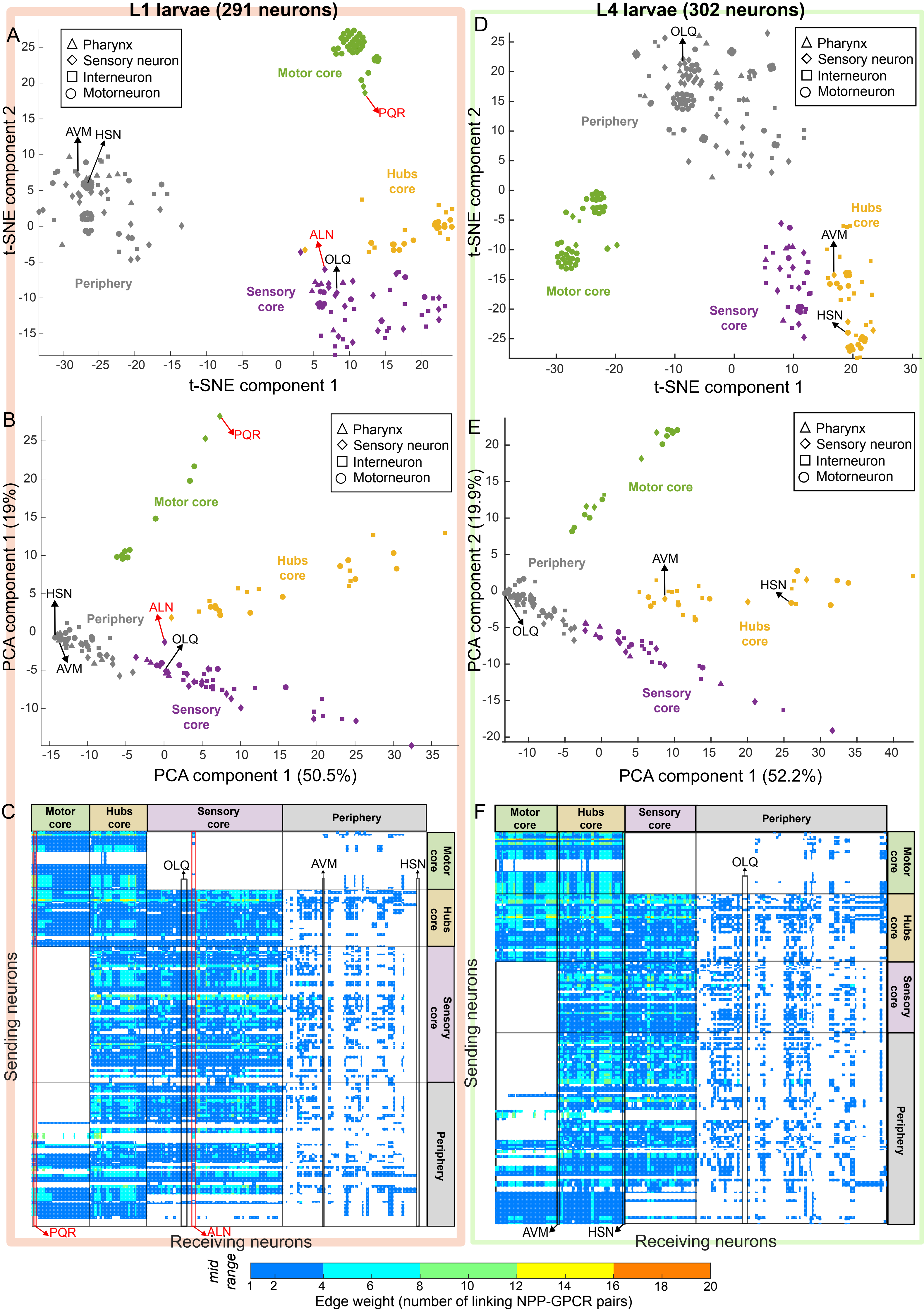
L1 larvae (291 neurons)
L4 larvae (302 neurons)
A
D
B
E
C
F
Pharynx
Sensory neuron
Interneuron
Motorneuron
Motor core
Hubs core
Sensory core
Periphery
PQR
ALN
OLQ
AVM
HSN
t-SNE component 1
t-SNE component 2
PCA component 1 (50.5%)
PCA component 1 (19%)
PCA component 1 (52.2%)
PCA component 2 (19.9%)
Sending neurons
Receiving neurons
mid range
Edge weight (number of linking NPP-GPCR pairs)

**Figure S17. Mesoscale structure of neuropeptide networks is largely conserved across development, related to Figure 6.** A-C) L1 larval data, D-F) L4 larval data. The L1 data shown is from 291 neurons present at the L1 stage, the L4 data includes the 302 neurons present at the L4 stage. (A and D) 2D t-SNE plot showing the grouping of neurons based on the incoming connection profiles for each neuron. Visually distinct clusters of neurons are apparent based on local proximity in t-SNE space and are indicated by different colors. Clusters are mostly distinct; any ambiguity in cluster assignment was resolved by comparing the relevant neuron's location in the clusters on the PCA plots in panels B and E. Marker shapes indicate neuron types. (B and E) 2D PCA plot showing the grouping of neurons based on the incoming connection profiles for each neuron. Visually distinct clusters of neurons are apparent based on local proximity in PCA space. Points are colored as in panels A and D. Marker shapes indicate neuron types. Axis labels indicate the percentage variance explained by each PCA component plotted. (C and F) Weighted adjacency matrices for L1 (C) and L4 (F), with rows and columns rearranged by the groupings derived from t-SNE/PCA (as illustrated by colors in panels A, B, D, E). The original weighted adjacency matrices for L1 and L4 are shown in Figure 6C. Weights (colors) represent the count of neuropeptide-receptor pairs mediating each connection. Neurons highlighted in black are those whose group assignment differs between L1 and L4, shown consistently across all panels. Neurons highlighted in red are those whose group assignment diverges between block modelling and t-SNE/PCA approaches.

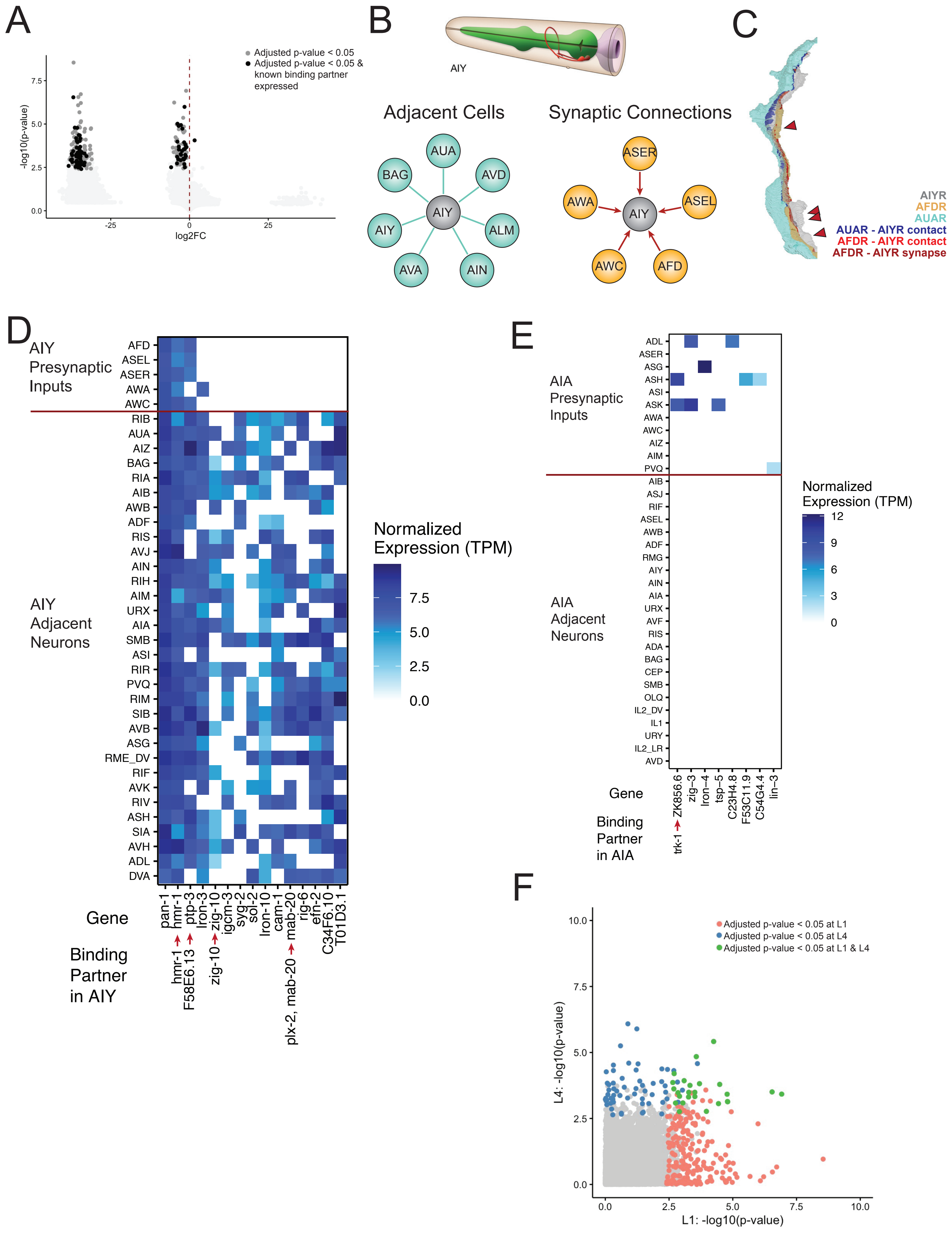
A
Adjusted p-value < 0.05
Adjusted p-value < 0.05 & known binding partner expressed
-log10(p-value)
log2FC
B
AIY
Adjacent Cells
AUA
BAG
AVD
AIY
ALM
AVA
AIN
Synaptic Connections
ASER
AWA
ASEL
AWC
AFD
C
AIYR
AFDR
AUAR
AUAR - AIYR contact
AFDR - AIYR contact
AFDR - AIYR synapse
D
AIY Presynaptic Inputs
AFD
ASEL
ASER
AWA
AWC
AIY Adjacent Neurons
RIB
AUA
AIZ
BAG
RIA
AIB
AWB
ADF
RIS
AVJ
AIN
RIH
AIM
URX
AIA
SMB
ASI
RIR
PVQ
RIM
SIB
AVB
ASG
RME_DV
RIF
AVK
RIV
ASH
SIA
AVH
ADL
DVA
Gene
pan-1
hmr-1
ptp-3
lron-3
zig-10
igcm-3
syg-2
sol-2
lron-10
cam-1
mab-20
rig-6
efn-2
C34F6.10
T01D3.1
Binding Partner in AIY
hmr-1
F58E6.13
zig-10
plx-2, mab-20
Normalized Expression (TPM)
7.5
5.0
2.5
0.0
E
AIA Presynaptic Inputs
ADL
ASER
ASG
ASH
ASI
ASK
AWA
AWC
AIZ
AIM
PVQ
AIA Adjacent Neurons
AIB
ASJ
RIF
ASEL
AWB
ADF
RMG
AIY
AIN
AIA
URX
AVF
RIS
ADA
BAG
CEP
SMB
OLQ
IL2_DV
IL1
URY
IL2_LR
AVD
Gene
ZK856.6
zig-3
lron-4
tsp-5
C23H4.8
F53C11.9
C54G4.4
lin-3
Binding Partner in AIA
trk-1
Normalized Expression (TPM)
12
9
6
3
0
F
Adjusted p-value < 0.05 at L1
Adjusted p-value < 0.05 at L4
Adjusted p-value < 0.05 at L1 & L4
L4: -log10(p-value)
L1: -log10(p-value)

**Figure S18. Connectivity analysis reveals genes enriched based on membrane contact or presynaptic input to nerve ring neurons, related to Figure 7.** A) Volcano plot showing enrichment of CAMs based on synaptic connectivity. Each dot represents a comparison of one gene tested in the synaptically-connected vs adjacent only neurons for a given neuron. Negative log fold change values represent enrichment in non-synaptic adjacent cells, whereas positive log fold change represent enrichment in synaptically-connected cells. Light gray dots represent instances with Benjamini-Hochberg adjusted p-values > 0.05. Dark gray dots represent cases with adjusted p-values < 0.05. Black dots represent cases with adjusted p-values < 0.05 and in which the tested gene has a known binding partner expressed in the neuron of interest. B) Top: Schematic representation of the AIY neurons, from WormAtlas. Bottom: Adjacent neurons with only membrane contact (left) or synaptic inputs (right) with AIY. Only a subset of adjacent neurons with membrane contact are shown. C) 3D reconstruction from NeuroScan of late L1 nerve ring featuring AIYR, AUAR and AFDR. Both AUAR (light blue) and AFDR (orange) contact AIYR (dark blue and red, respectively), but only AFDR is presynaptic to AIYR (red triangle). D) Heatmap showing the log fold change of 15 CAMs enriched in AIY + non-synaptic adjacent cells (below red line) compared to AIY + presynaptic inputs (above red line). Genes are sorted by log fold change (lowest fold change on left). Genes with known binding partners expressed in AIY are denoted with red arrows, and the respective binding partners are listed below. E) Heatmap showing several genes with large fold changes between AIA synaptically connected neurons and membrane adjacent-only neurons. These eight genes are expressed in only a subset of AIA presynaptic inputs and therefore have adjusted p-values > 0.05. They may, however, still regulate specific individual connections. F) Scatterplot showing -log10 transformed p-values for each gene-neuron combination in L1 (x-axis) and in L4 (y-axis). Genes only detected at one age are not shown. Significant cases (Benjamini-Hochberg adjusted p-value <0.05) of enrichment in L1 (pink), cases significantly enriched at L4 (blue), and cases enriched at both L1 and L4 (green). The p-value adjustments were made for multiple genes within each neuron.
